# An intrinsically disordered region controls viperin (SAND) antiviral synthesis via a clamp-and-latch mechanism

**DOI:** 10.64898/2026.09.10.750538

**Authors:** Mengdi Wu, Denise Poire, Hannah Florance, Simone Ciofi-Baffoni, William S. James, Maria Andrea Mroginski, Kourosh H. Ebrahimi

**Affiliations:** Institute of Pharmaceutical Science, Faculty of Life Sciences and Medicine, King’s College London; Franklin-Wilkins Building, 150 Stamford Street, London, SE1 9NH, UK; Department of Chemistry, Technical University of Berlin, Chemiegebäude, Str. des 17. Juni 115, 10623 Berlin, Germany; Agilent Technologies, Cheadle, UK; Magnetic Resonance Center and Department of Chemistry, University of Florence, Via Sacconi, 6, 50019 Sesto Fiorentino, Florence, Italy; Sir William Dunn School of Pathology, University of Oxford, South Parks Road, Oxford, OX1 3RE, UK

## Abstract

Inter-individual susceptibility to viral infections is a public health challenge. Understanding the structure and function of antiviral restriction-factor enzymes is necessary to link genetic variants to viral susceptibility. Viperin (SAND) is a sentinel antiviral restriction factor with broad-spectrum activity believed to preferentially select cytidine triphosphate (CTP) based on the structure of the active site. Here, we studied fungal, rat, and human SANDs. We found that a C-terminal intrinsically disordered region (IDR) ensemble, not the active site, is the key structural element enabling the enzyme to select multiple ribonucleoside triphosphates (rNTPs). We demonstrate that the IDR ensemble functions via a clamp-and-latch mechanism that depends on the IDR-adjacent folded domain (context). Consequently, human and rat enzymes with different contexts have different catalytic outputs. These findings revise our understanding of viperin catalytic function and identify an IDR-dependent element in the evolution of antiviral defense.

## Introduction

The cellular innate immune antiviral response, the first line of defense against viral infections, comprises several antiviral restriction factors, such as SAND (viperin)^1–3^, RNase L^4^, and ADAR1^5^. Many of these enzymes have intrinsically disordered regions (IDRs), whose structures and antiviral functions have remained largely unclear. Mechanistic understanding of IDR function in antiviral restriction factors could help identify genetic variants leading to inter-individual viral susceptibility. However, understanding IDR functions remains challenging. IDRs are ubiquitous, comprising ∼20% of bacterial and ∼70% of human proteomes^6–9^. Lacking a stable 3D structure (dark proteome) ^10^, IDRs are invisible to X-ray crystallography and unresolved by Cryo-EM. They are best described as an ensemble, that is, a collection of dynamically shifting conformations ^6–9^. Their function is altered by their adjacent folded protein (context), and their dynamics can modulate context structure-function. This complex dynamic IDR sequence-context-ensemble function poses a challenge in dissecting their mechanistic rules in various biological processes^6,11–19^.

A conserved feature of the cellular defense system against viral infections, from bacteria to humans,^1,20^ is the widely studied antiviral radical S-adenosylmethionine (SAM)-dependent nucleotide dehydratase (SAND), the UniProt-recommended name for viperin (virus-inhibitory protein endoplasmic reticulum-associated interferon-inducible) or RSAD2.^3,21–23^ In humans, the enzyme has broad-spectrum antiviral activity, restricting replication of several viruses, including Zika virus, influenza A virus (IAV), HIV-1, Hepatitis C virus (HCV), and West Nile Virus (WNV), among others.^2,24–29^ Vertebrate SANDs (viperin) are composed of three domains: an N-terminal amphipathic domain, a central radical-SAM domain, and a C-terminal domain.^26,27^ The N-terminal amphipathic domain localizes the protein to the cytosolic face of the endoplasmic reticulum (ER) membrane or lipid droplets.^30,31^ ER membrane localization is essential for efficient electron transfer from the putative electron-transfer partner, the membrane-anchored cytochrome b5-reductase 3 (CYB5R3), and catalysis.^22^ The enzymatic transformation is carried out by the central radical-SAM domain with the CxxxCxxC motif coordinating a [4Fe-4S] cluster.^2,32^ The cluster reductively cleaves SAM to generate a 5’-deoxyadenosyl radical, which abstracts the H4’ hydrogen atom of a ribonucleoside triphosphate (rNTP) to initiate catalysis (Fig. 1a). Based on studies with rat SAND (rSAND), it is widely accepted that vertebrates’ SANDs with identical catalytic pockets catalyze the transformation of CTP as the primary substrate to its antiviral nucleotide analog 3’-deoxy-3’,4’-didehydro-CTP (ddhCTP) (Fig. 1a).^3,33,34^ The catalysis occurs through a diol dehydration mechanism^35^ and has a low turnover rate (2-4 hr^-1^) similar to many other radical-SAM enzymes.^27,36^ Fungal and microbial SANDs catalyze similar transformations using other rNTPs as substrates.^33,35,37^ The ddhNTP analogs lack a 3′-hydroxyl group and can chain-terminate phage and viral replication,^34,37,38^ and ddhCTP is proposed to regulate immunometabolism.^39,40^

**Figure 1.**
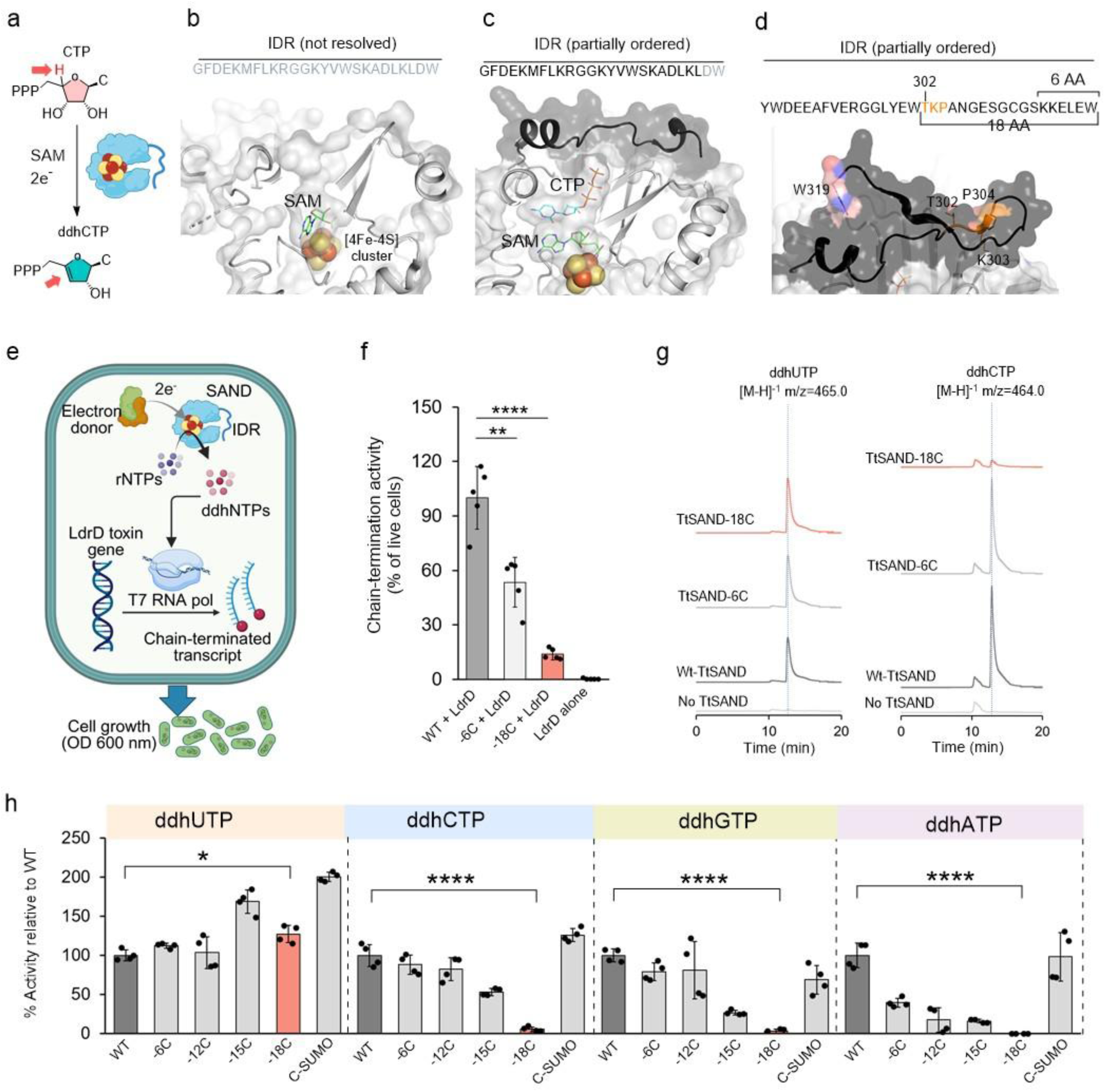
The IDR tripeptide TKP confers substrate promiscuity in fungal SAND. **(a)** The biocatalytic synthesis of ddhCTP by vertebrate SANDs (viperin). The catalysis involves SAM and two electrons, through a 5’-dA• radical that abstracts H4’ of CTP (red arrow) to generate ddhCTP. This nucleotide analog lacks the 3’-OH group and has a double bond between C3’ and C4’ (Red arrow), therefore acting as a chain terminator for viral RNA polymerase. **(b)** The X-ray crystal structure of rat SAND (rat viperin) in the absence (PDB code 5VSL) or **(c)** in the presence of CTP (PDB code 6Q2P). The grey residues in the sequence are not well resolved in the X-ray structure. **(d)** The C-terminus of fungal TtSAND has an IDR predicted by AlphaFold (Black). 6 and 18 amino acid (AA) residues of the C-terminal IDR are highlighted. **(e)** VITAS assay. Enzymatically produced ddhNTP is incorporated into the LdrD toxin transcript by phage T7 RNA polymerase. Consequently, transcription is chain-terminated, and a functional LdrD is not produced, and *E. coli* cells will grow. **(f)** Removal of 18 residues from the C-terminal IDR reduced chain-termination activity in cells by >80%. Data are the average of five biologically independent measurements ± standard deviation. **(g)** Ion chromatograms of ddhUTP and ddhCTP recorded in negative mode at the specified m/z ratio are shown as an example. **(h)** Loss of the TKP tripeptide in the -18C variant made the enzyme specific to ddhUTP production. The amount of ddhNTP was quantified for each variant, and the results are plotted as the percentage of activity of wild-type (WT) TtSAND. Data are the average of four independent measurements using two different protein batches ± standard deviation. One-way ANOVA was used to calculate p-values. (*p value < 0.05; **p value < 0.01; ****p value < 0.0001).

While the functions of the N-terminal domain and catalytic core are well defined, the C-terminal domain remains structurally and functionally uncharacterized. Mutations within this domain are associated with a loss of antiviral function via an unknown mechanism.^41–43^ Here, we found that this domain, from bacteria to humans, contains an IDR that calibrates antiviral outputs. We show that the IDR has two motifs: a highly conserved RxxxY motif that acts as a clamp and a variable tripeptide that acts as a latch to control the IDR ensemble. The tripeptide and its context, a hotspot for mutations, modulate IDR ensemble gating of substrates. Variations in the latch context enable human and rat enzymes to generate different profiles of ddhNTP antivirals. These findings revise our understanding of SAND (viperin) catalytic function and antiviral activity.^3,33,34^

## Results and discussion

### C-terminus has an IDR required for chain-termination activity

The structure of the C-terminus domain of rSAND is unresolved in the absence of substrate (Fig. 1b) and is only partially ordered in the presence of CTP substrate (Fig. 1c).^44,45^ These observations suggest this domain has an intrinsically disordered region (IDR). Residue-level IDR prediction using the Novopro tool confirmed the presence of a C-terminal IDR of 20-30 amino acid residues across SANDs from all kingdoms of life (Supplementary Fig. 1). The C-terminal tryptophan in human SAND (hSAND) (RSAD2 or viperin) is suggested to play a role in [4Fe-4S] cluster delivery to the enzyme.^46,47^ A tryptophan residue is present in the C-terminal IDR of some bacterial and fungal SANDs, such as the thermophilic fungus *Thielavia terrestris* SAND (TtSAND), with more than 40% identity to human SAND (viperin)^48^ (Fig. 1d and Supplementary Fig. 2). Therefore, we investigated the role of the IDR in [4Fe-4S] cluster delivery to fungal SANDs. To this end, we first created two C-terminal truncated variants of the enzyme by deleting 6 or 18 amino acid residues of the IDR, TtSAND-6C and -18C, respectively (Fig. 1d). The -6C deletion removes the terminal tryptophan, while the -18C deletion restores it.

Truncation did not affect the enzymes’ [4Fe-4S] cluster UV-visible absorbance characteristics or their [4Fe-4S] cluster content (Supplementary Fig. 3). Next, we analyzed in-cell chain-termination activity using a live-dead assay named VITAS (viral polymerase inhibition toxin-associated selection) (Fig. 1e).^22,49^ In VITAS, ectopic expression of a SAND generates ddhNTP using the cellular rNTP pool. The resulting ddhNTP chain-terminates phage T7 RNA polymerase activity, blocking T7 promoter-dependent transcription of toxin LdrD and rescuing cell growth. Thus, chain-termination activity of SAND variants can be described as a percentage of cell growth relative to the live control, in which the wild-type enzyme is expressed (Supplementary Methods).^22^ While the TtSAND-6C variant retained 50-60% of wild-type activity in rescuing cell growth, the TtSAND-18C variant lost > 80% of its activity (Fig. 1f). We conclude that specific amino acid residues in the IDR region of TtSAND regulate its chain-termination activity.

### An IDR tripeptide confers substrate selection in fungi

Since truncation did not affect the [4Fe-4S] cluster content but did affect chain termination, we hypothesized that the C-terminal IDR in TtSAND modulates enzymatic activity. Therefore, we analyzed ddhNTP formation by purified enzymes. We first created two new truncated variants by removing 12 or 15 residues (the -12C and -15C variants, respectively). Additionally, a variant in which SUMO was added to the C-terminus was created as a control to analyze the difference between truncation of the C-terminus and its extension. As before, mutations did not alter the UV-visible absorbance spectrum of the enzyme [4Fe-4S] cluster and its content (Supplementary Fig. 4).

Subsequently, we used liquid chromatography-mass spectrometry (LC-MS) to measure the formation of ddhUTP, ddhCTP, ddhGTP, or ddhATP from the corresponding rNTP substrates (Fig. 1g and Supplementary Fig. 5). The amount of each ddhNTP produced was quantified using a standard curve (Supplementary Fig. 5), and the results were plotted as a % of relative change to Wild-type enzyme. The -18C truncated variant only produced ddhUTP, whereas all other variants produced all four ddhNTPs (Fig. 1h). While the ability of -6C, -12C, and -15C truncated variants to produce ddhCTP, ddhGTP, or ddhATP was reduced, it was not abolished, unlike that of -18C truncated variant. The ability of -18C variant to selectively produce more ddhUTP than wild-type (WT) enzyme suggests that the IDR is not a mere “flap” or “lid” for the catalytic pocket. Instead, it dynamically participates in substrate selection and promiscuity. Because the -18C variant lacks the tripeptide TKP (Fig. 1d), this motif in the fungal enzyme plays the principal role in controlling IDR dynamic gating of substrates. The observations that the -18C variant only produces ddhUTP, but at a higher level than WT-TtSAND (Fig. 1h), and that it lost >80% of its chain-termination activity (Fig. 1f) confirm that IDR-dependent selection of different rNTP substrates to generate a ddhNTP cocktail (at least a mixture of ddhUTP and ddhCTP) is essential for efficient in-cell chain-termination activity.

### IDR tripeptide confers substrate selection and chain termination in humans

We tested whether the C-terminal IDR of hSAND (viperin) (Fig. 2a), which is highly conserved in vertebrates (Supplementary Fig. 2), confers substrate selection and promiscuity, as we observed in the fungal enzyme. While prior work on rSAND concluded that hSAND mainly produces ddhCTP,^34^ our in vitro and in-cell targeted metabolomic analysis in the presence of the electron donor CYB5R3^22^ (Supplementary Fig. 6 and Supplementary Table 1) confirmed that hSAND is not restricted to a single ddhNTP product (Figs. 2b-c). Instead, it displays intrinsic promiscuity, producing both ddhUTP and ddhCTP (Figs. 2b-c). Next, we created two C-terminal-truncated variants by removing 3 or 9 residues, hSAND-3C and -9C, respectively. In addition, three chimeric variants were created by adding 12 or 18 residues of the C-terminus of TtSAND to the WT-hSAND (WT+12TtC and WT+18TtC) or adding 18 residues of the C-terminus of TtSAND to the hSAND-9C truncated variant (hSAND-9C+18TtC). The chimeric variants were created to test if fungal IDR affects the human enzyme’s substrate selection. The C-terminal IDR variations did not affect the [4Fe-4S] cluster UV-visible absorbance spectrum and its content (Supplementary Fig. 7).

**Figure 2.**
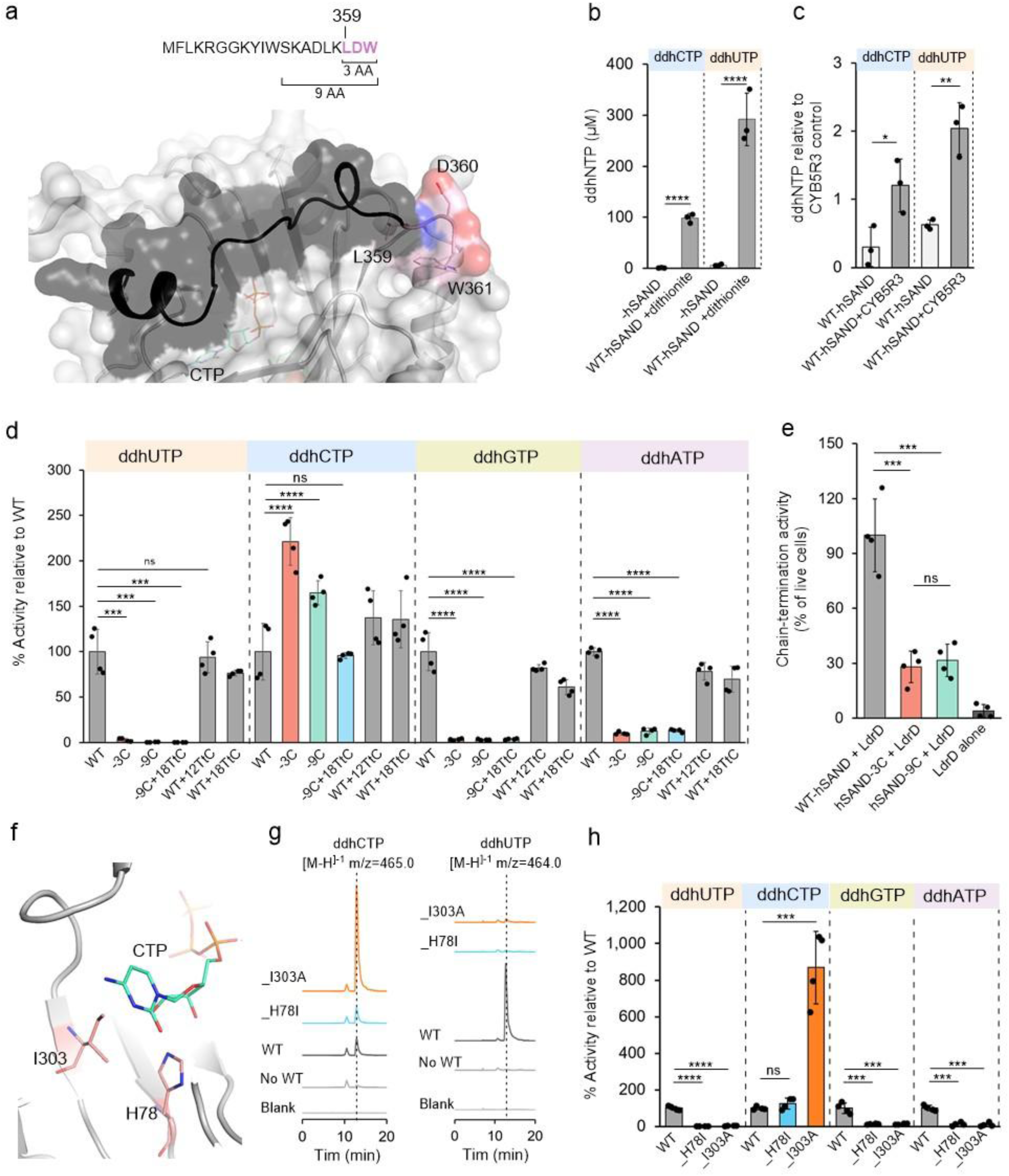
The IDR tripeptide LDW is required for substrate promiscuity of hSAND (viperin). **(a)** The AlphaFold-modeled structure of hSAND showing the C-terminal IDR (Black) and its tripeptide LDW (Pink). Three and nine amino acid (AA) residues of the C-terminus are highlighted. **(b-c)** hSAND has promiscuous activity, producing at least ddhCTP and ddhUTP **(b)** in a test tube and **(c)** in HEK293 cells. **(b)** Anaerobic dithionite solution was used as a reducing agent, and **(c)** the electron donor CYB5R3 was co-expressed with hSAND. Data are the average of three biologically independent measurements ± standard deviation. **(d)** Removing 3 (-3C) or 9 (-9C) amino acid residues from the C-terminus of hSAND makes the enzyme specific for ddhCTP formation, while chimeric variants with the C-terminal tripeptide (WT+12TtC and WT+18TtC) can produce all ddhNTPs like the WT enzyme. The amount of ddhNTP was quantified for each variant, and the results are plotted as the percentage of activity of wild-type (WT) hSAND. The data are averages of four measurements ± standard deviation, obtained from two different protein batches. **(e)** Removal of 3 or 9 residues from the C-terminal IDR abolished chain-termination activity in cells, as measured by the VITAS assay. **(f)** Two residues of the NTP-binding site, His78 and I303, were subject to mutagenesis. **(g-h)** hSAND active site is optimized to accept different rNTPs as substrates. **(g)** Ion chromatograms of ddhUTP or ddhCTP formation recorded in negative mode at the specified m/z ratio are shown as an example. **(h)** The amount of ddhNTP produced by hSAND_H78I and hSAND_I303A was quantified, and the data are presented as the % of activity relative to the wild-type (WT) SAND. Data are the average of four biologically independent measurements ± standard deviation. One-way ANOVA was used to calculate p-values. (*p value < 0.05; **p value < 0.01; ***p value < 0.001; ****p value < 0.0001).

Analysis of ddhNTP formation and quantification (Supplementary Fig. 8) showed that chimeric variants of WT-hSAND retained the promiscuous activity of WT-hSAND (Fig. 2d). However, the -3C and -9C truncated variants and the hSAND-9C+18TtC chimeric variant fully lost their ability to select UTP, GTP, and ATP substrates and were only selective for CTP and produced ddhCTP. Both -3C and– 9C variants showed improved activity toward ddhCTP (Fig. 2d). This clear separation and improved activity toward ddhCTP confirm that truncating the IDR C-terminus does not affect the [4Fe-4S] cluster. The IDR tripeptide LDW acts as the principal motif for dynamic gating and substrate selection. Therefore, removing the C-terminal tripeptide (LDW) was sufficient to make the enzyme CTP-specific. Even though removing the LDW tripeptide in hSAND-3C and -9C increased ddhCTP formation significantly (Fig. 2d), the in-cell chain-termination activity towards phage RNA polymerase decreased by more than 60-70% (Fig. 2e). Therefore, like TtSAND, the tripeptide is required for selection of different rNTPs as substrate, production of a cocktail of ddhNTPs, and efficient chain-termination activity inside cells.

### hSAND active site is optimized to accommodate different rNTPs

Our findings suggest that the IDR is a core regulatory element for substrate selection, enabling the production of a cocktail of antivirals. Consequently, the hSAND active site should be optimized for binding to different rNTPs and their transformation to their corresponding ddhNTPs. To test this, we aimed to determine whether active-site mutagenesis makes the enzyme specific to one substrate. We created two active-site variants of human SAND, hSAND_His78Ile and hSAND_Ile303Ala. His78 and Ile303 form the substrate-binding site (Fig. 2f), which is highly conserved in rats and humans (Supplementary Fig. 9). Neither the His78Ile nor the Ile303Ala substitution affected the enzyme’s [4Fe-4S] cluster content or the UV-visible absorbance spectrum of the purified proteins (Supplementary Fig. 9). The amount of ddhNTP produced was quantified (Supplementary Fig. 9). Both mutations abolished activity using UTP, GTP, and ATP as substrates and the formation of their corresponding ddhNTPs (Figs. 2g-h). The His78Ile mutation did not affect ddhCTP, and the Ile303Ala mutation increased ddhCTP formation by approximately 10-fold compared with the wild-type (WT) enzyme (Fig. 2g-h). These findings confirm that the hSAND active site is optimized to accept different rNTP substrates.

### The tripeptide acts as a latch to regulate the IDR ensemble

Next, we aimed to gain mechanistic insight into IDR-mediated substrate selection. Our modeled SAND structures (Fig. 2b and Supplementary Fig. 10a) suggest that the IDR tripeptide interacts with the radical-SAM domain. Enzyme kinetic analysis of C-terminal truncated variants lacking the tripeptide revealed increased accessibility of the only selected substrate (CTP for hSAND or UTP for TtSAND) (Fig. 3a and Supplementary Fig. 10b). The progress curves of ddhCTP or ddhUTP were obtained at different substrate concentrations, and the resulting data were fitted using a slow-binding inhibition model (Supplementary Methods). Deletion of the tripeptide coincided with a sharp increase in the initial rate (*v_i_*) (Fig. 3a and Supplementary Fig. 10b); however, the observed rate constant (*K_obs_*) (approximately 2 hr^-1^, Supplementary Table 2) was not affected. This pattern can only be explained by tripeptide loss, an altered IDR ensemble, and increased accessibility of one substrate to the active site: CTP for hSAND (viperin) and UTP for TtSAND.

**Figure 3.**
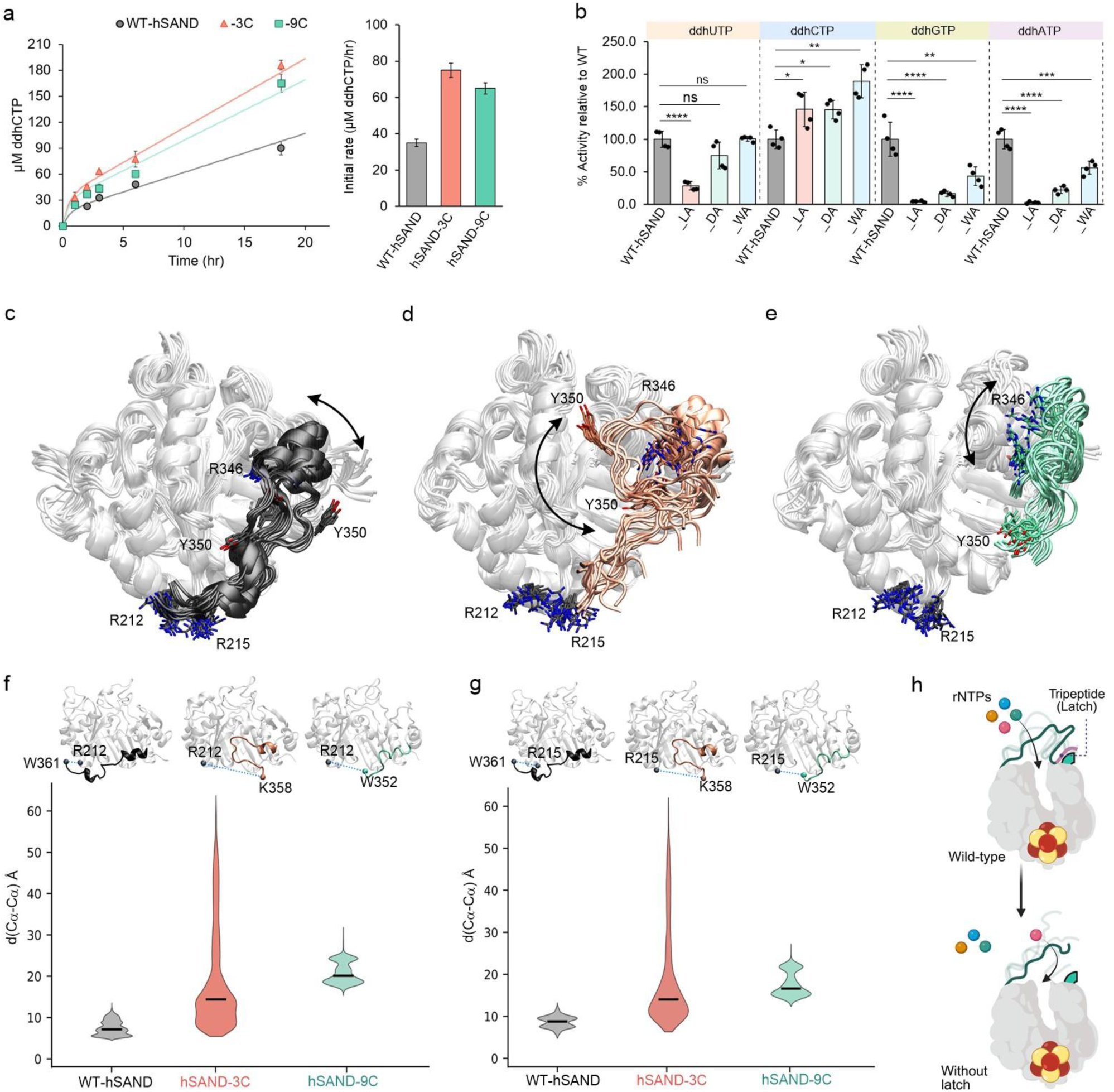
The IDR tripeptide acts as a latch and tunes the IDR ensemble. **(a)** Removal of the tripeptide in hSAND (-3C or -9C) increased the initial rate. **(b)** Alanine mutagenesis of the LDW tripeptide in hSAND confirms the tripeptide’s complex and dynamic role in substrate promiscuity. **(c-e)** Prediction of the (c) WT-hSAND, (d) hSAND-3C, or (e) hSAND-9C IDR ensemble in the absence of substrate CTP. Images are from three independent MD simulations per enzyme. Residues in the middle of the IDR (R346 and Y350) and two residues in the folded domain adjacent to the LDW tripeptide (R212 and R215) are labeled. **(f-g)** Distribution of the distances between the Cα atom of (f) R212 or (g) R215 and the Cα atom of the amino acid residue at the terminal C-tail of WT-hSAND, hSAND-3C, and hSAND-9C. The terminal residue is Trp361 in WT hSAND, Lys358 in hSAND-3C, or Trp352 in hSAND-9C. Violin plots summarize the distance distributions obtained from three independent MD simulations, with the median indicated by the horizontal line. **(h)** The proposed function of the tripeptide is to act as a latch that regulates the IDR ensemble and gates substrate selection. One-way ANOVA was used to calculate p-values. (*p value < 0.05; **p value < 0.01; ****p value < 0.0001).

To further probe the tripeptide’s role in modulating the IDR ensemble, we introduced alanine substitutions in the tripeptide. Mutations did not affect the FeS cluster (Supplementary Fig. 11-12). In TtSAND, where the TKP tripeptide lies in the middle of an extended IDR (Fig. 1d), alanine substitutions reduced activity towards all substrates relative to wild-type (Supplementary Fig. 10c). Therefore, the IDR in TtSAND lost its optimal ensemble, causing steric hindrance for substrate entry into the active site. In hSAND, where the LDW tripeptide lies at the IDR terminus (Fig. 2a), the L359A mutation abolished ddhUTP, ddhGTP, and ddhATP formation and moderately enhanced ddhCTP production relative to wild-type (Fig. 3b and Supplementary Fig. 10d). This pattern parallels that of tripeptide-truncated variants. The D360A and W361A substitutions moderately enhanced ddhCTP formation, reduced ddhGTP and ddhATP formation by > 50%, and had no obvious impact on ddhUTP formation (Fig. 3b). Collectively, these findings show that the tripeptide functions as a regulatory latch, anchoring the IDR into the radical-SAM domain to govern substrate selection.

### Structural insight into IDR ensemble

To test how the IDR tripeptide latch functions to govern substrate selection, we probed the IDR ensemble with and without the tripeptide latch. IDR structure in the absence of substrate is invisible to X-ray crystallography (Fig. 1b); therefore, we performed molecular dynamics (MD) simulations of WT-hSAND and its C-terminal truncation variants lacking the LDW tripeptide (Fig. 3c-e and Supplementary Fig. 13-14). In the simulations with the wild-type enzyme, the C-terminal tripeptide remains anchored in its context pocket (Fig. 3c). Removing the LDW tripeptide disrupts C-tail anchoring while preserving the overall structure and dynamics of the protein core (Fig. 3d-e). In WT-hSAND, the IDR can adopt a broad range of conformations over the active site. This specific ensemble covers the entry to the active site (Fig. 3c). The LDW motif anchored in its pocket as determined by measuring the Cα-Cα distance between the last amino acid residues of the IDR and two arginine residues of the context forming the tripeptide anchoring pocket, i.e., Arg212 (Fig. 3f) or Arg215 (Fig. 3g). Loss of the LDW tripeptide in the hSAND-3C and hSAND-9C truncated variants re-shaped the IDR ensemble (Figs. 3d-g). In hSAND-3C, the IDR adopted new conformations, as measured by Cα-Cα distances (Figs. 3f-g). This new ensemble leaves the active-site entrance exposed and fully open. Similarly, in the hSAND-9C truncated variant, the significantly shortened IDR also adopted new conformations, as indicated by the displacement of Arg346 (Fig. 3e) and variation in Cα-Cα distances (Figs. 3f-g). Although it samples a narrower conformational space than the IDR in hSAND-3C, its shorter length leaves the active-site entrance fully exposed. Therefore, consistent with enzyme kinetic analysis, we conclude that the IDR ensemble forms a dynamic gate at the active-site entrance, regulating substrate access through conformational rearrangements. (Fig. 3h). Removing the tripeptide abolishes this ensemble-mediated substrate selection by leaving the active-site entry open.

### Tripeptide context confers different outputs in rat and human enzymes

The tripeptide LDW is highly conserved in vertebrates (Supplementary Fig. 2). Therefore, we tested its function as the IDR latch in rat SAND (rSAND). The removal of LDW in rSAND (rSAND-3C) did not affect the FeS cluster (Supplementary Fig. 15). Quantification of ddhNTP formation (Fig. 4a) confirmed that tripeptide LDW is required for promiscuity in rSAND. It improved ddhCTP synthesis but abolished production of other ddhNTPs (Fig. 4a). This matches our observations for hSAND-3C (Fig. 2) and TtSAND-18C (Fig. 1), indicating that the tripeptide latch function is conserved. We noticed that rSAND produced approximately threefold more ddhCTP than ddhUTP (Fig. 4a), consistent with published data suggesting that rSAND mainly produces ddhCTP.^34,44^ This finding is opposed to our observation that hSAND produced three- to fourfold more ddhUTP than ddhCTP (Fig. 2b). Because the rat and human SANDs (viperin) have identical catalytic pockets (Supplementary Fig. 9) and we found that the IDR tripeptide LDW acts as a latch to control IDR ensemble gating of substrates, we hypothesized that the difference between hSAND and rSAND activities towards CTP or UTP is at least partially due to variation in the folded protein adjacent to the tripeptide LDW.

**Figure 4.**
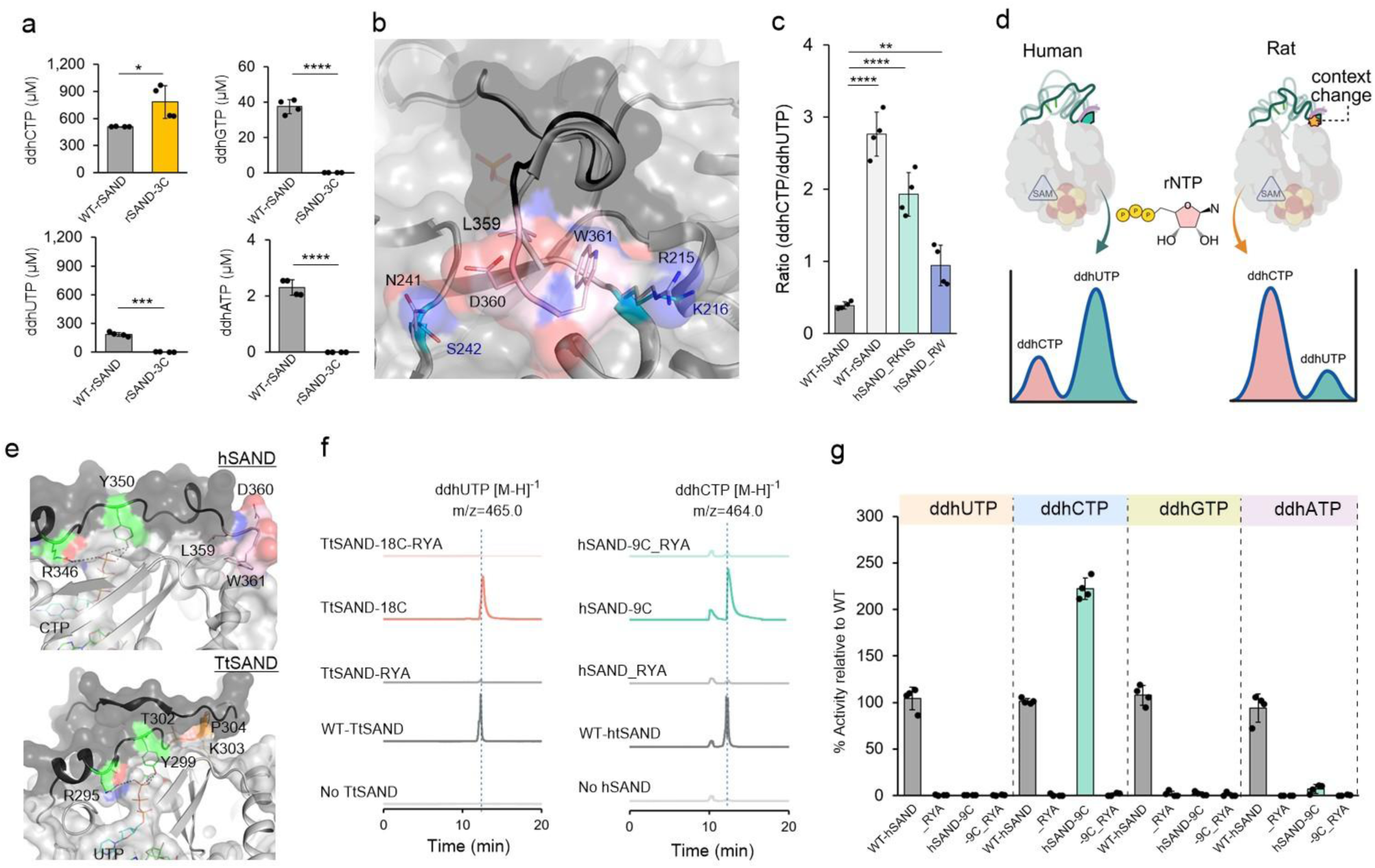
The IDR function is context-dependent and requires a clamp. **(a)** Removal of the tripeptide LDW from rSAND abolished the synthesis of ddhUTP, ddhGTP, or ddhATP but improved ddhCTP formation. **(b)** Alignment of human and rat SAND (viperin) structures shows that the LDW tripeptide context differs in two positions: Arg215 and Asn241 in hSAND (Gray) are replaced by lysine and serine in rSAND (Cyan), respectively. **(c)** Transplanting the rSAND context into hSAND, i.e., Arg215Lys and Asn241Ser substitutions in hSAND (hSAND_RKNS), recapitulates the activity profile of WT-rSAND. However, the Arg215Trp variant (hSAND_RW) did not. **(d)** A change in the context of the IDR tripeptide LDW in rat SAND, as compared to human SAND, alters the profile of ddhNTP products formed. **(e)** The modeled structures of hSAND and TtSAND show two highly conserved residues, an arginine and a tyrosine, that form hydrogen bonds with the terminal phosphate group of an rNTP substrate. **(f-g)** Substitution of arginine and tyrosine residues with alanine (designated as RYA mutant) fully abolishes the formation of all ddhNTPs. **(f)** Ion chromatograms of ddhCTP (hSAND) or ddhUTP (TtSAND) recorded at the specified m/z ratio are presented as examples. **(g)** The amount of each ddhNTP was quantified and the data are presented as % of activity relative to wild-type (WT) hSAND (viperin). Data are the average of two measurements ± errors from two different batches of protein. One-way ANOVA was used to calculate p-values. (*p value < 0.05; **p value < 0.01; ***p value < 0.001; ****p value < 0.0001; *****p value < 0.00001)

Alignment of the solved structure of rSAND and our modeled structure of hSAND shows two variations (Fig. 4b). Specifically, Arg215 and Asn241 in hSAND are replaced by lysine and serine in rSAND, respectively. Ancestral sequence reconstruction (ASR) (Supplementary Fig. 16) revealed that these variations are located in a substitution hotspot. Therefore, we replaced Arg215 and Asn241 in hSAND with the corresponding residues in rSAND, i.e., the Arg215Lys and Asn241Ser substitutions in hSAND (hSAND_RKNS). These mutations did not affect the FeS cluster (Supplementary Fig. 17). Quantification of the amount of ddhNTP produced by enzymes (Supplementary Fig. 18) revealed that while the ratio of ddhCTP to ddhUTP for WT-hSAND was approximately 0.5, for hSAND_RKNS and rSAND it was approximately 2.0 and 2.5, respectively (Fig. 4c). Therefore, hSAND_RKNS recapitulates rSAND activity by producing more ddhCTP than ddhUTP. Replacement of Arg215 in hSAND with the bulky amino acid tryptophan (hSAND_RW) did not affect the [4Fe-4S] cluster (Supplementary Fig. 17). Unlike the hSAND-RKNS variant, it showed a ddhCTP to ddhUTP ratio of approximately 1 (Fig. 4c and Supplementary Fig. 18). Similarly, a mutation in the folded protein domain adjacent to the TtSAND tripeptide TKP affected the activity profile of the enzyme (Supplementary Fig. 18). Therefore, we conclude that the amino acid residues adjacent to the IDR tripeptide latch regulate the IDR ensemble. Changes in these residues likely alter IDR ensemble kinetics or its conformational landscape, thereby changing the activity profile in human SAND compared with the rat enzyme (Fig. 4d).

### IDR has a clamp sequence

Because the tripeptide acts as a latch and tunes the IDR ensemble, we hypothesized that specific IDR residues serve as a clamp, participate in substrate recognition, and partially order the IDR structure upon substrate binding. In the solved structures of SANDs (Supplementary Fig. 19) and our modeled structures of hSAND and TtSAND (Fig. 4e), highly conserved arginine and tyrosine residues (Supplementary Fig. 20), Arg346 and Tyr350 in hSAND (viperin), form hydrogen bonds with the terminal phosphate group of an rNTP. We replaced these residues with alanine (designated as the _RYA variant) in both the wild-type (WT) and C-terminal-truncated variants of hSAND and TtSAND lacking the tripeptide. Mutations did not affect the [4Fe-4S] cluster (Supplementary Fig. 21). LC-MS analysis (Figure 4f) and quantification of ddhNTP formation (Supplementary Fig. 22) confirmed that substituting tyrosine and arginine with alanine fully abolished both fungal TtSAND (Supplementary Fig. 22) and hSAND (viperin) (Fig. 4f-g) activities toward all substrates. This effect was independent of the tripeptide latch. Therefore, we conclude that the highly conserved arginine and tyrosine residues of the RxxxY motif clamp various substrates and act as an essential motif for IDR-dependent ensemble gating.

### Discusion

While investigating the role of a C-terminal tryptophan in cofactor delivery to SAND, we found that the C-terminal region has an IDR vital for in-cell chain-termination activity. This serendipitous discovery revealed that the IDR is a key regulatory element of substrate selection and antiviral chain-termination activity from fungi to humans. The IDR has a tripeptide latch and a highly conserved clamp sequence. We demonstrate that an IDR-associated tripeptide dynamically regulates access to the active site and modulates production of different ddhNTPs in a context-dependent manner. Specifically, by transplanting the tripeptide context from rat SAND to the human enzyme, we showed that the human enzyme shifts its substrate preference from UTP to CTP, producing more ddhCTP similar to rat SAND. The IDR-dependent substrate-selection mechanism explains how rat and human enzymes with identical catalytic pockets differ in substrate preference and produce different ddhNTP profiles. Together, our data underscore that efficient chain termination relies on IDR-dependent substrate selection to produce a cocktail of ddhNTPs (at least two different ddhNTPs). This mechanism explains why ddhCTP alone is a poor chain terminator,^34,38,50^ why IDR deletion abolishes chain-termination activity,^41^ and why substitutions in the LDW, e.g., L359A, abrogate the enzyme’s ability to restrict ZIKA virus replication.^42^

We propose a clamp-and-latch function for the IDR to regulate species-specific antiviral output (Fig. 5). The clamp-and-latch action could modulate substrate promiscuity via two possible models. (i) Ensemble-gating and substrate recognition. The IDR ensemble acts as a kinetic gate. The conserved arginine and tyrosine act as a "clamp" to recognize and position different NTPs, and the tripeptide acts as a "latch" to modulate the IDR ensemble and stabilize substrates in the catalytic pocket. (ii) Ensemble-mediated allosteric substrate recognition. Guided by a clamp-and-latch mechanism, the IDR ensemble reshapes the catalytic pocket’s conformational dynamics. It stabilizes a distinct active-site conformation for each rNTP, biasing the pocket toward binding of that nucleotide and governing the enzyme’s substrate promiscuity. In both pathways, hydrogen-bonding of the clamp to the terminal phosphate yields a favorable negative enthalpy (ΔH) that offsets the entropic penalty (ΔS) of partial IDR ordering. This IDR partial ordering locks the substrate in the pocket to enable catalysis.

**Figure 5.**
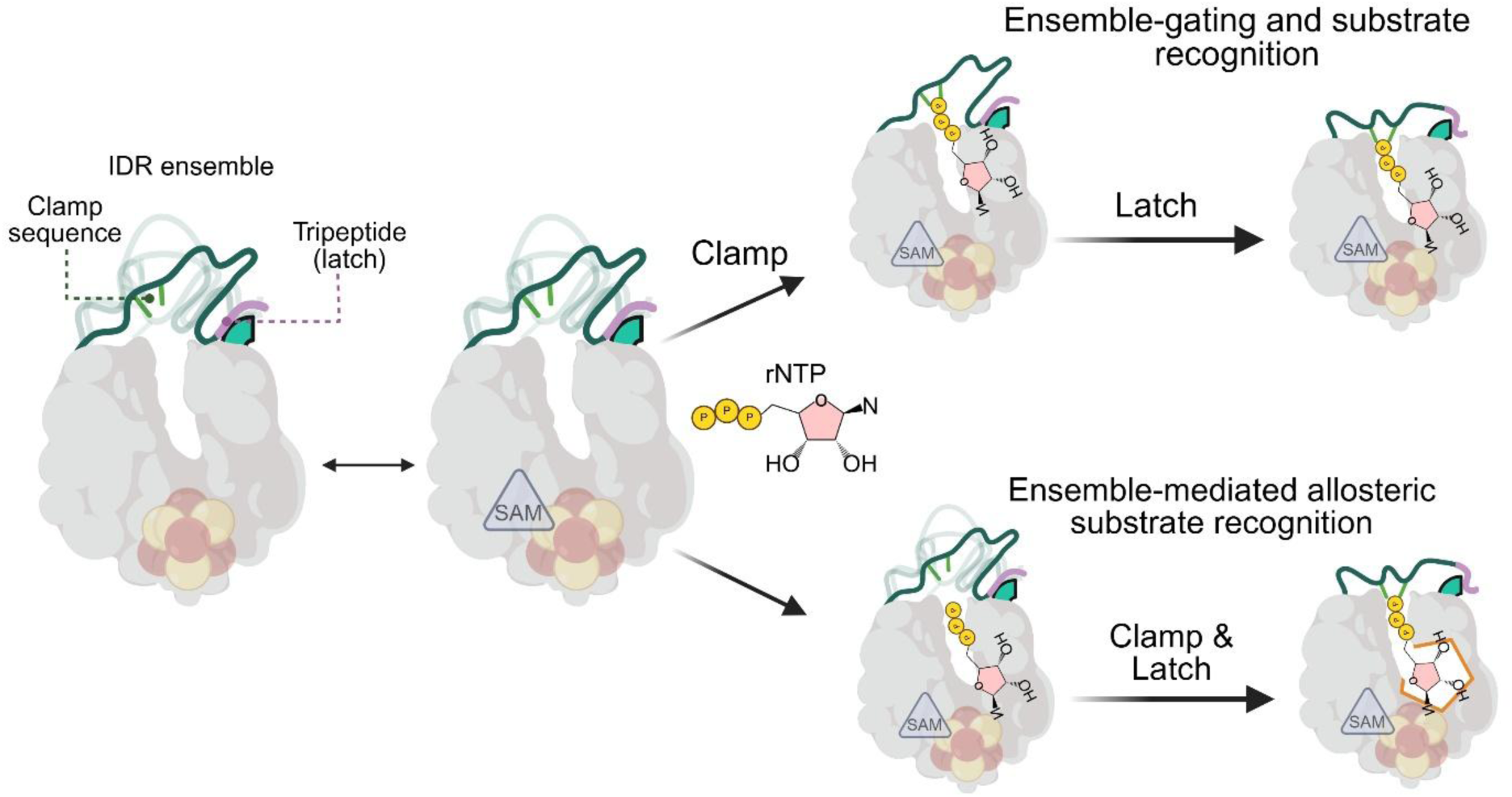
The proposed models for the clamp-and-latch action of IDR. The C-terminal IDR conformational ensemble regulates substrate promiscuity through a clamp-and-latch action. It can occur via two possible models. (i) Ensemble-gating and substrate recognition. The latch sequence and its context control the IDR ensemble and kinetically filter substrate selection. Conserved arginine and tyrosine act as a clamp to lock an rNTP substrate in the active site. (ii) Ensemble-mediated allosteric substrate recognition. The clamp-and-latch action of IDR regulates its specific ensemble, which directs the catalytic pocket’s conformational dynamics. The ensemble stabilizes a unique active-site geometry to accommodate a specific nucleoside substrate. This geometry is fixed upon clamp binding to rNTP to initiate catalysis.

This dynamic clamp-and-latch mechanism presents a paradigm shift in our understanding of enzyme-substrate recognition and promiscuity. However, several important questions remain and require future detailed analysis. First, the impact of inflammatory conditions and viral infection context on the clamp-and-latch action and SAND activity remains unclear. It is possible that, under specific cellular conditions, the IDR clamp-and-latch action and ensemble are modulated to tailor the antiviral response or, conversely, to subvert host defense. Second, the exact function of individual IDR residues in clamp-and-latch, and how residues in the folded protein adjacent to the IDR regulate its ensemble and substrate selection, remains unknown. Future detailed mutagenesis studies, combined with structural studies using NMR and MD simulations, should help address these fundamental questions and elucidate the molecular mechanism of the IDR’s clamp-and-latch action.

Looking forward, our findings suggest that genetic variations within the IDR region or its adjacent folded context may confer viral susceptibility. They will stimulate functional studies of IDRs in numerous other enzymes. These IDR studies are expected to generate essential training data for machine-learning models applied to drug discovery and enzyme engineering.

## Online Methods

### Chemical

All chemicals were reagent grade and purchased from ThemoFisher Scientific (Waltham, MA, USA), Fisher Scientific, (Loughborough, UK) or Merck, (Rahway, NJ, USA). All anaerobic buffers were prepared a day prior to the experiments and stored in the glovebox overnight to ensure complete removal of dioxygen.

### Genes and cloning

The *E. coli* codon-optimized synthetic genes for different enzymes were obtained from GeneArt (ThermoFisher) either in pMQ plasmid or final expression vector, pBAD/His C. Cloning from the pMQ vector to the pBAD/His C vector was performed using 5′ and 3′ restriction sites for KpnI and HindIII, respectively. The presence of the correct insert was confirmed using restriction enzyme double digestion and DNA sequencing (GENEWIZ). The plasmids were transformed into *E. Coli* TOP10 cells, and a glycerol stock of the cells was prepared and stored at −80 °C. The glycerol stock was used to prepare LB Agar (100 μm ampicillin) plates for protein expression. All cloned enzymes lacked the N-terminal hydrophobic domain. Residue numbering is based on the full-length proteins.

**TtSAND-18C** MPVSVNYHFTRQCNKSCGFCFHTATTSHIENLERQQAALRLLAQAGMRKINFAGGEPFLYPKALGAMIDFCKRELRLESVSVVTNGSLVREDFLRRHAGQLDILAVSCDSFNEQTNVAIGRGSGDQVPRLYQIAEWCRTYGVLFKINTVVNRLNVAEDMNAHIAALRPFRWKCFQVLIVRGENDSDETLRNAHRFTISDEEFEAFCERHRGQACLVPEPNRLMAKSYLILDEYLRFLDRTGKQPSKPILEVGVEKALESVYWDEEAFVERGGLYEW

**TtSAND-6C** MPVSVNYHFTRQCNKSCGFCFHTATTSHIENLERQQAALRLLAQAGMRKINFAGGEPFLYPKALGAMIDFCKRELRLESVSVVTNGSLVREDFLRRHAGQLDILAVSCDSFNEQTNVAIGRGSGDQVPRLYQIAEWCRTYGVLFKINTVVNRLNVAEDMNAHIAALRPFRWKCFQVLIVRGENDSDETLRNAHRFTISDEEFEAFCERHRGQACLVPEPNRLMAKSYLILDEYLRFLDRTGKQPSKPILEVGVEKALESVYWDEEAFVERGGLYEWTKPANGESGCGS

**TtSAND-12C** MPVSVNYHFTRQCNKSCGFCFHTATTSHIENLERQQAALRLLAQAGMRKINFAGGEPFLYPKALGAMIDFCKRELRLESVSVVTNGSLVREDFLRRHAGQLDILAVSCDSFNEQTNVAIGRGSGDQVPRLYQIAEWCRTYGVLFKINTVVNRLNVAEDMNAHIAALRPFRWKCFQVLIVRGENDSDETLRNAHRFTISDEEFEAFCERHRGQACLVPEPNRLMAKSYLILDEYLRFLDRTGKQPSKPILEVGVEKALESVYWDEEAFVERGGLYEWTKPANG

**TtSAND-15C** MPVSVNYHFTRQCNKSCGFCFHTATTSHIENLERQQAALRLLAQAGMRKINFAGGEPFLYPKALGAMIDFCKRELRLESVSVVTNGSLVREDFLRRHAGQLDILAVSCDSFNEQTNVAIGRGSGDQVPRLYQIAEWCRTYGVLFKINTVVNRLNVAEDMNAHIAALRPFRWKCFQVLIVRGENDSDETLRNAHRFTISDEEFEAFCERHRGQACLVPEPNRLMAKSYLILDEYLRFLDRTGKQPSKPILEVGVEKALESVYWDEEAFVERGGLYEWTKP

**TtSAND-SUMO** MPVSVNYHFTRQCNKSCGFCFHTATTSHIENLERQQAALRLLAQAGMRKINFAGGEPFLYPKALGAMIDFCKRELRLESVSVVTNGSLVREDFLRRHAGQLDILAVSCDSFNEQTNVAIGRGSGDQVPRLYQIAEWCRTYGVLFKINTVVNRLNVAEDMNAHIAALRPFRWKCFQVLIVRGENDSDETLRNAHRFTISDEEFEAFCERHRGQACLVPEPNRLMAKSYLILDEYLRFLDRTGKQPSKPILEVGVEKALESVYWDEEAFVERGGLYEWTKPANGESGCGSKKELEWMHHHHHHGSLQDSEVNQEAKPEVKPEVKPETHINLKVSDGSSEIFFKIKKTTPLRRLMEAFAKRQGKEMDSLTFLYDGIEIQADQTPEDLDMEDNDIIEAHREQIGGATY

**TtSAND_RYA (Arg295Ala and Tyr299Ala)** MPVSVNYHFTRQCNKSCGFCFHTATTSHIENLERQQAALRLLAQAGMRKINFAGGEPFLYPKALGAMIDFCKRELRLESVSVVTNGSLVREDFLRRHAGQLDILAVSCDSFNEQTNVAIGRGSGDQVPRLYQIAEWCRTYGVLFKINTVVNRLNVAEDMNAHIAALRPFRWKCFQVLIVRGENDSDETLRNAHRFTISDEEFEAFCERHRGQACLVPEPNRLMAKSYLILDEYLRFLDRTGKQPSKPILEVGVEKALESVYWDEEAFVEAGGLAEWTKPANGESGCGSKKELEW

**TtSAND-18C_RYA (Arg295Ala and Tyr299Ala)** MPVSVNYHFTRQCNKSCGFCFHTATTSHIENLERQQAALRLLAQAGMRKINFAGGEPFLYPKALGAMIDFCKRELRLESVSVVTNGSLVREDFLRRHAGQLDILAVSCDSFNEQTNVAIGRGSGDQVPRLYQIAEWCRTYGVLFKINTVVNRLNVAEDMNAHIAALRPFRWKCFQVLIVRGENDSDETLRNAHRFTISDEEFEAFCERHRGQACLVPEPNRLMAKSYLILDEYLRFLDRTGKQPSKPILEVGVEKALESVYWDEEAFVEAGGLAEW

**TtSAND_TKP-AAA** MPVSVNYHFTRQCNKSCGFCFHTATTSHIENLERQQAALRLLAQAGMRKINFAGGEPFLYPKALGAMIDFCKRELRLESVSVVTNGSLVREDFLRRHAGQLDILAVSCDSFNEQTNVAIGRGSGDQVPRLYQIAEWCRTYGVLFKINTVVNRLNVAEDMNAHIAALRPFRWKCFQVLIVRGENDSDETLRNAHRFTISDEEFEAFCERHRGQACLVPEPNRLMAKSYLILDEYLRFLDRTGKQPSKPILEVGVEKALESVYWDEEAFVERGGLYEWAAAANGESGCGSKKELEW

**TtSAND-TA** MPVSVNYHFTRQCNKSCGFCFHTATTSHIENLERQQAALRLLAQAGMRKINFAGGEPFLYPKALGAMIDFCKRELRLESVSVVTNGSLVREDFLRRHAGQLDILAVSCDSFNEQTNVAIGRGSGDQVPRLYQIAEWCRTYGVLFKINTVVNRLNVAEDMNAHIAALRPFRWKCFQVLIVRGENDSDETLRNAHRFTISDEEFEAFCERHRGQACLVPEPNRLMAKSYLILDEYLRFLDRTGKQPSKPILEVGVEKALESVYWDEEAFVERGGLYEWAKPANGESGCGSKKELEW

**TtSAND-KA** MPVSVNYHFTRQCNKSCGFCFHTATTSHIENLERQQAALRLLAQAGMRKINFAGGEPFLYPKALGAMIDFCKRELRLESVSVVTNGSLVREDFLRRHAGQLDILAVSCDSFNEQTNVAIGRGSGDQVPRLYQIAEWCRTYGVLFKINTVVNRLNVAEDMNAHIAALRPFRWKCFQVLIVRGENDSDETLRNAHRFTISDEEFEAFCERHRGQACLVPEPNRLMAKSYLILDEYLRFLDRTGKQPSKPILEVGVEKALESVYWDEEAFVERGGLYEWTAPANGESGCGSKKELEW

**hSAND-3C** MPTTPTSVNYHFTRQCNYKCGFCFHTAKTSFVLPLEEAKRGLLLLKEAGMEKINFSGGEPFLQDRGEYLGKLVRFCKVELRLPSVSIVSNGSLIRERWFQNYGEYLDILAISCDSFDEEVNVLIGRGQGKKNHVENLQKLRRWCRDYRVAFKINSVINRFNVEEDMTEQIKALNPVRWKVFQCLLIEGENCGEDALREAERFVIGDEEFERFLERHKEVSCLVPESNQKMKDSYLILDEYMRFLNCRKGRKDPSKSILDVGVEEAIKFSGFDEKMFLKRGGKYIWSKADLK

**hSAND-9C** MPTTPTSVNYHFTRQCNYKCGFCFHTAKTSFVLPLEEAKRGLLLLKEAGMEKINFSGGEPFLQDRGEYLGKLVRFCKVELRLPSVSIVSNGSLIRERWFQNYGEYLDILAISCDSFDEEVNVLIGRGQGKKNHVENLQKLRRWCRDYRVAFKINSVINRFNVEEDMTEQIKALNPVRWKVFQCLLIEGENCGEDALREAERFVIGDEEFERFLERHKEVSCLVPESNQKMKDSYLILDEYMRFLNCRKGRKDPSKSILDVGVEEAIKFSGFDEKMFLKRGGKYIW

**hSAND-9C+18TtC** MPTTPTSVNYHFTRQCNYKCGFCFHTAKTSFVLPLEEAKRGLLLLKEAGMEKINFSGGEPFLQDRGEYLGKLVRFCKVELRLPSVSIVSNGSLIRERWFQNYGEYLDILAISCDSFDEEVNVLIGRGQGKKNHVENLQKLRRWCRDYRVAFKINSVINRFNVEEDMTEQIKALNPVRWKVFQCLLIEGENCGEDALREAERFVIGDEEFERFLERHKEVSCLVPESNQKMKDSYLILDEYMRFLNCRKGRKDPSKSILDVGVEEAIKFSGFDEKMFLKRGGKYIWTKPANGESGCGSKKELEW

**hSAND+18TtC** MPTTPTSVNYHFTRQCNYKCGFCFHTAKTSFVLPLEEAKRGLLLLKEAGMEKINFSGGEPFLQDRGEYLGKLVRFCKVELRLPSVSIVSNGSLIRERWFQNYGEYLDILAISCDSFDEEVNVLIGRGQGKKNHVENLQKLRRWCRDYRVAFKINSVINRFNVEEDMTEQIKALNPVRWKVFQCLLIEGENCGEDALREAERFVIGDEEFERFLERHKEVSCLVPESNQKMKDSYLILDEYMRFLNCRKGRKDPSKSILDVGVEEAIKFSGFDEKMFLKRGGKYIWSKADLKLDWTKPANGESGCGSKKELEW

**hSAND+12TtC** MPTTPTSVNYHFTRQCNYKCGFCFHTAKTSFVLPLEEAKRGLLLLKEAGMEKINFSGGEPFLQDRGEYLGKLVRFCKVELRLPSVSIVSNGSLIRERWFQNYGEYLDILAISCDSFDEEVNVLIGRGQGKKNHVENLQKLRRWCRDYRVAFKINSVINRFNVEEDMTEQIKALNPVRWKVFQCLLIEGENCGEDALREAERFVIGDEEFERFLERHKEVSCLVPESNQKMKDSYLILDEYMRFLNCRKGRKDPSKSILDVGVEEAIKFSGFDEKMFLKRGGKYIWSKADLKLDWTKPANGESGCGS

**hSAND_RYA (Arg346Ala & Tyr350Ala)** MPTTPTSVNYHFTRQCNYKCGFCFHTAKTSFVLPLEEAKRGLLLLKEAGMEKINFSGGEPFLQDRGEYLGKLVRFCKVELRLPSVSIVSNGSLIRERWFQNYGEYLDILAISCDSFDEEVNVLIGRGQGKKNHVENLQKLRRWCRDYRVAFKINSVINRFNVEEDMTEQIKALNPVRWKVFQCLLIEGENCGEDALREAERFVIGDEEFERFLERHKEVSCLVPESNQKMKDSYLILDEYMRFLNCRKGRKDPSKSILDVGVEEAIKFSGFDEKMFLKAGGKAIWSKADLKLDW

**hSAND-9C_RYA (Arg346Ala & Tyr350Ala)** MPTTPTSVNYHFTRQCNYKCGFCFHTAKTSFVLPLEEAKRGLLLLKEAGMEKINFSGGEPFLQDRGEYLGKLVRFCKVELRLPSVSIVSNGSLIRERWFQNYGEYLDILAISCDSFDEEVNVLIGRGQGKKNHVENLQKLRRWCRDYRVAFKINSVINRFNVEEDMTEQIKALNPVRWKVFQCLLIEGENCGEDALREAERFVIGDEEFERFLERHKEVSCLVPESNQKMKDSYLILDEYMRFLNCRKGRKDPSKSILDVGVEEAIKFSGFDEKMFLKAGGKAIW

**hSAND_RKNS (R215K and N241S)** MPLPTTPTSVNYHFTRQCNYKCGFCFHTAKTSFVLPLEEAKRGLLLLKEAGMEKINFSGGEPFLQDRGEYLGKLVRFCKVELRLPSVSIVSNGSLIRERWFQNYGEYLDILAISCDSFDEEVNVLIGRGQGKKNHVENLQKLRRWCRDYKVAFKINSVINRFNVEEDMTEQIKALSPVRWKVFQCLLIEGENCGEDALREAERFVIGDEEFERFLERHKEVSCLVPESNQKMKDSYLILDEYMRFLNCRKGRKDPSKSILDVGVEEAIKFSGFDEKMFLKRGGKYIWSKADLKLDW

**hSAND_RW (R215W)** MPLPTTPTSVNYHFTRQCNYKCGFCFHTAKTSFVLPLEEAKRGLLLLKEAGMEKINFSGGEPFLQDRGEYLGKLVRFCKVELRLPSVSIVSNGSLIRERWFQNYGEYLDILAISCDSFDEEVNVLIGRGQGKKNHVENLQKLRRWCRDYWVAFKINSVINRFNVEEDMTEQIKALNPVRWKVFQCLLIEGENCGEDALREAERFVIGDEEFERFLERHKEVSCLVPESNQKMKDSYLILDEYMRFLNCRKGRKDPSKSILDVGVEEAIKFSGFDEKMFLKRGGKYIWSKADLKLDW

**hSAND_WA (W361A)** MPTTPTSVNYHFTRQCNYKCGFCFHTAKTSFVLPLEEAKRGLLLLKEAGMEKINFSGGEPFLQDRGEYLGKLVRFCKVELRLPSVSIVSNGSLIRERWFQNYGEYLDILAISCDSFDEEVNVLIGRGQGKKNHVENLQKLRRWCRDYRVAFKINSVINRFNVEEDMTEQIKALNPVRWKVFQCLLIEGENCGEDALREAERFVIGDEEFERFLERHKEVSCLVPESNQKMKDSYLILDEYMRFLNCRKGRKDPSKSILDVGVEEAIKFSGFDEKMFLKRGGKYIWSKADLKLDA

**hSAND_DA (D360A)** MPTTPTSVNYHFTRQCNYKCGFCFHTAKTSFVLPLEEAKRGLLLLKEAGMEKINFSGGEPFLQDRGEYLGKLVRFCKVELRLPSVSIVSNGSLIRERWFQNYGEYLDILAISCDSFDEEVNVLIGRGQGKKNHVENLQKLRRWCRDYRVAFKINSVINRFNVEEDMTEQIKALNPVRWKVFQCLLIEGENCGEDALREAERFVIGDEEFERFLERHKEVSCLVPESNQKMKDSYLILDEYMRFLNCRKGRKDPSKSILDVGVEEAIKFSGFDEKMFLKRGGKYIWSKADLKLAW

**hSAND_LA (L359A)** MPTTPTSVNYHFTRQCNYKCGFCFHTAKTSFVLPLEEAKRGLLLLKEAGMEKINFSGGEPFLQDRGEYLGKLVRFCKVELRLPSVSIVSNGSLIRERWFQNYGEYLDILAISCDSFDEEVNVLIGRGQGKKNHVENLQKLRRWCRDYRVAFKINSVINRFNVEEDMTEQIKALNPVRWKVFQCLLIEGENCGEDALREAERFVIGDEEFERFLERHKEVSCLVPESNQKMKDSYLILDEYMRFLNCRKGRKDPSKSILDVGVEEAIKFSGFDEKMFLKRGGKYIWSKADLKADW

**Rat SAND (rSAND)** MQPTTPVSVNYHFTRQCNYKCGFCFHTAKTSFVLPLEEAKRGLLLLKQAGMEKINFSGGEPFLQDRGEYLGKLVRFCKEELALPSVSIVSNGSLIRERWFKDYGDYLDILAISCDSFDEQVNVLIGRGQGKKNHVENLQKLRKWCRDYKVAFKINSVINRFNVDEDMNEHIKALSPVRWKVFQCLLIEGENSGEDALREAERFLISNEEFEAFLQRHKDVSCLVPESNQKMKDSYLILDEYMRFLNCTGGRKDPSRSILDVGVEEAIKFSGFDEKMFLKRGGKYVWSKADLKLDW

**rSAND-3C** MQPTTPVSVNYHFTRQCNYKCGFCFHTAKTSFVLPLEEAKRGLLLLKQAGMEKINFSGGEPFLQDRGEYLGKLVRFCKEELALPSVSIVSNGSLIRERWFKDYGDYLDILAISCDSFDEQVNVLIGRGQGKKNHVENLQKLRKWCRDYKVAFKINSVINRFNVDEDMNEHIKALSPVRWKVFQCLLIEGENSGEDALREAERFLISNEEFEAFLQRHKDVSCLVPESNQKMKDSYLILDEYMRFLNCTGGRKDPSRSILDVGVEEAIKFSGFDEKMFLKRGGKYVWSKADLK

### Expression and purification

Cells were spread on an LB agar plate containing ampicillin (100 µg/ml). The next day, a single colony was picked up and inoculated into 50 mL LB medium containing ampicillin (100 µg/ml). The mini culture was grown overnight (16-18 hours) in a shaker incubator at 200 rpm and 37 °C, and the next day, it was added to 500 mL TB medium, and the flasks were incubated in a shaker/incubator at 200 rpm and 37 °C. The expression of all enzymes was induced by the addition of 0.04% W/v arabinose (final concentration). Six-seven hours after induction, cells were harvested using centrifugation at 5000 x g at 4 °C for 20 min. The cell pellet was stored at −80 °C. A volume of the cell pellet was mixed with an equal volume of lysis buffer (containing 2% Triton X-100, 50 mM Tris-HCl pH 7.6, 300 mM NaCl, 0.05mg PMSF, 20mg Lysozyme, and xxx mg PMSF). The cell suspension was aliquoted into 5 mL and lysed using sonication: power 30%, 10 cycles, every 30 seconds on and 10 seconds off. During sonication, samples were kept on ice to prevent overheating. Subsequently, the lysate was subjected to centrifugation (12000×g) for 20 minutes at 4°C. The supernatant was collected and immediately transferred into an anaerobic glovebox (O_2_ < 5 ppm) (Belle Technologies). Anaerobic purification was achieved using a gravity column packed with HisPure Ni^2+^ resin (Thermo Scientific). The following buffers were used: Equilibration buffer (50 mM Tris, 300 mM NaCl, 2% triton and 5 mM imidazole, pH 7.5); wash buffer (50 mM Tris, 300 mM NaCl, 2% v/v triton and 10 mM imidazole, pH 7.5); and elution buffer (50 mM Tris, 300 mM NaCl, 2% v/v triton and 500 mM imidazole, pH 7.5). Subsequently, the PD10 desalting column was used to exchange the buffer and remove imidazole and Triton. The final buffer was 50 mM MOPS (3-(Morpholin-4-yl) propane-1-sulfonic acid) and 100 mM NaCl, pH 7.0 for hSAND or 50 mM MOPS (3-(Morpholin-4-yl) propane-1-sulfonic acid) and 300 mM NaCl, pH 7.0 for TtSAND. The sample was flash frozen in liquid nitrogen and stored in liquid N_2_. Protein purity was confirmed by 10% SDS/PAGE, and protein concentration was measured using the BCA assay. Protein concentrations are the average of three independent measurements.

### Analysis of [4Fe-4S] cluster content

To estimate the percentage of clusters per protein, the ferene assay was used to determine the amount of iron. The assay was performed as described before ^51^. First, a standard curve was prepared using ferrous ammonium sulfate as standards and based on concentrations described in Table S2. An Fe (II) solution stock (1 mg/ml) was prepared by dissolving ferrous ammonium sulfate hexahydrate in Milli-Q water. Subsequently, this solution was further diluted using Milli-Q water to prepare a solution of 0.1 mg/ml. This working stock was used to prepare the samples for the standard curve (Table S2). To prepare samples using protein, each purified enzyme solution was diluted 2 times, 5 times, and 10 times. Subsequently, 100 µl of each standard or sample was mixed with 100 µl of 1% HCl and incubated at 100°C for 10 minutes. Next, to each sample or standard, 500 µL of ammonium acetate 15% w/v, 100 µL of SDS 2.5%, 100 µL of ascorbic acid 4%, and 100 µL of ferene 1.5% w/v were added in order. The sample and standards were subjected to centrifugation at 13000 rpm for 15 minutes. Finally, they were analyzed using UV-visible spectrophotometry by recording absorbance at 593 nm. Milli-Q water was used as a blank. After determining the amount of iron, the amount of cluster per protein was determined using Equation 1.

**Table method.**
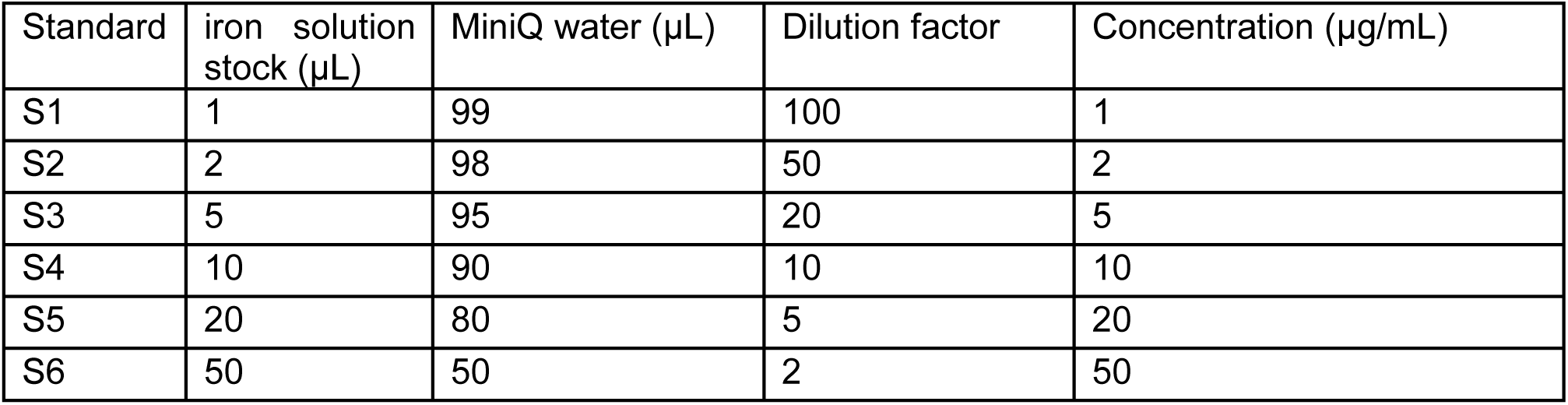
1. The samples were prepared to create the standard curve.

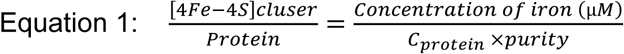

In which

c_protein_: Concentration of protein as measured by BCA.

Purity: The percentage of purified protein as determined by SDS analysis.

### UV-visible spectrophotometry

To record the UV–visible absorbance spectra, samples were added anaerobically inside the glovebox into a 1 mL disposable plastic cuvette (BRAND™, Willimantic, CT, USA) with a path length of 1 cm and a silicone lid (BRAND™). Each sample was transported outside, and the UV–visible absorbance spectra were recorded immediately using an Agilent UV-visible spectrophotometer. Spectra were recorded from 300 to 700 nm. Measurements were performed at room temperature.

### Preparation of samples for liquid chromatography–mass spectrometry (LC–MS) analysis of 3՛-deoxy-3՛,4՛-didehydro-CTP (ddhCTP), 3՛-deoxy-3՛,4՛-didehydro-UTP (ddhUTP), 3՛-deoxy-3՛,4՛-didehydro-GTP (ddhGTP) and 3՛-deoxy-3՛,4՛-didehydro-CTP (ddhATP) formation

Anaerobic buffer, 50 mm MOPS (pH 7.0) containing 100 mm NaCl for hSAND or 50 mm MOPS (pH 7.0) containing 300 mm NaCl for TtSAND, was used to prepare stock solutions of SAM, cytidine triphosphate (CTP), uridine triphosphate (UTP), guanosine triphosphate (GTP), adenosine triphosphate (ATP) and sodium dithionite (SD) under anaerobic conditions in a glovebox (Belle Technology (O_2_ < 5 ppm)). An aliquot of chemicals was transported into the glovebox and mixed with the buffer under anaerobic conditions. The buffer was prepared a day prior to the experiments and stored in the glovebox overnight to ensure complete anaerobicity. For accurate quantification and to account for day-to-day variability, we prepared a sample using wild-type (WT) SAND as an external standard and ran it in parallel with all measurements. External standards were used since sample preparation is minimal and clean. WT-hSAND was used as the standard for all measurements with hSAND variants and rSAND, and WT-TtSAND was used as the standard for all measurements with TtSAND variants (Table 1). We prepared a standard curve for ddhCTP, ddhUTP, ddhGTP, and ddhATP using enzymatic production of each ddh analogue and quantifying the amount of rNTP consumed and converted to its corresponding ddhNTP (Figure S23). The complete reaction contained 100 μL of purified hSAND or TtSAND mutant solution, 10 μL 50 mM SAM (S-(5′-adenosyl)-L-methionine (S-methyl-13C) chloride), 5 μL 40 mM CTP, GTP, ATP, or UTP, and 20 μL 80 mM Na_2_S_2_O_4_ (sodium dithionite). The internal standard had 100 μL of purified WT-hSAND or WT-TtSAND enzyme, 10 μL 50mM SAM (S-(5′-adenosyl)-l-methionine-(S-methyl-13C) chloride), 5μL of 40mM CTP, GTP, ATP, or UTP, and 20 μL of 80mM Na_2_S_2_O_4_ (sodium dithionite) as described in Table S1. When needed, an anaerobic MOPS buffer was added to adjust the volume. A negative control sample was prepared. In the negative control, the enzyme was replaced with buffer. The reaction mixture was incubated anaerobically. Following incubation for a specific time (typically overnight, 20-22 hours based on cell biological data showing ddhCTP formation after circa 16-24 hours ^34^), the sample was centrifuged at 13400 rpm for 3 min. Then, the reaction was added to a 0.5-mL Amicon Ultra Centrifugal Filter 3 kDa (Merck) and centrifuged at 13400 rpm for 10 min. The resulting flow-through was then carefully transferred to an HPLC low-volume (300 μL) recovery vial (Fisher Scientific) and sealed tightly with a polypropylene cap (Fisher Scientific). Concentrations of hSAND and its variants (consisting of the FeS cluster) were 40 µM unless otherwise stated. Concentrations of TtSAND and its variants (consisting of the FeS cluster) were 100 µM. The concentration of rSAND and its variants (consisting of the FeS cluster) was 120 µM.

**Table Method.**
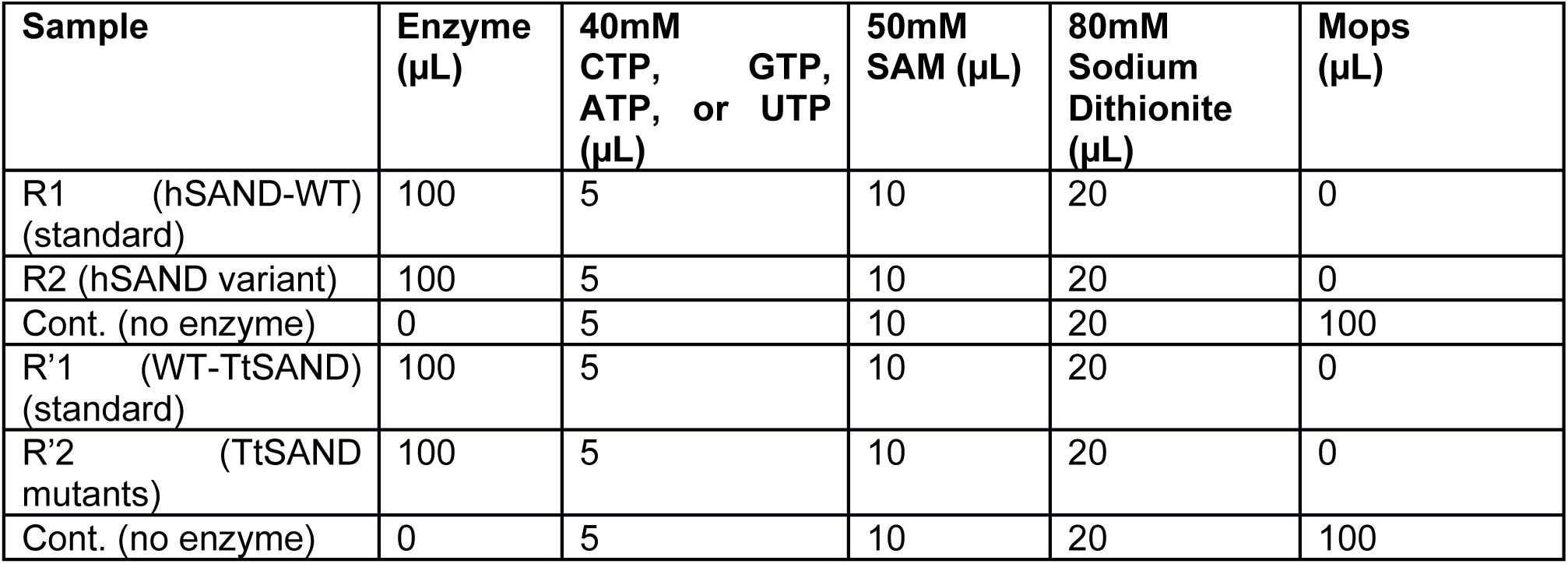
2. Details of reaction setup using different enzymes for analysis of ddhNTP formation.

### Liquid chromatography-Mass spectrometry

Measurements were performed using an Agilent InfinityLab G6160A LC/MSD iQ mass spectrometer equipped with an Agilent 1260 Infinity II liquid chromatography system. The column was an Agilent ZORBAX RR HILIC Plus Column, 2.1 x 100 mm, 3.5 µm, or RRHD 2.1x150 mm, 1.8 µm. Instrument control and data processing were performed using OpenLab CDS (Agilent Technologies, Santa Clara, CA, USA). The system was calibrated on the day of the analysis. Electrospray source conditions were adjusted to maximize sensitivity, and the detection mode was set to detect negative (−) ions. For each run, 50 μL of solution was injected with a flow rate of 0.2 mL·min−1 and an oven temperature of 45 °C:

Buffer A: 90 vol.% MeCN (acetonitrile) (Fisher, 99.9%): 10 vol.% LC–MS grade water (Fisher), 20 mM ammonium acetate, pH 7.4–7.5.

Buffer B: 10 vol.% LC–MS-grade water (Fisher), 20 mM ammonium acetate, pH 7.4–7.5. Column Flow rate: 0.2 mL·min−1.

Gradient: [0–1 min]: 100 : 0 (A : B); [1–10 min] 100 : 0 (A : B) linearly changed to 10 : 90 (A : B); [10–15 min] 10 : 90 (A : B); [15 : 17] linearly changed to 100 : 0 (A : B) and [17–35 min] 100–0 (A : B). Analysis of the LC–MS data was performed using Open Lab CDS software. Experiments with each variant were performed in parallel to the WT enzyme as a positive control. The areas under the curve were obtained and used to calculate the amount of each ddhNTP using a standard curve. The data were corrected for any variation in protein concentration relative to the WT control (Equation 2).

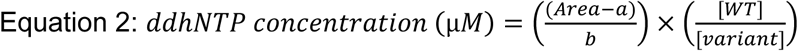

In which

Area: area under the curve

a: the Y-intercept for the standard curve b: the slope for the standard curve

[WT]: concentration (µM) of wild-type enzyme

[variant]: concentration (µM) of a variant

### Analysis of enzyme kinetics

Samples for analysis of the kinetics of TtSAND and hSAND (wild-type and C-terminal truncated variants) were prepared as shown in Table Method 2. Reactions were started by the addition of sodium dithionite and were quenched after 1, 2, 3, 6, 18, or 20 hr. The reaction was added to a 0.5-mL Amicon Ultra Centrifugal Filter 3 kDa (Merck) and centrifuged at 13400 rpm for 10 min. The resulting flow-through was then carefully transferred to an HPLC low-volume (300 μL) recovery vial (Fisher Scientific) and sealed tightly with a polypropylene cap (Fisher Scientific). The resulting progress curves were fitted using a slow-binding inhibition model (Equation 3):

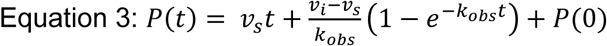

In which:

P(t): product formed at time t

*v_s_*: steady-state rate

*v_i_*: initial rate

*k_obs_*: observed rate constant

P(0): the amount of product at time zero

T: time

### Chain-termination activity measurements inside cells (VITAS)

The chain-termination activity was analyzed in cells using the Viral polymerase-Inhibition Toxin-Associated Selection (VITAS) assay ^49^. Each pBAD/His-C plasmid containing a SAND enzyme was transformed together with a pET28a plasmid carrying the toxin protein LdrD gene (pET-28) in BL21-AI cells. The dead control consists of cells transformed only with a plasmid expressing the LdrD toxin. After transformation, 15 µl of 20% L-(+)-Arabinose (Sigma-Aldrich, 98%) stock was added to each 300µl transformation reaction. Subsequently, 180µl of the transformation reaction was spread on LB-agar plates containing the suitable antibiotics: for cells transformed with hSAND and LdrD, the antibiotics were ampicillin (100 µg/ml) and kanamycin (50 µg/ml), and for cells transformed with LdrD alone, the antibiotic was kanamycin (50 µg/ml). The plates were incubated overnight in an incubator at 37°C. The next day, for each transformation, a single colony was picked and inoculated into 1 mL LB medium including 0.5 µl 20% L-(+)-Arabinose (Sigma-Aldrich, 98%) stock. The samples were then incubated in a shaker/incubator (200 RPM, 37°C) for 6 hours. Bacterial growth associated with each transformation condition was measured by recording the optical density (OD) at 600 nm. LB medium was used as a blank. The cell growth was corrected for the intracellular level of each SAND as measured using western blot (Figures S24-25) and described previously.^22^

### Western blot analysis

Protein samples were extracted from cells using lysis buffer supplemented with protease and phosphatase inhibitors. 12% SDS-PAGE gels were prepared using the SureCast Gel Handcast System (Invitrogen), according to the manufacturer’s protocol. 3-5 µl of PageRuler™ Plus Pre-stained (ThermoFisher) was used as a ladder. 10 µl of sample or control was loaded into the gel. Following electrophoresis, the proteins were transferred to a PVDF Membrane (ThermoFisher) using the Power Blotter system (ThermoFisher). Antibody incubation was performed using the iBind Flex system (ThermoFisher). The antibodies were HRP-conjugated 6x-His Tag Monoclonal Antibody (Invitrogen, MA1-21315-HRP) (1:1000), GAPDH HRP-conjugated monoclonal antibody (GA1R) (Invitrogen, MA5-15738-HRP) (1:1000). The Clarity Western ECL Substrate was used for staining. 5 mL of chemiluminescent substrate was applied to each membrane for 3 minutes. The membrane was imaged using the iBright imaging system (ThermoFisher), and quantification was performed with iBright software.

### Preparation of metabolomic samples using HEK293 cells

HEK293T cells were transfected using Lipofectamine 3000 Transfection Kit (Thermo Fisher Scientific) as described before ^22^. Cells were transfected with the hSAND (RSAD2) plasmid, the CYB5R3 plasmid, or both CYB5R3 and hSAND plasmids ^22^. Transfection efficiency was monitored using GFP and was circa 60-70%. 48 hours after transfection, metabolites were extracted as described previously ^22,39^. DNA concentrations of samples were measured and normalized to the lowest DNA concentration using an aliquot of 100% methanol. Next, 350 µl of each sample was added to a 0.5 ml Amicon Ultra Centrifugal Filter 3 kDa (Merck) and centrifuged at 16250 x g for 30 minutes at room temperature. The flow-through was transferred to a recovery HPLC vial (Fisher Scientific) and sealed tightly with a Polypropylene cap (Fisher Scientific). The samples were stored at -80°C and later analyzed by HR LC-MS.

### High-resolution LC-MS

Samples were analyzed using an Agilent 6546 LC/Q-TOF coupled to a 1290 Infinity II Bio LC (Agilent Technologies, Santa Clara, CA) with an Agilent Jet Stream electrospray ionization source (AJS). The instrument was run in negative-ion mode with an m/z range of 100-1100 and a scan rate of 1 spectrum per second in MS1 mode. Source parameters were as follows; gas temperature, 225°C; gas flow, 9 L/min; Nebulizer, 30 psig; sheath gas temperature, 375 °C, sheath gas flow 12 L/min; capillary voltage, 3 kV; nozzle voltage 500 V and fragmentor voltage 175 V. For each run, 5 µl of sample was injected with a flow rate of 0.2 mL/min and an oven temperature of 45 °C:

- Buffer A: 90 vol.% MeCN (Fisher, 99.9%): 10 vol.% LC-MS grade water (Fisher), 20 mM ammonium acetate, pH 7.4-7.5.
- Buffer B: 20 mM ammonium acetate prepared using LC-MS grade water, pH 7.4-7.5.
- Column: Agilent ZORBAX RR HILIC Plus Column, 2.1 x 100 mm, 3.5 µm
- Flow rate: 0.2 mL/min
- Gradient: [0-1 min]: 100:0 (A:B); [1-10 min] 100:0 (A:B) linearly changed to 10:90 (A:B); [10-15 min] 10:90 (A:B); [15:17] linearly changed to 100:0 (A:B) and [17-35 min] 100-0 (A:B).

Analysis of data was performed using Agilent MassHunter Quantitative Analysis Version 10.

### Molecular Dynamics Simulations of hSAND variants

Three substrate-free systems were simulated: hSAND wild-type (residues 68–361), hSAND-3C (Δ359–361), and hSAND-9C (Δ353–361). All models contained the [4Fe4S]^1+^ (SF4) and radical S-adenosylmethionine (SAM) cofactors. Consistent with the experimentally expressed construct, the N-terminal region (residues 1–67), which had been removed for protein purification, was omitted from all simulation models. Starting coordinates were derived from the hSAND AlphaFold v3 ^52^ model (average pLDDT= 96.11) (Figure S13); truncation variants were prepared by deletion of the corresponding C-terminal residues prior to system building.

Each system was parametrized with the CHARMM36 force field using custom parameters for the [4Fe–4S] cluster (SF4), SAM, and the Fe–Cys coordination patch, ^53,54^ solvated in a TIP3P water box with 15 Å padding and 0.15 M NaCl, and energy-minimized in three stages with progressively released positional restraints using NAMD 3.0b2 ^55,56^. Equilibration consisted of a 2 ns NVT heating phase (1–298 K) followed by 12 ns of NPT equilibration (298 K, 1 bar) with stepwise reduction of positional restraints. Finally, three independent 200 ns production simulations were performed for each system in the NPT ensemble at 298 K and 1 bar.

A 2 fs integration time step was employed by constraining all covalent bonds involving hydrogen atoms using the SHAKE algorithm ^57^. Short-range electrostatic and van der Waals interactions were evaluated using a 12 Å cutoff, with a switching function applied between 10 and 12 Å to ensure a smooth decay of the interaction potential. Long-range electrostatic interactions were treated using the particle mesh Ewald (PME) method ^58^. Temperature and pressure were maintained at 298 K and 1 bar, respectively, using Langevin piston dynamics ^59^.

All apo simulations were analyzed over a 15-200 ns production window. Trajectories were subsampled every 10 frames, corresponding to 100 ps intervals. The conformational ensembles sampled by the three protein variants are shown in Figure S13. Protein dynamics were first characterized by calculating the per-residue Cα root-mean-square fluctuations (Cα-RMSF), with particular attention to the phosphate-coordinating residues Arg346 and Tyr350 (Figure S14). The average Cα-RMSF of Arg346 was 1.15 ± 0.40 Å, 4.10 ± 1.97 Å, and 2.64 ± 2.22 Å for WT hSAND, hSAND-3C, and hSAND-9C, respectively. Similarly, Tyr350 exhibited average Cα-RMSF values of 1.87 ± 2.32 Å, 5.18 ± 10.88 Å, and 2.52 ± 1.44 Å for WT hSAND, hSAND-3C, and hSAND-9C, respectively. These results indicate substantially increased mobility of the C-terminal phosphate-coordinating residues following partial C-terminal truncation, particularly in the hSAND-3C variant. In contrast, the remaining phosphate-coordinating residues (Lys119, Lys219, and Arg244), previously identified from crystal structures of viperin bound to nucleotide substrates (Fenwick *et al*., 2017, PDB 5VSL; Patel *et al*., 2020, PDB 6Q2Q), remained highly rigid throughout the simulations, exhibiting average Cα-RMSF values below 0.63 Å.

To identify regions of the protein core that interact with the C-tail, Cα–Cα contact occupancies were calculated between C-tail residues 340–361 and core residues 68–339 using three independent MD simulations of WT hSAND (Figure S14). The highest contact occupancies were observed between Leu359 of the C-tail and the core residues Arg212 (0.999 ± 0.004) and Arg215 (0.956 ± 0.037). Owing to the high persistence of these interactions, Arg212 and Arg215 were defined as the principal anchoring residues for C-tail binding and were used as reference residues in the subsequent analyses.

To quantify the effect of C-terminal truncation on C-tail anchoring, we measured the distance between Arg212 Cα and the Cα atom of the terminal C-tail residue in the apo simulations. For WT hSAND, the terminal residue is Trp361, whereas the corresponding terminal residues are Lys358 in hSAND-3C (Δ359–361) and Trp352 in hSAND-9C (Δ353–361). The distance between the terminal C-tail residue and Arg212 provides a measure of the maximum reach of the C-tail toward the anchoring pocket. If the terminal residue cannot approach Arg212, the C-tail is geometrically excluded from forming the native anchoring interactions.

## Ancestral sequence reconstruction (ASR)

To perform ASR, the amino acid sequence of hSAND (UniProt ID: Q8WXG1) was used. ASR was performed using FireProt server.^60,61^ The hotspot amino acid residues identified around the LDW tripeptide were then visualized on the modeled structure of hSAND.

## Data and materials availability

All data supporting the findings of this study are available in the main text or supplementary data. Source data for all figures are available from the corresponding author upon reasonable request. The use of plasmids expressing variants will be limited to non-commercial use, and the plasmids will be provided upon signing a material transfer agreement (MTA).

## Acknowledgments

KHE acknowledges the generous gift of an anaerobic glovebox from Prof Annalisa Pastore. KHE and SCB acknowledge support from the Royal Society for an International Exchanges Grant (IES\R1\221117). KHE acknowledges support from the Royal Society for a Research Grant (RGS\R1\231135) and the Medical Research Council, MRC Gap Fund (MR/Y50337X/1). MW is supported by CSC-KCL Doctoral Scholarship (202408410098). MAM acknowledges support from the German Research Foundation (DFG) under Germany’s Excellence Strategy – EXC 2008-390540038 – UniSysCat. All authors acknowledge support from European Cooperation in Science and Technology (CA21115).

## Author contributions

Conceptualization: MW, KHE; Data collection: MW, DP, HF; Data curation: MW, DP, HF, MAM, KHE; Methodology: MW, DP, MAM, KHE; Investigation: MW, DP, MAM, KHE; Analysis and discussion: MW, DP, WSJ, MAM, KHE; Visualization: MW, DP, MAM, KHA; Funding acquisition: MW, KHE, MAM; Project administration: KHE; Supervision: MAM, KHE; Writing – original draft: WSJ, MW, KHE; Writing – review & editing: MW, DP, SCB, WSJ, MAM, KHE

## Competing interests

Authors declare that they have no competing interests.

## Notes

### Competing Interest Statement

The authors have declared no competing interest.

### Summary of Updates

Textual modification to clarify the message and some additional data to further support the conclusions.

