## Supplementary File for "An intrinsically disordered region controls viperin (SAND) antiviral synthesis via a clamp-and-latch mechanism"

### Supplementary Methods

**Chemical**

All chemicals were reagent grade and purchased from ThemoFisher Scientific (Waltham, MA, USA), Fisher Scientific, (Loughborough, UK) or Merck, (Rahway, NJ, USA). All anaerobic buffers were prepared a day prior to the experiments and stored in the glovebox overnight to ensure complete removal of dioxygen.

**Genes and cloning**

The *E. coli* codon-optimized synthetic genes for different enzymes were obtained from GeneArt (ThermoFisher) either in pMQ plasmid or final expression vector, pBAD/His C. Cloning from the pMQ vector to the pBAD/His C vector was performed using 5′ and 3′ restriction sites for KpnI and HindIII, respectively. The presence of the correct insert was confirmed using restriction enzyme double digestion and DNA sequencing (GENEWIZ). The plasmids were transformed into *E. Coli* TOP10 cells, and a glycerol stock of the cells was prepared and stored at −80 °C. The glycerol stock was used to prepare LB Agar (100 μm ampicillin) plates for protein expression. All cloned enzymes lacked the N-terminal hydrophobic domain. Residue numbering is based on the full-length proteins.

**TtSAND-18C**
MPVSVNYHFTRQCNKSCGFCFHTATTSHIENLERQQAALRLLAQAGMRKINFAGGEPFLYPKA
LGAMIDFCKRELRLESVSVVTNGSLVREDFLRRHAGQLDILAVSCDSFNEQTNVAIGRGSGDQ
VPRLYQIAEWCRTYGVLFKINTVVNRLNVAEDMNAHIAALRPFRWKCFQVLIVRGENDSDETLR
NAHRFTISDEEFEAFCERHRGQACLVPEPNRLMAKSYLILDEYLRFLDRTGKQPSKPILEVGVE
KALESVYWDEEAFVERGGLYEW

**TtSAND-6C**
MPVSVNYHFTRQCNKSCGFCFHTATTSHIENLERQQAALRLLAQAGMRKINFAGGEPFLYPKA
LGAMIDFCKRELRLESVSVVTNGSLVREDFLRRHAGQLDILAVSCDSFNEQTNVAIGRGSGDQ
VPRLYQIAEWCRTYGVLFKINTVVNRLNVAEDMNAHIAALRPFRWKCFQVLIVRGENDSDETLR
NAHRFTISDEEFEAFCERHRGQACLVPEPNRLMAKSYLILDEYLRFLDRTGKQPSKPILEVGVE
KALESVYWDEEAFVERGGLYEWTKPANGESGCGS

**TtSAND-12C**
MPVSVNYHFTRQCNKSCGFCFHTATTSHIENLERQQAALRLLAQAGMRKINFAGGEPFLYPKA
LGAMIDFCKRELRLESVSVVTNGSLVREDFLRRHAGQLDILAVSCDSFNEQTNVAIGRGSGDQ
VPRLYQIAEWCRTYGVLFKINTVVNRLNVAEDMNAHIAALRPFRWKCFQVLIVRGENDSDETLR
NAHRFTISDEEFEAFCERHRGQACLVPEPNRLMAKSYLILDEYLRFLDRTGKQPSKPILEVGVE
KALESVYWDEEAFVERGGLYEWTKPANG

**TtSAND-15C**
MPVSVNYHFTRQCNKSCGFCFHTATTSHIENLERQQAALRLLAQAGMRKINFAGGEPFLYPKA
LGAMIDFCKRELRLESVSVVTNGSLVREDFLRRHAGQLDILAVSCDSFNEQTNVAIGRGSGDQ
VPRLYQIAEWCRTYGVLFKINTVVNRLNVAEDMNAHIAALRPFRWKCFQVLIVRGENDSDETLR
NAHRFTISDEEFEAFCERHRGQACLVPEPNRLMAKSYLILDEYLRFLDRTGKQPSKPILEVGVE
KALESVYWDEEAFVERGGLYEWTKP

**TtSAND-SUMO**

MPVSVNYHFTRQCNKSCGFCFHTATTSHIENLERQQAALRLLAQAGMRKINFAGGEPFLYPKA
LGAMIDFCKRELRLESVSVVTNGSLVREDFLRRHAGQLDILAVSCDSFNEQTNVAIGRGSGDQ
VPRLYQIAEWCRTYGVLFKINTVVNRLNVAEDMNAHIAALRPFRWKCFQVLIVRGENDSDETLR
NAHRFTISDEEFEAFCERHRGQACLVPEPNRLMAKSYLILDEYLRFLDRTGKQPSKPILEVGVE
KALESVYWDEEAFVERGGLYEWTKPANGESGCGSKKELEWMHHHHHHGSLQDSEVNQEAK
PEVKPEVKPETHINLKVSDGSSEIFFKIKKTTPLRRLMEAFAKRQGKEMDSLTFLYDGIEIQADQT
PEDLDMEDNDIIEAHREQIGGATY

**TtSAND_RYA (Arg295Ala and Tyr299Ala)**
MPVSVNYHFTRQCNKSCGFCFHTATTSHIENLERQQAALRLLAQAGMRKINFAGGEPFLYPKA
LGAMIDFCKRELRLESVSVVTNGSLVREDFLRRHAGQLDILAVSCDSFNEQTNVAIGRGSGDQ
VPRLYQIAEWCRTYGVLFKINTVVNRLNVAEDMNAHIAALRPFRWKCFQVLIVRGENDSDETLR
NAHRFTISDEEFEAFCERHRGQACLVPEPNRLMAKSYLILDEYLRFLDRTGKQPSKPILEVGVE
KALESVYWDEEAFVEAGGLAEWTKPANGESGCGSKKELEW

**TtSAND-18C_RYA (Arg295Ala and Tyr299Ala)**
MPVSVNYHFTRQCNKSCGFCFHTATTSHIENLERQQAALRLLAQAGMRKINFAGGEPFLYPKA
LGAMIDFCKRELRLESVSVVTNGSLVREDFLRRHAGQLDILAVSCDSFNEQTNVAIGRGSGDQ
VPRLYQIAEWCRTYGVLFKINTVVNRLNVAEDMNAHIAALRPFRWKCFQVLIVRGENDSDETLR
NAHRFTISDEEFEAFCERHRGQACLVPEPNRLMAKSYLILDEYLRFLDRTGKQPSKPILEVGVE
KALESVYWDEEAFVEAGGLAEW

**TtSAND_TKP-AAA**
MPVSVNYHFTRQCNKSCGFCFHTATTSHIENLERQQAALRLLAQAGMRKINFAGGEPFLYPKA
LGAMIDFCKRELRLESVSVVTNGSLVREDFLRRHAGQLDILAVSCDSFNEQTNVAIGRGSGDQ
VPRLYQIAEWCRTYGVLFKINTVVNRLNVAEDMNAHIAALRPFRWKCFQVLIVRGENDSDETLR
NAHRFTISDEEFEAFCERHRGQACLVPEPNRLMAKSYLILDEYLRFLDRTGKQPSKPILEVGVE
KALESVYWDEEAFVERGGLYEWAAAANGESGCGSKKELEW

**TtSAND-TA**
MPVSVNYHFTRQCNKSCGFCFHTATTSHIENLERQQAALRLLAQAGMRKINFAGGEPFLYPKA
LGAMIDFCKRELRLESVSVVTNGSLVREDFLRRHAGQLDILAVSCDSFNEQTNVAIGRGSGDQ
VPRLYQIAEWCRTYGVLFKINTVVNRLNVAEDMNAHIAALRPFRWKCFQVLIVRGENDSDETLR
NAHRFTISDEEFEAFCERHRGQACLVPEPNRLMAKSYLILDEYLRFLDRTGKQPSKPILEVGVE
KALESVYWDEEAFVERGGLYEWAKPANGESGCGSKKELEW

**TtSAND-KA**
MPVSVNYHFTRQCNKSCGFCFHTATTSHIENLERQQAALRLLAQAGMRKINFAGGEPFLYPKA
LGAMIDFCKRELRLESVSVVTNGSLVREDFLRRHAGQLDILAVSCDSFNEQTNVAIGRGSGDQ
VPRLYQIAEWCRTYGVLFKINTVVNRLNVAEDMNAHIAALRPFRWKCFQVLIVRGENDSDETLR
NAHRFTISDEEFEAFCERHRGQACLVPEPNRLMAKSYLILDEYLRFLDRTGKQPSKPILEVGVE
KALESVYWDEEAFVERGGLYEWTAPANGESGCGSKKELEW

**hSAND-3C**
MPTTPTSVNYHFTRQCNYKCGFCFHTAKTSFVLPLEEAKRGLLLLKEAGMEKINFSGGEPFLQ
DRGEYLGKLVRFCKVELRLPSVSIVSNGSLIRERWFQNYGEYLDILAISCDSFDEEVNVLIGRGQ
GKKNHVENLQKLRRWCRDYRVAFKINSVINRFNVEEDMTEQIKALNPVRWKVFQCLLIEGENC
GEDALREAERFVIGDEEFERFLERHKEVSCLVPESNQKMKDSYLILDEYMRFLNCRKGRKDPS
KSILDVGVEEAIKFSGFDEKMFLKRGGKYIWSKADLK

**hSAND-9C**
MPTTPTSVNYHFTRQCNYKCGFCFHTAKTSFVLPLEEAKRGLLLLKEAGMEKINFSGGEPFLQ
DRGEYLGKLVRFCKVELRLPSVSIVSNGSLIRERWFQNYGEYLDILAISCDSFDEEVNVLIGRGQ
GKKNHVENLQKLRRWCRDYRVAFKINSVINRFNVEEDMTEQIKALNPVRWKVFQCLLIEGENC
GEDALREAERFVIGDEEFERFLERHKEVSCLVPESNQKMKDSYLILDEYMRFLNCRKGRKDPS
KSILDVGVEEAIKFSGFDEKMFLKRGGKYIW

**hSAND-9C+18TtC**
MPTTPTSVNYHFTRQCNYKCGFCFHTAKTSFVLPLEEAKRGLLLLKEAGMEKINFSGGEPFLQ
DRGEYLGKLVRFCKVELRLPSVSIVSNGSLIRERWFQNYGEYLDILAISCDSFDEEVNVLIGRGQ
GKKNHVENLQKLRRWCRDYRVAFKINSVINRFNVEEDMTEQIKALNPVRWKVFQCLLIEGENC
GEDALREAERFVIGDEEFERFLERHKEVSCLVPESNQKMKDSYLILDEYMRFLNCRKGRKDPS
KSILDVGVEEAIKFSGFDEKMFLKRGGKYIWTKPANGESGCGSKKELEW

**hSAND+18TtC**
MPTTPTSVNYHFTRQCNYKCGFCFHTAKTSFVLPLEEAKRGLLLLKEAGMEKINFSGGEPFLQ
DRGEYLGKLVRFCKVELRLPSVSIVSNGSLIRERWFQNYGEYLDILAISCDSFDEEVNVLIGRGQ
GKKNHVENLQKLRRWCRDYRVAFKINSVINRFNVEEDMTEQIKALNPVRWKVFQCLLIEGENC
GEDALREAERFVIGDEEFERFLERHKEVSCLVPESNQKMKDSYLILDEYMRFLNCRKGRKDPS
KSILDVGVEEAIKFSGFDEKMFLKRGGKYIWSKADLKLDWTKPANGESGCGSKKELEW

**hSAND+12TtC**
MPTTPTSVNYHFTRQCNYKCGFCFHTAKTSFVLPLEEAKRGLLLLKEAGMEKINFSGGEPFLQ
DRGEYLGKLVRFCKVELRLPSVSIVSNGSLIRERWFQNYGEYLDILAISCDSFDEEVNVLIGRGQ
GKKNHVENLQKLRRWCRDYRVAFKINSVINRFNVEEDMTEQIKALNPVRWKVFQCLLIEGENC
GEDALREAERFVIGDEEFERFLERHKEVSCLVPESNQKMKDSYLILDEYMRFLNCRKGRKDPS
KSILDVGVEEAIKFSGFDEKMFLKRGGKYIWSKADLKLDWTKPANGESGCGS

**hSAND_RYA (Arg346Ala & Tyr350Ala)**
MPTTPTSVNYHFTRQCNYKCGFCFHTAKTSFVLPLEEAKRGLLLLKEAGMEKINFSGGEPFLQ
DRGEYLGKLVRFCKVELRLPSVSIVSNGSLIRERWFQNYGEYLDILAISCDSFDEEVNVLIGRGQ
GKKNHVENLQKLRRWCRDYRVAFKINSVINRFNVEEDMTEQIKALNPVRWKVFQCLLIEGENC
GEDALREAERFVIGDEEFERFLERHKEVSCLVPESNQKMKDSYLILDEYMRFLNCRKGRKDPS
KSILDVGVEEAIKFSGFDEKMFLKAGGKAIWSKADLKLDW

**hSAND-9C_RYA (Arg346Ala & Tyr350Ala)**
MPTTPTSVNYHFTRQCNYKCGFCFHTAKTSFVLPLEEAKRGLLLLKEAGMEKINFSGGEPFLQ
DRGEYLGKLVRFCKVELRLPSVSIVSNGSLIRERWFQNYGEYLDILAISCDSFDEEVNVLIGRGQ
GKKNHVENLQKLRRWCRDYRVAFKINSVINRFNVEEDMTEQIKALNPVRWKVFQCLLIEGENC
GEDALREAERFVIGDEEFERFLERHKEVSCLVPESNQKMKDSYLILDEYMRFLNCRKGRKDPS
KSILDVGVEEAIKFSGFDEKMFLKAGGKAIW

**hSAND_RKNS (R215K and N241S)**
MPLPTTPTSVNYHFTRQCNYKCGFCFHTAKTSFVLPLEEAKRGLLLLKEAGMEKINFSGGEPFL
QDRGEYLGKLVRFCKVELRLPSVSIVSNGSLIRERWFQNYGEYLDILAISCDSFDEEVNVLIGRG
QGKKNHVENLQKLRRWCRDYKVAFKINSVINRFNVEEDMTEQIKALSPVRWKVFQCLLIEGEN
CGEDALREAERFVIGDEEFERFLERHKEVSCLVPESNQKMKDSYLILDEYMRFLNCRKGRKDP
SKSILDVGVEEAIKFSGFDEKMFLKRGGKYIWSKADLKLDW

**hSAND_RW (R215W)**

MPLPTTPTSVNYHFTRQCNYKCGFCFHTAKTSFVLPLEEAKRGLLLLKEAGMEKINFSGGEPFL
QDRGEYLGKLVRFCKVELRLPSVSIVSNGSLIRERWFQNYGEYLDILAISCDSFDEEVNVLIGRG
QGKKNHVENLQKLRRWCRDYWVAFKINSVINRFNVEEDMTEQIKALNPVRWKVFQCLLIEGEN
CGEDALREAERFVIGDEEFERFLERHKEVSCLVPESNQKMKDSYLILDEYMRFLNCRKGRKDP
SKSILDVGVEEAIKFSGFDEKMFLKRGGKYIWSKADLKLDW

**hSAND_WA (W361A)**
MPTTPTSVNYHFTRQCNYKCGFCFHTAKTSFVLPLEEAKRGLLLLKEAGMEKINFSGGEPFLQ
DRGEYLGKLVRFCKVELRLPSVSIVSNGSLIRERWFQNYGEYLDILAISCDSFDEEVNVLIGRGQ
GKKNHVENLQKLRRWCRDYRVAFKINSVINRFNVEEDMTEQIKALNPVRWKVFQCLLIEGENC
GEDALREAERFVIGDEEFERFLERHKEVSCLVPESNQKMKDSYLILDEYMRFLNCRKGRKDPS
KSILDVGVEEAIKFSGFDEKMFLKRGGKYIWSKADLKLDA

**hSAND_DA (D360A)**
MPTTPTSVNYHFTRQCNYKCGFCFHTAKTSFVLPLEEAKRGLLLLKEAGMEKINFSGGEPFLQ
DRGEYLGKLVRFCKVELRLPSVSIVSNGSLIRERWFQNYGEYLDILAISCDSFDEEVNVLIGRGQ
GKKNHVENLQKLRRWCRDYRVAFKINSVINRFNVEEDMTEQIKALNPVRWKVFQCLLIEGENC
GEDALREAERFVIGDEEFERFLERHKEVSCLVPESNQKMKDSYLILDEYMRFLNCRKGRKDPS
KSILDVGVEEAIKFSGFDEKMFLKRGGKYIWSKADLKLAW

**hSAND_LA (L359A)**
MPTTPTSVNYHFTRQCNYKCGFCFHTAKTSFVLPLEEAKRGLLLLKEAGMEKINFSGGEPFLQ
DRGEYLGKLVRFCKVELRLPSVSIVSNGSLIRERWFQNYGEYLDILAISCDSFDEEVNVLIGRGQ
GKKNHVENLQKLRRWCRDYRVAFKINSVINRFNVEEDMTEQIKALNPVRWKVFQCLLIEGENC
GEDALREAERFVIGDEEFERFLERHKEVSCLVPESNQKMKDSYLILDEYMRFLNCRKGRKDPS
KSILDVGVEEAIKFSGFDEKMFLKRGGKYIWSKADLKADW

**Rat SAND (rSAND)**
MQPTTPVSVNYHFTRQCNYKCGFCFHTAKTSFVLPLEEAKRGLLLLKQAGMEKINFSGGEPFL
QDRGEYLGKLVRFCKEELALPSVSIVSNGSLIRERWFKDYGDYLDILAISCDSFDEQVNVLIGRG
QGKKNHVENLQKLRKWCRDYKVAFKINSVINRFNVDEDMNEHIKALSPVRWKVFQCLLIEGEN
SGEDALREAERFLISNEEFEAFLQRHKDVSCLVPESNQKMKDSYLILDEYMRFLNCTGGRKDP
SRSILDVGVEEAIKFSGFDEKMFLKRGGKYVWSKADLKLDW

**rSAND-3C**
MQPTTPVSVNYHFTRQCNYKCGFCFHTAKTSFVLPLEEAKRGLLLLKQAGMEKINFSGGEPFL
QDRGEYLGKLVRFCKEELALPSVSIVSNGSLIRERWFKDYGDYLDILAISCDSFDEQVNVLIGRG
QGKKNHVENLQKLRKWCRDYKVAFKINSVINRFNVDEDMNEHIKALSPVRWKVFQCLLIEGEN
SGEDALREAERFLISNEEFEAFLQRHKDVSCLVPESNQKMKDSYLILDEYMRFLNCTGGRKDP
SRSILDVGVEEAIKFSGFDEKMFLKRGGKYVWSKADLK

**Table method 1.** The samples were prepared to create the standard curve.

| Standard | iron solution stock (µL) | MiniQ water (µL) | Dilution factor | Concentration (µg/mL) |
|---|---|---|---|---|
| S1 | 1 | 99 | 100 | 1 |
| S2 | 2 | 98 | 50 | 2 |
| S3 | 5 | 95 | 20 | 5 |
| S4 | 10 | 90 | 10 | 10 |
| S5 | 20 | 80 | 5 | 20 |
| S6 | 50 | 50 | 2 | 50 |

Equation 1: $\frac{[4Fe-4S]cluser}{Protein} = \frac{Concentration\ of\ iron\ (\mu M)}{C_{protein} \times purity}$

In which

$c_{protein}$: Concentration of protein as measured by BCA.
Purity: The percentage of purified protein as determined by SDS analysis.

**Table Method 2.** Details of reaction setup using different enzymes for analysis of ddhNTP formation.

| Sample | Enzyme (µL) | 40mM CTP, GTP, ATP, or UTP (µL) | 50mM SAM (µL) | 80mM Sodium Dithionite (µL) | Mops (µL) |
|---|---|---|---|---|---|
| R1 (hSAND-WT) (standard) | 100 | 5 | 10 | 20 | 0 |
| R2 (hSAND variant) | 100 | 5 | 10 | 20 | 0 |
| Cont. (no enzyme) | 0 | 5 | 10 | 20 | 100 |
| R'1 (WT-TtSAND) (standard) | 100 | 5 | 10 | 20 | 0 |
| R'2 (TtSAND mutants) | 100 | 5 | 10 | 20 | 0 |
| Cont. (no enzyme) | 0 | 5 | 10 | 20 | 100 |

**Liquid chromatography-Mass spectrometry**

Measurements were performed using an Agilent InfinityLab G6160A LC/MSD iQ mass spectrometer equipped with an Agilent 1260 Infinity II liquid chromatography system. The column was an Agilent ZORBAX RR HILIC Plus Column, 2.1 x 100 mm, 3.5 µm, or RRHD 2.1x150 mm, 1.8 µm. Instrument control and data processing were performed using OpenLab CDS (Agilent Technologies, Santa Clara, CA, USA). The system was calibrated on the day of the analysis. Electrospray source conditions were adjusted to maximize sensitivity, and the detection mode was set to detect negative (−) ions. For each run, 50 µL of solution was injected with a flow rate of 0.2 mL·min−1 and an oven temperature of 45 °C:

Buffer A: 90 vol.% MeCN (acetonitrile) (Fisher, 99.9%): 10 vol.% LC–MS grade water (Fisher), 20 mM ammonium acetate, pH 7.4–7.5.

Buffer B: 10 vol.% LC–MS-grade water (Fisher), 20 mM ammonium acetate, pH 7.4–7.5.

Column Flow rate: 0.2 mL·min−1.

Gradient: [0–1 min]: 100 : 0 (A : B); [1–10 min] 100 : 0 (A : B) linearly changed to 10 : 90 (A : B); [10–15 min] 10 : 90 (A : B); [15 : 17] linearly changed to 100 : 0 (A : B) and [17–35 min] 100–0 (A : B).

Analysis of the LC–MS data was performed using Open Lab CDS software. Experiments with each variant were performed in parallel to the WT enzyme as a positive control. The areas under the curve were obtained and used to calculate the amount of each ddhNTP using a standard curve. The data were corrected for any variation in protein concentration relative to the WT control (Equation 2).

Equation 2: $ddhNTP\ concentration\ (\mu M) = \left(\frac{(Area-a)}{b}\right) \times \left(\frac{[WT]}{[variant]}\right)$

In which

Area: area under the curve

a: the Y-intercept for the standard curve

b: the slope for the standard curve

[WT]: concentration (µM) of wild-type enzyme

[variant]: concentration (µM) of a variant

**Analysis of enzyme kinetics**

Samples for analysis of the kinetics of TtSAND and hSAND (wild-type and C-terminal truncated variants) were prepared as shown in Table Method 2. Reactions were started by the addition of sodium dithionite and were quenched after 1, 2, 3, 6, 18, or 20 hr. The reaction was added to a 0.5-mL Amicon Ultra Centrifugal Filter 3 kDa (Merck) and centrifuged at 13400 rpm for 10 min. The resulting flow-through was then carefully transferred to an HPLC low-volume (300 µL) recovery vial (Fisher Scientific) and sealed tightly with a polypropylene cap (Fisher Scientific). The resulting progress curves were fitted using a slow-binding inhibition model (Equation 3):

Equation 3: $P(t) = v_s t + \frac{v_i - v_s}{k_{obs}}\left(1 - e^{-k_{obs}t}\right) + P(0)$

In which:
P(t): product formed at time t
$v_s$: steady-state rate
$v_i$: initial rate
$k_{obs}$: observed rate constant
P(0): the amount of product at time zero
T: time

### Supplementary Figures

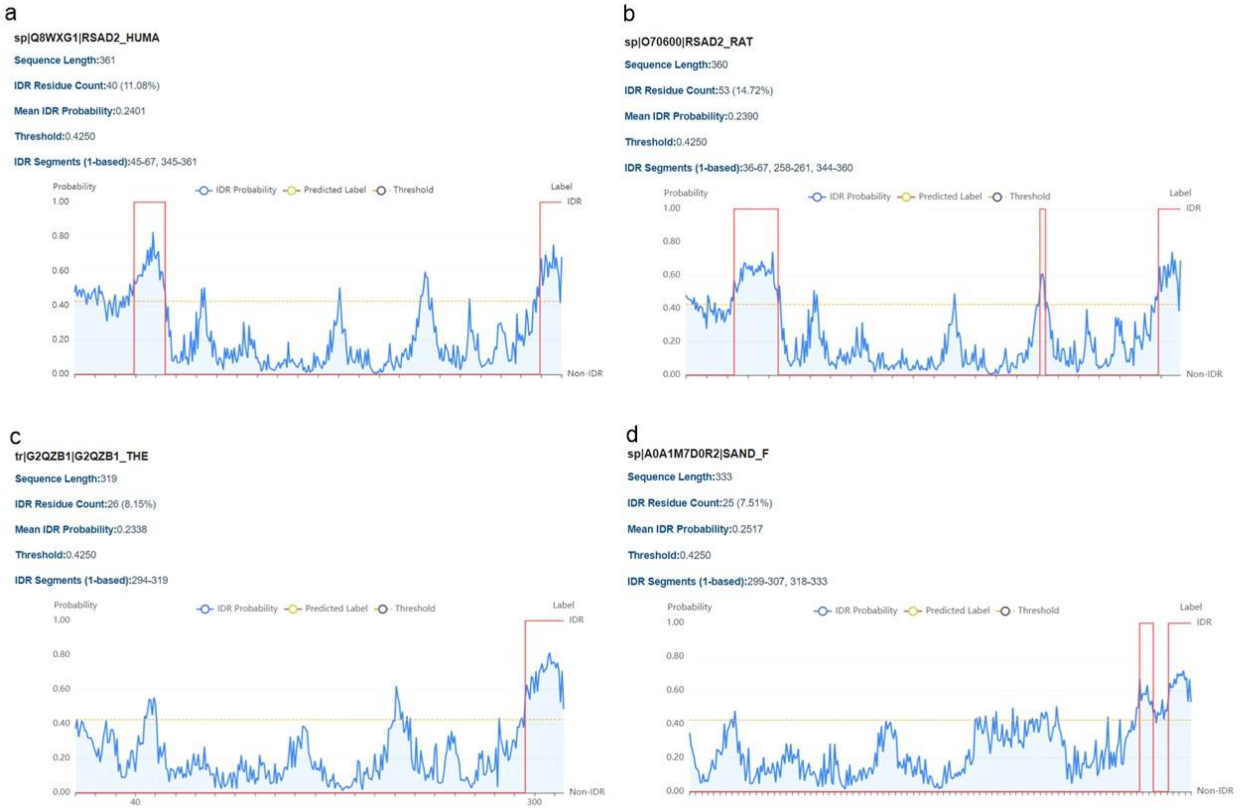


**Figure S1.** The C-terminal domain of SAND from bacteria to humans has a predicted IDR. Prediction of IDRs in representative SAND enzymes from **(a)** human, **(b)** rat, **(c)** fungus *Thielavia terrestris*, and **(d)** bacteria *Fibrobacter sp*. IDRs were predicted using NovoPro Tool. It is built on a protein model trained on approximately 2000 high-quality annotated sequences. It performs residue-level IDR predictions.

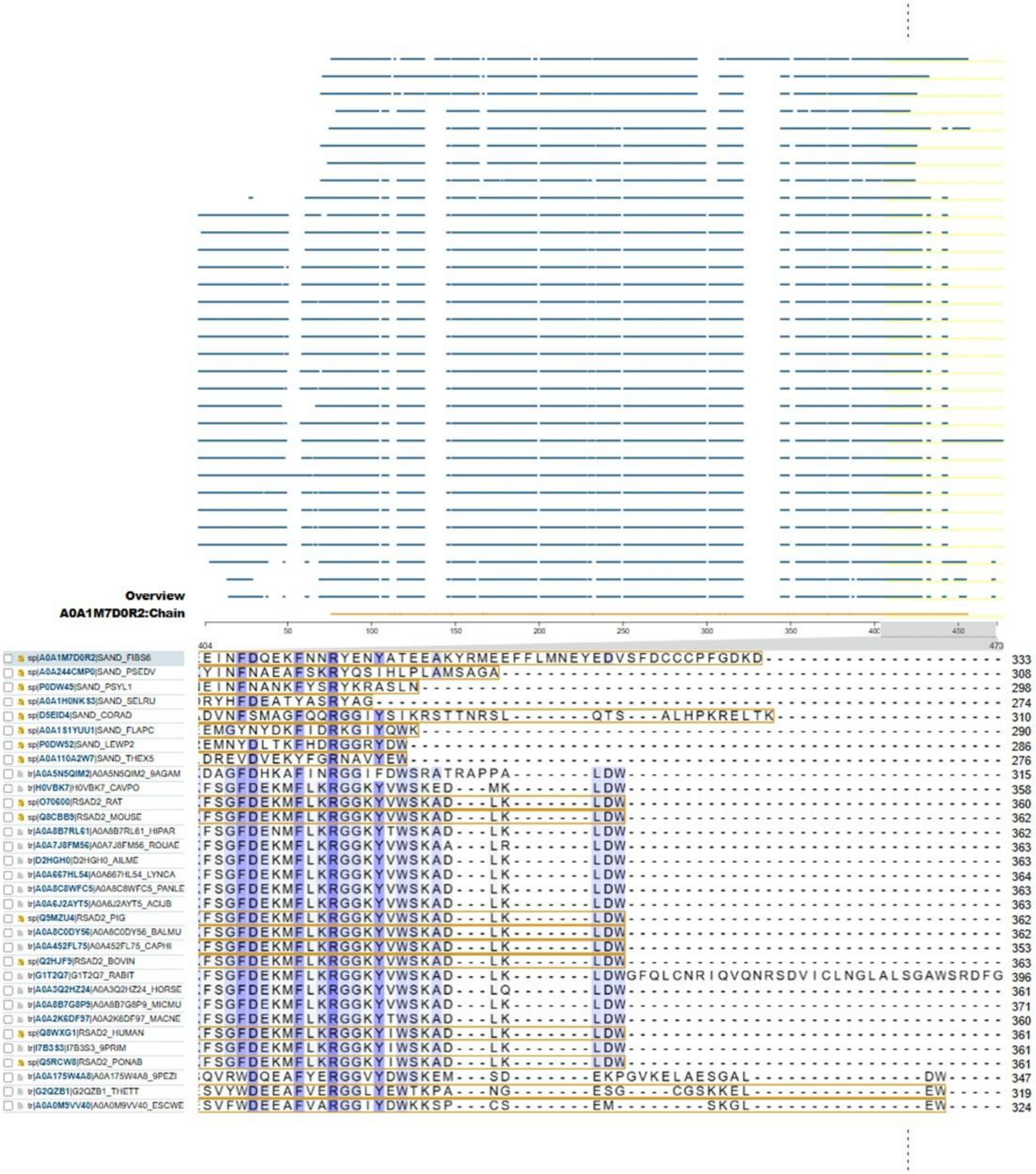

**Figure S2.** Multiple sequence alignment showing the C-terminal variation across SANDs from different species. The length and sequence of the C-terminal IDR are highly variable across bacteria and humans. In some prokaryotic and fungal SANDs, the C-terminal IDR has a tryptophan.

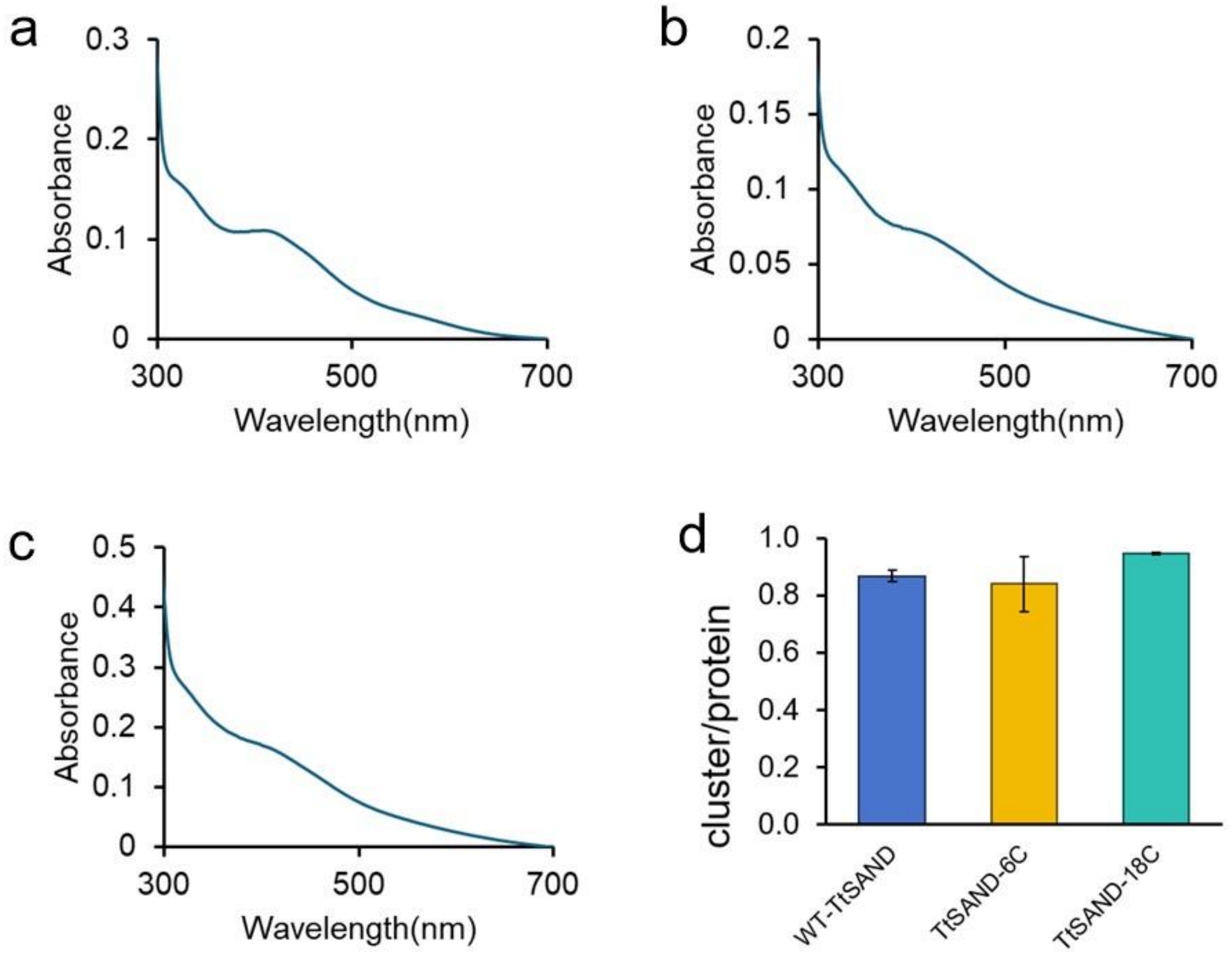


**Figure S3.** Truncation of the C-terminal IDR does not affect the FeS cluster. (a-c) The UV-visible absorbance spectrum of the oxidized $[4Fe\text{-}4S]^{1+}$ cluster in (a) wild-type TtSAND (15.2 μM), (b) TtSAND-6C (8.2 μM), and (c) TtSAND-18C (14.7 μM). The spectra show a peak at around 420 nm characteristic of the oxidized $[4Fe\text{-}4S]^{1+}$ cluster. (d) Quantification of the number of clusters per protein by measuring iron content. Measurements were repeated using at least two different protein batches to confirm reproducibility. Data are the average of two measurements ± errors.

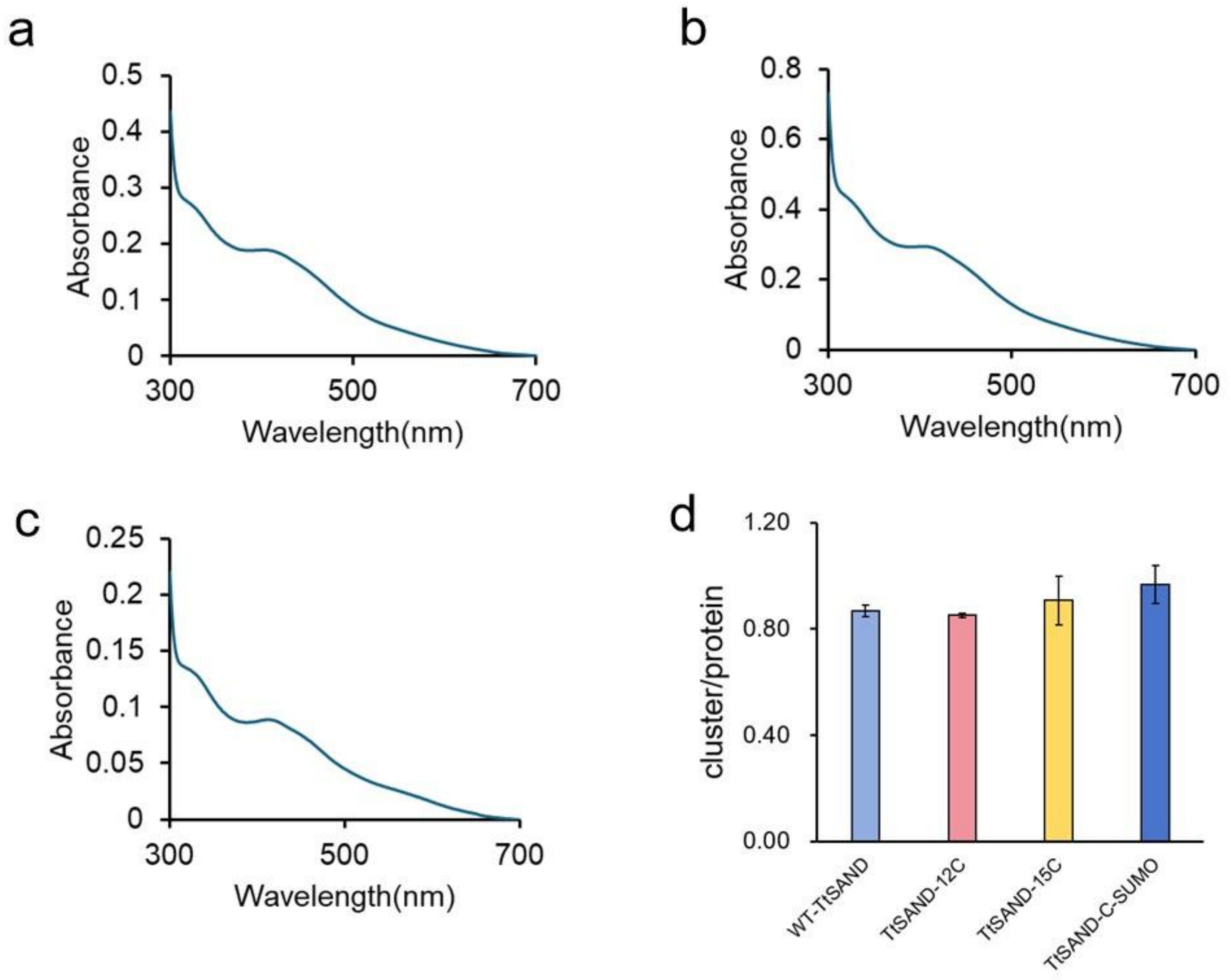


**Figure S4.** Truncation of the C-terminal IDR or addition of SUMO protein to the C-terminus of TtSAND does not affect the FeS cluster. (a-c) The UV-visible absorbance spectrum of the oxidised [4Fe-4S]1+ cluster in (a) TtSAND-12C (24.8 μM), (b) TtSAND-15C (18.1 μM), and (c) TtSAND-C-SUMO (32 μM). The spectra show a peak at around 420 nm characteristic of the oxidised [4Fe-4S]1+ cluster. (d) Quantification of cluster numbers per protein by measuring iron content. Measurements were repeated using at least two different protein batches to confirm reproducibility. Data are the average of two measurements ± errors.

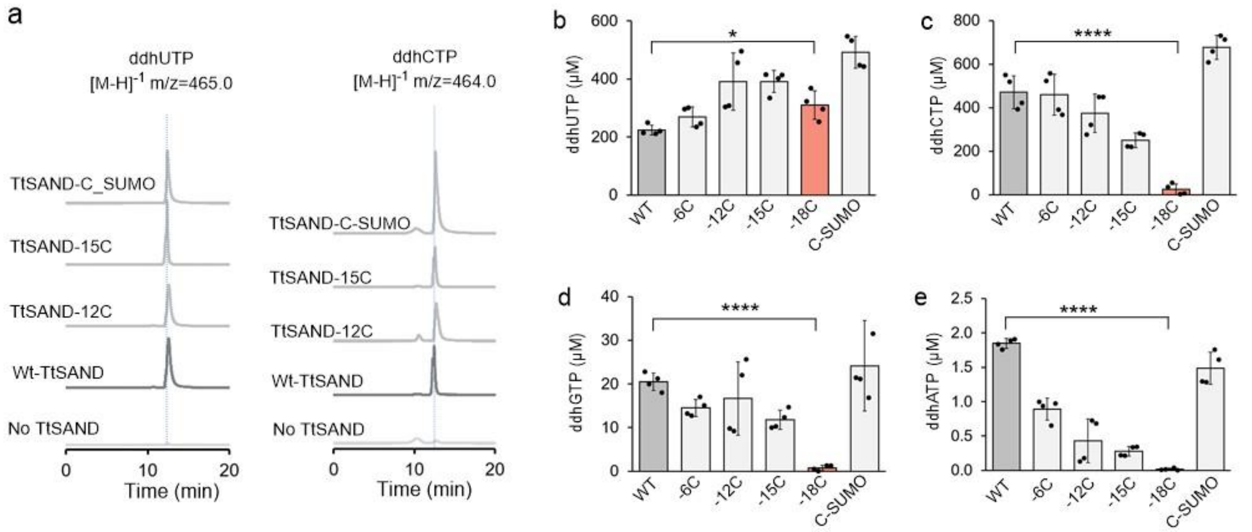


**Figure S5.** When tripeptide TKP is removed, substrate promiscuity in TtSAND is fully abolished. **(a)** Total ion chromatograms of ddhUTP and ddhCTP for WT-TtSAND, TtSAND-12C, TtSAND-6C, and TtSAND-SUMO are shown as an example. **(b-e)** Quantification of the amount of (b) ddhUTP, (c) ddhCTP, (d) ddhGTP, and (e) ATP. The data are averages of four measurements ± standard deviation, obtained from two different protein batches. One-way ANOVA was used to calculate p values. (*p value < 0.05; *****p value < 0.00001).

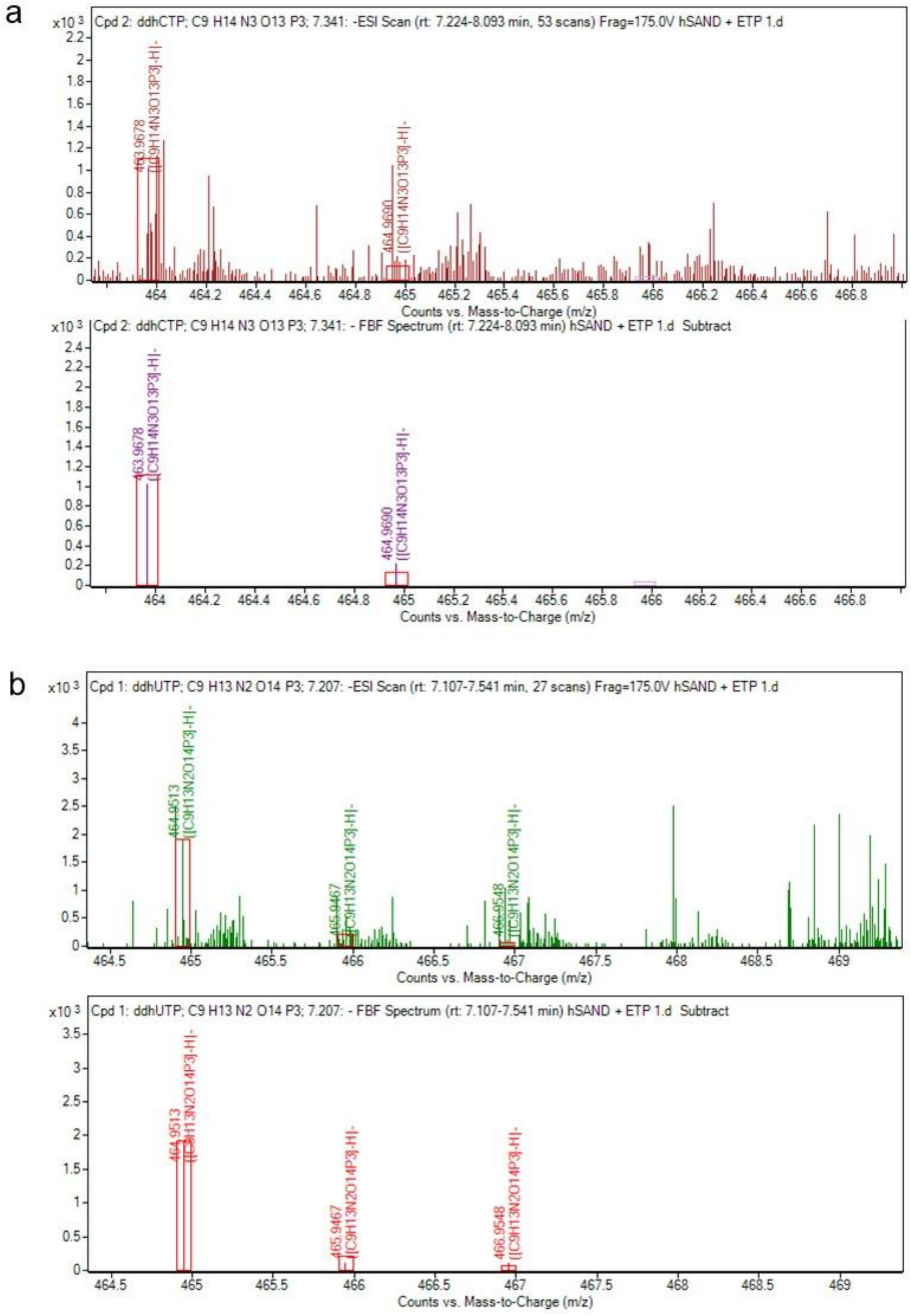


**Figure S6.** High-resolution LC-MS analysis of ddhCTP and ddhUTP formation in HEK293T cells when hSAND and its electron transfer partner protein CYB5R3 are co-expressed. (a) ddhCTP ($[M-H]^{-1}$ m/z=463.9678) and (b) ddhUTP ($[M-H]^{-1}$ m/z=464.9513).

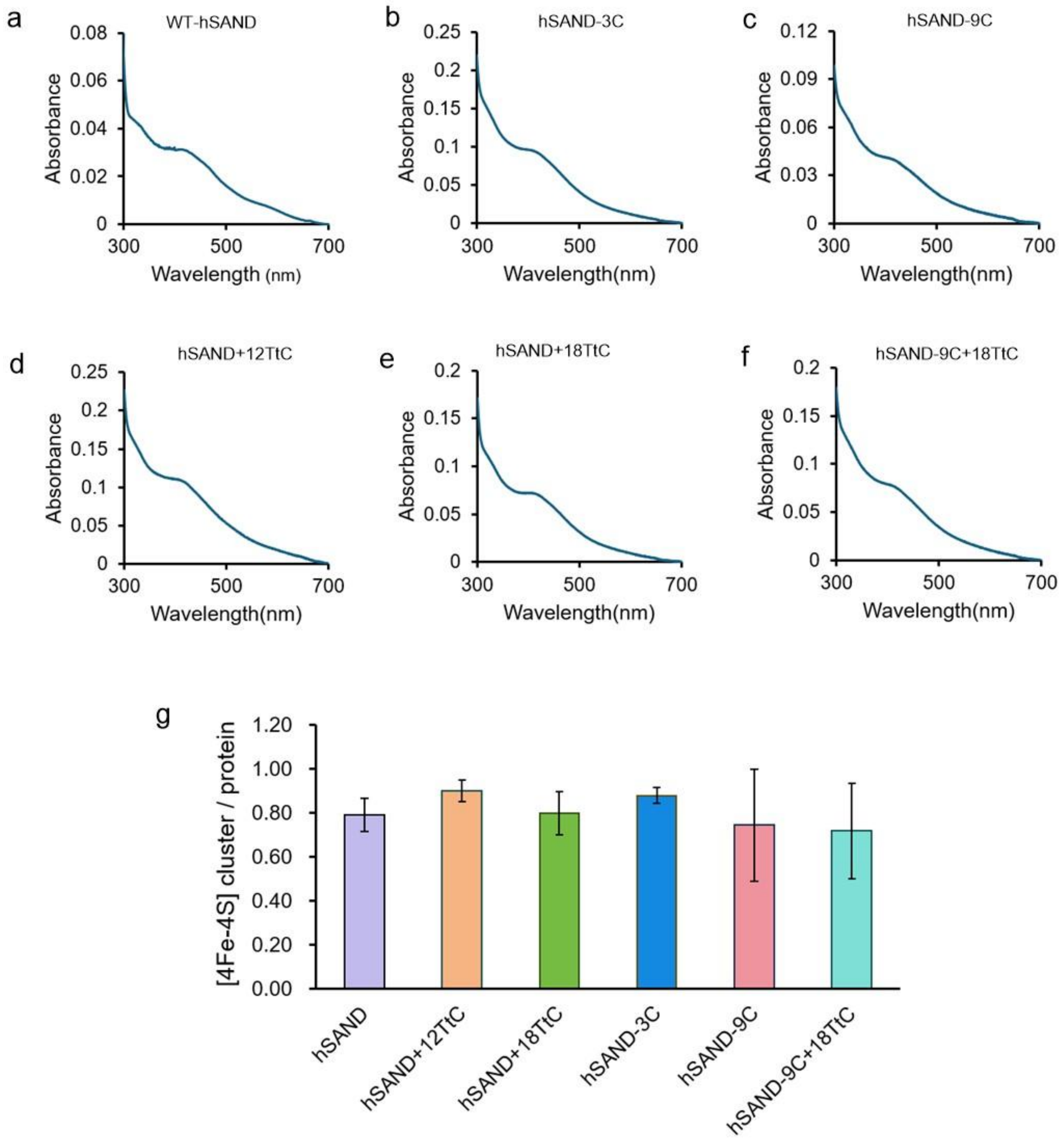


**Figure S7.** Truncating the C-terminus of hSAND or creating chimeric hSAND variants does not affect the [4Fe-4S] cluster. UV-visible absorbance spectrum of the oxidized $[4Fe\text{-}4S]^{1+}$ cluster in (a) WT-hSAND (4.1 μM), (b) hSAND-3C (11 μM), (c) hSAND-9C (7.1 μM), (d) hSAND+12TtC (11.3 μM), (e) hSAND+18TtC (9.9 μM), (f) TtSAND-9C+18TtC (8.5 μM). All measurements were repeated at least two times to confirm reproducibility. The spectra show a peak at around 420 nm characteristic of the oxidized $[4Fe\text{-}4S]^{1+}$ cluster. (g) Analysis of the cluster content of each purified protein. Data are the average of two measurements ± errors.

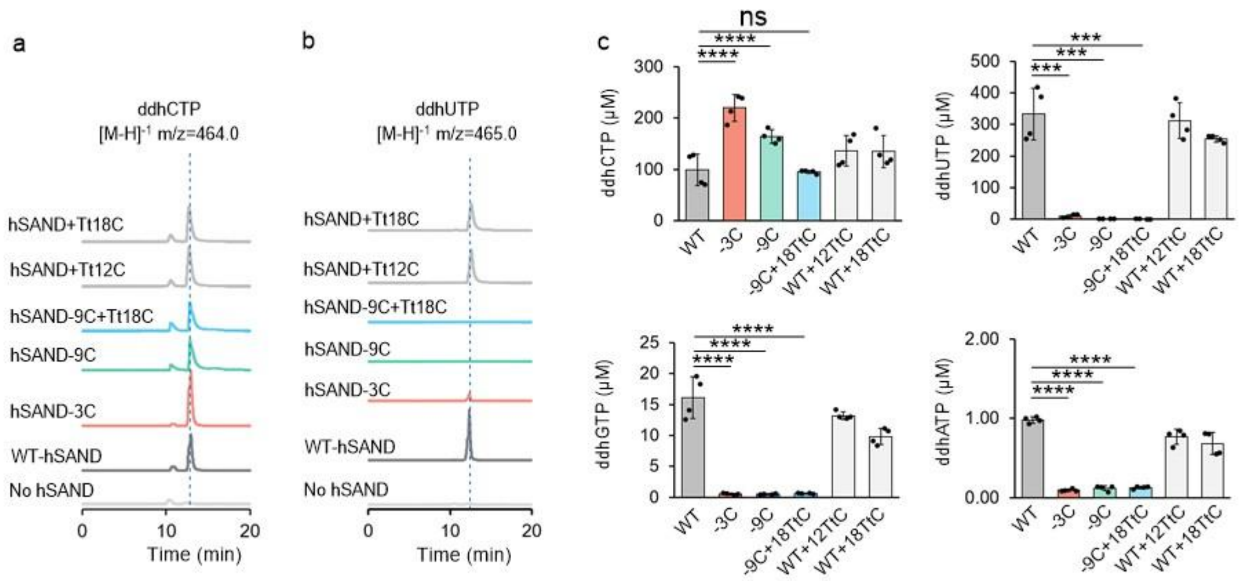


**Figure S8**. The tripeptide LDW in hSAND (viperin) is required for substrate promiscuity. **(a-b)** Total ion chromatograms of (a) ddhCTP and (b) ddhUTP formation. **(c)** Quantification of the amount of ddhCTP, ddhUTP, ddhGTP, and ddhATP. The data are averages of four independent measurements ± standard deviation, obtained from two different protein batches. One-way ANOVA was used to calculate p-values. (***p value < 0.001; ****p value < 0.0001).

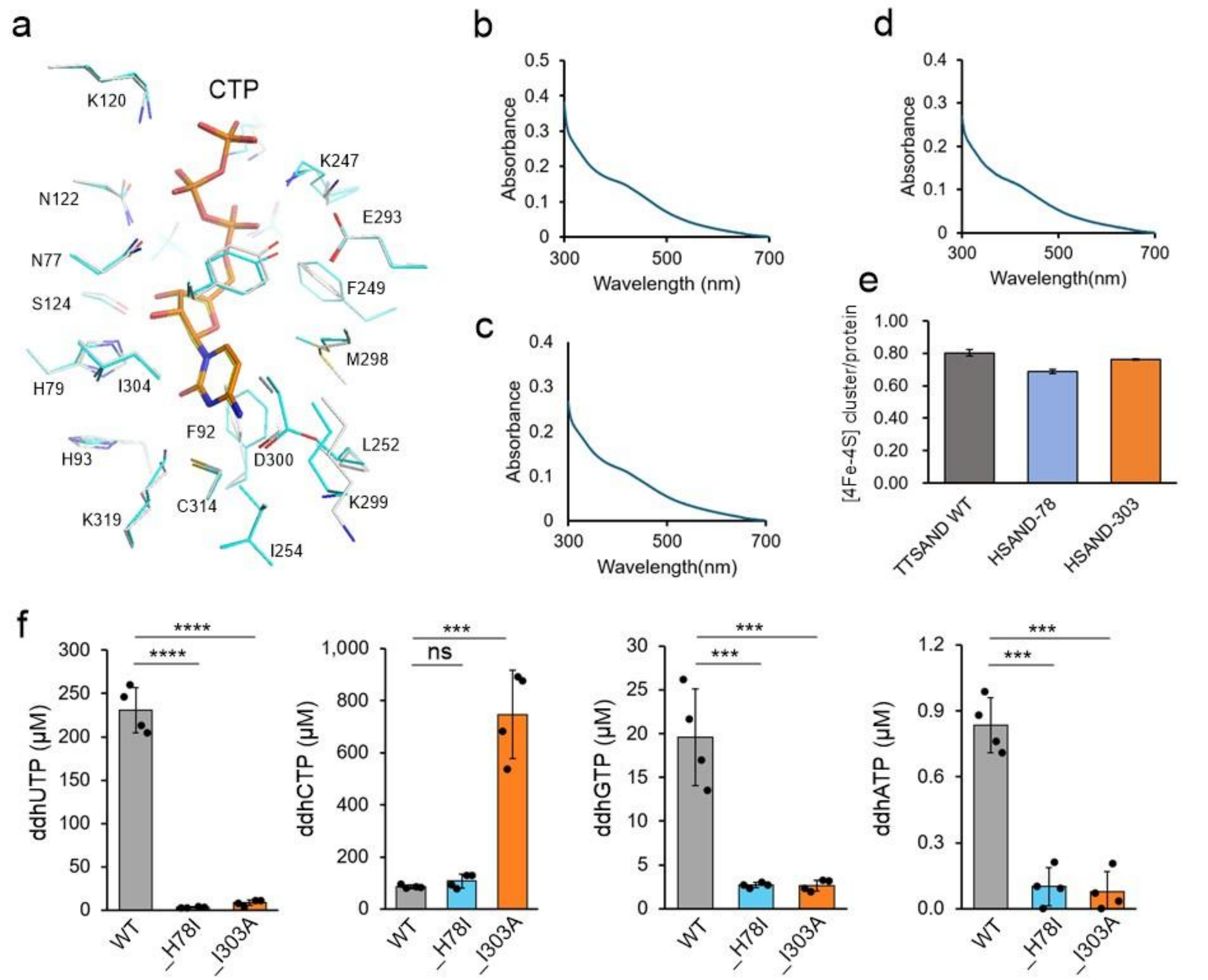


**Figure S9.** hSAND catalytic site is optimized to accept different rNTP substrates. (a) The residues of the substrate-binding site are highly conserved in rat and human SAND (viperin). The numbering is for rat SAND (rSAND). (b-d) UV-visible absorbance spectrum of (b) WT-hSAND (13 µM), (c) hSAND_H78I (8 µM), and (d) hSAND_I303A (9 µM). (e) Quantification of the amount of [4Fe-4S] clusters per protein. Data are averaged from two measurements using two different batches of protein. (f) Quantification of the amount of ddhUTP, ddhCTP, ddhGTP, and ddhATP formed by WT-hSAND, hSAND_H78I, and hSAND_I303A. The data are averages of four independent measurements ± standard deviation, obtained from two different protein batches. One-way ANOVA was used to calculate p-values. (***p value < 0.001; ****p value < 0.0001).

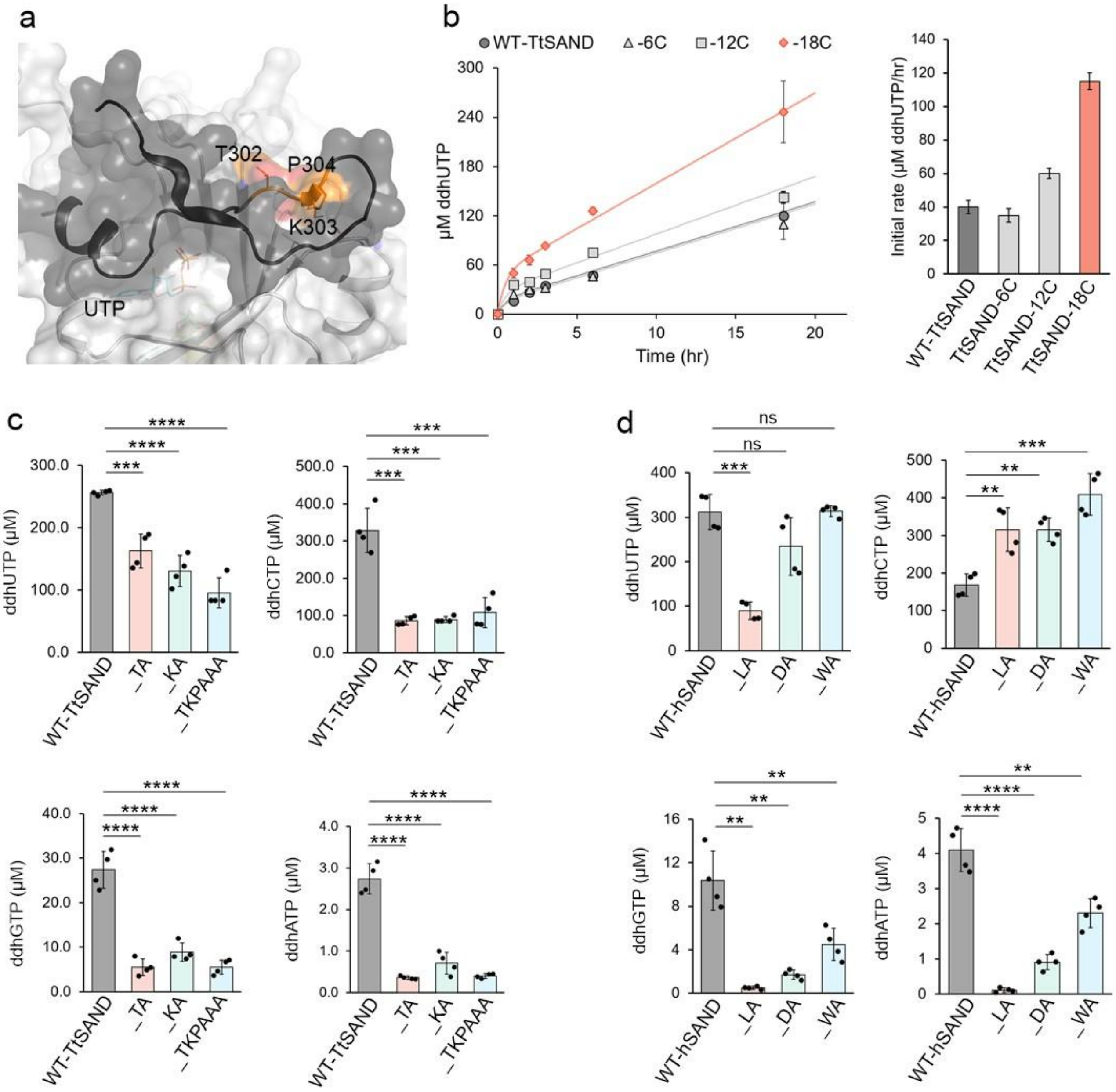


**Figure S10.** The IDR tripeptide acts as a latch and tunes the IDR ensemble. **(a)** Analysis of the modeled TtSAND structures shows that the IDR tripeptide is anchored in a pocket on the surface of the radical-SAM domain. **(b)** Removal of the tripeptide sharply increased the initial rate of TtSAND (110 μM ± 20). **(c-d)** Alanine mutagenesis of the tripeptide affects synthesis of different ddhNTPs. (c) Quantification of the amount of ddhNTP produced by WT-TtSAND and its variants (T302A, K303A, or TKP-AAA) (110 μM ± 20). (d) Quantification of the amount of ddhNTP produced by WT-hSAND and its variants (L359A, D360A, or W361A) (40 μM). One-way ANOVA was used to calculate p-values. (*p value < 0.05; **p value < 0.01; ***p value < 0.001; ****p value < 0.0001).

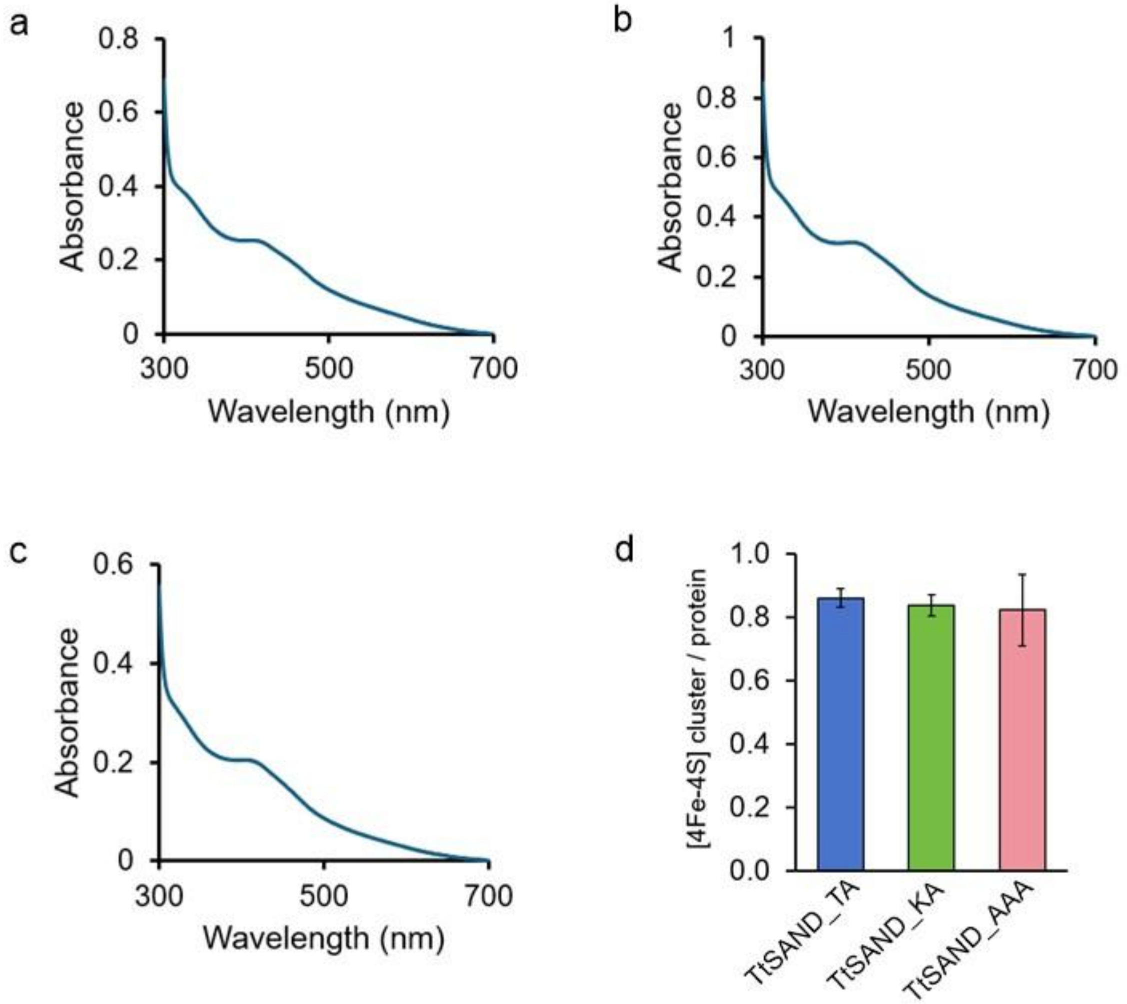


**Figure S11.** Alanine-scanning mutagenesis of the tripeptide TKP does not affect the FeS cluster in TtSAND. (a-c) UV-visible absorbance spectra of (a) TtSAND-TA (T302A) (27.7 µM), (b) TtSAND-KA (K303A) (34 µM), and (c) TtSAND-AAA (TKP-AAA) (24.3 µM) showing the absorbance peak of the oxidized $[4Fe\text{-}4S]^{1+}$ cluster with a peak around 420 nm. (d) Measurement of the amount of [4Fe-4S] cluster per protein. Measurements were repeated twice with different protein batches. Data are the average of two measurements ± errors.

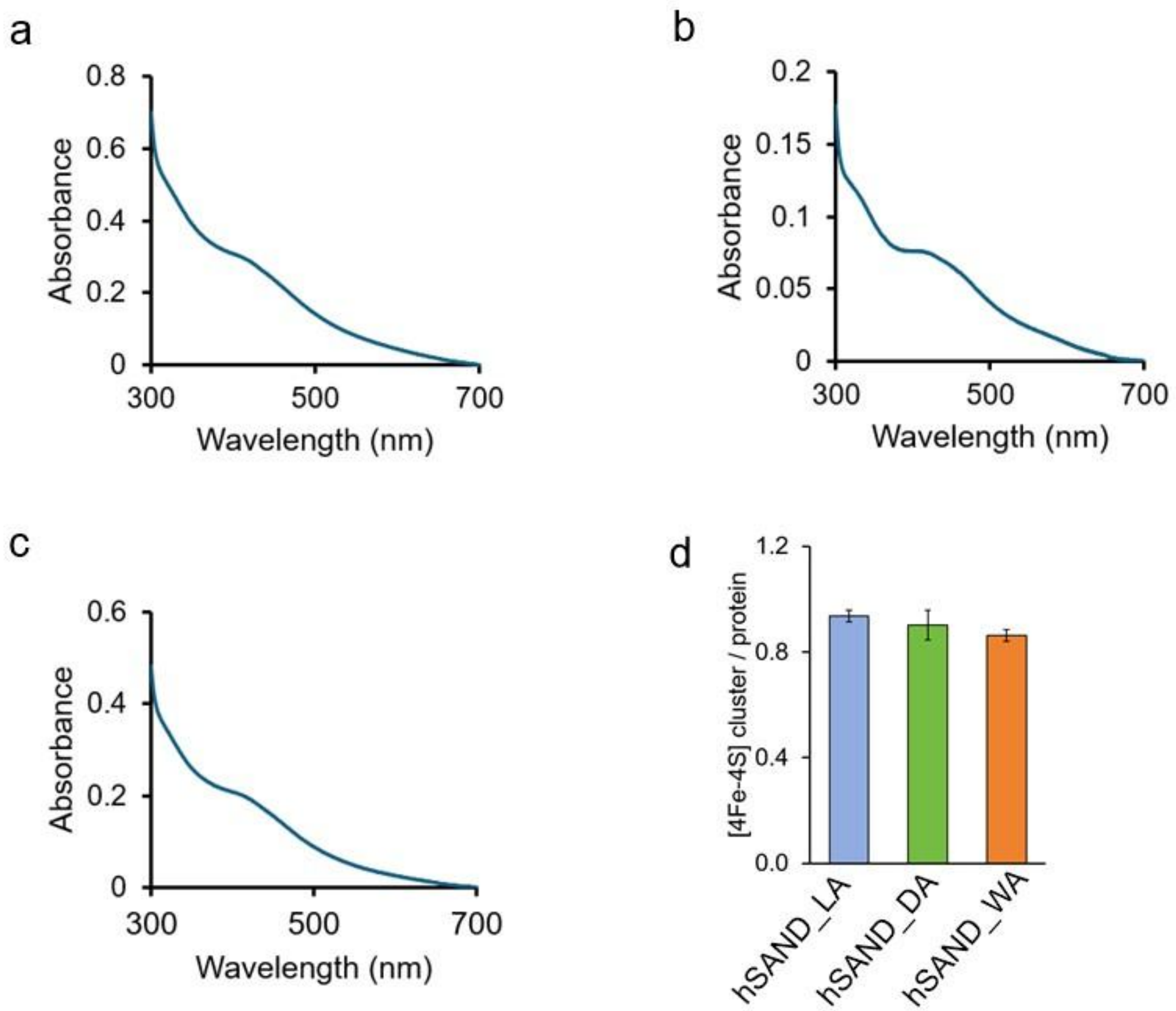


**Figure S12.** Alanine-scanning mutagenesis of the tripeptide LDW does not affect the FeS cluster in hSAND. (a) UV-visible absorbance spectrum of hSAND_LA (L359A) (16.7 μM), (b) hSAND_DA (D360A) (8.8 μM), and (c) hSAND_WA (W361A) (15 μM). The spectra show the absorbance peak (420 nm) of the oxidized $[4Fe\text{-}4S]^{1+}$ cluster. (d) Measurement of the amount of [4Fe-4S] cluster per protein. Measurements were repeated twice with different batches of protein. Data are the average of two measurements ± errors.

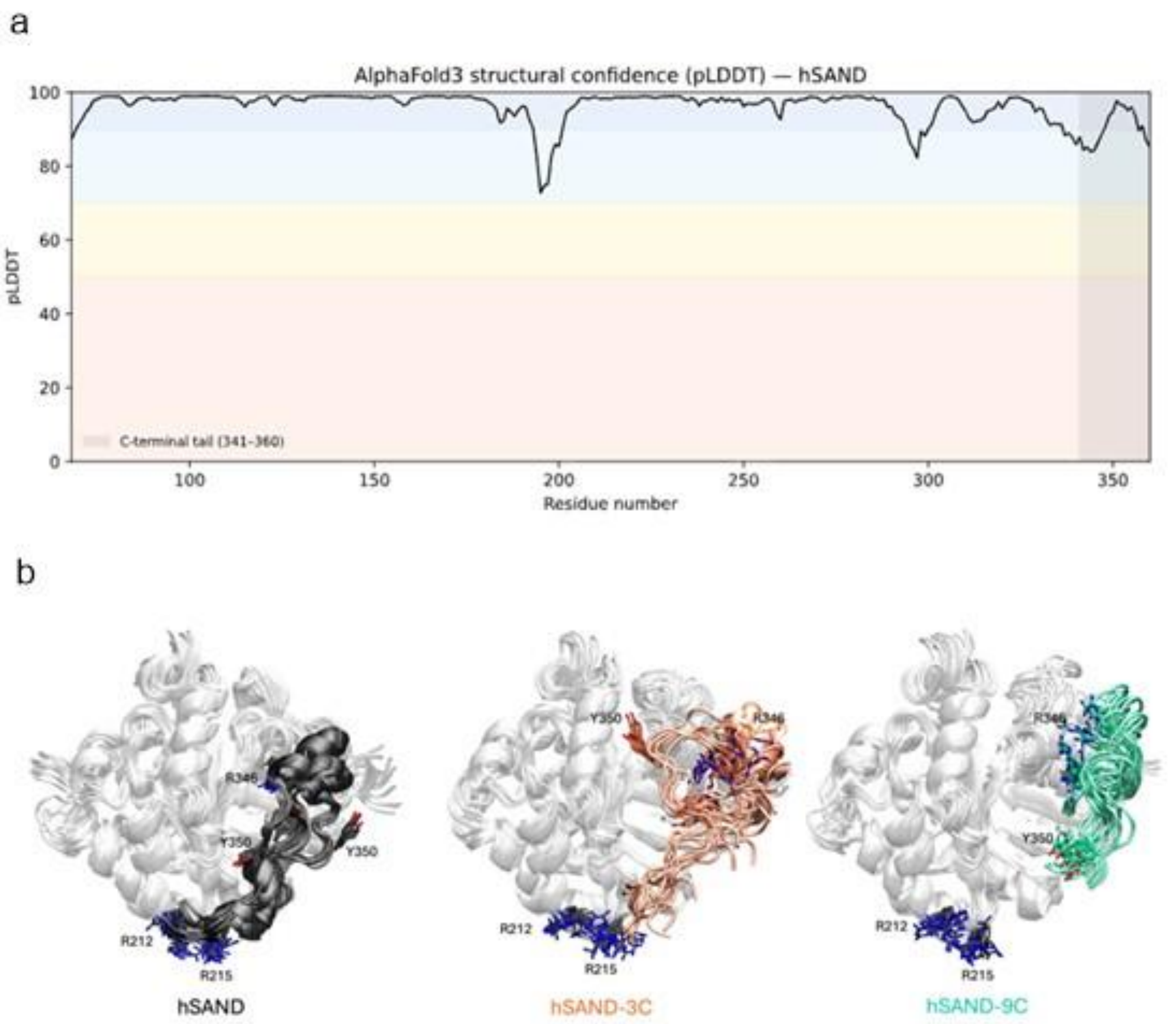


**Figure S13.** MD simulations of IDR ensemble. (a) Per-residue pLDDT values of the top-ranked AlphaFold3 model (model_0) for hSAND (residues 68–361). pLDDT scores were extracted from the Cα B-factor field, yielding a mean confidence score of 96.11, indicative of a highly reliable structural model. C-terminal tail is shaded in gray. Residue numbering was remapped from the AlphaFold3 sequence numbering (1–293) to the native hSAND numbering (68–361) to maintain consistency throughout the study. (b) Conformational ensemble of the apo WT hSAND, hSAND-3C, and hSAND-9C variants from three independent MD simulations per variant. The protein core is depicted as a light gray cartoon, while the C-terminal extension (residues 340–361) is shown in black (WT), orange (hSAND-3C), and green (hSAND-9C). The putative anchoring residues Arg212 and Arg215, along with the phosphate-coordinating residues Arg346 and Tyr350 in the C-terminal extension, are shown in licorice representation.

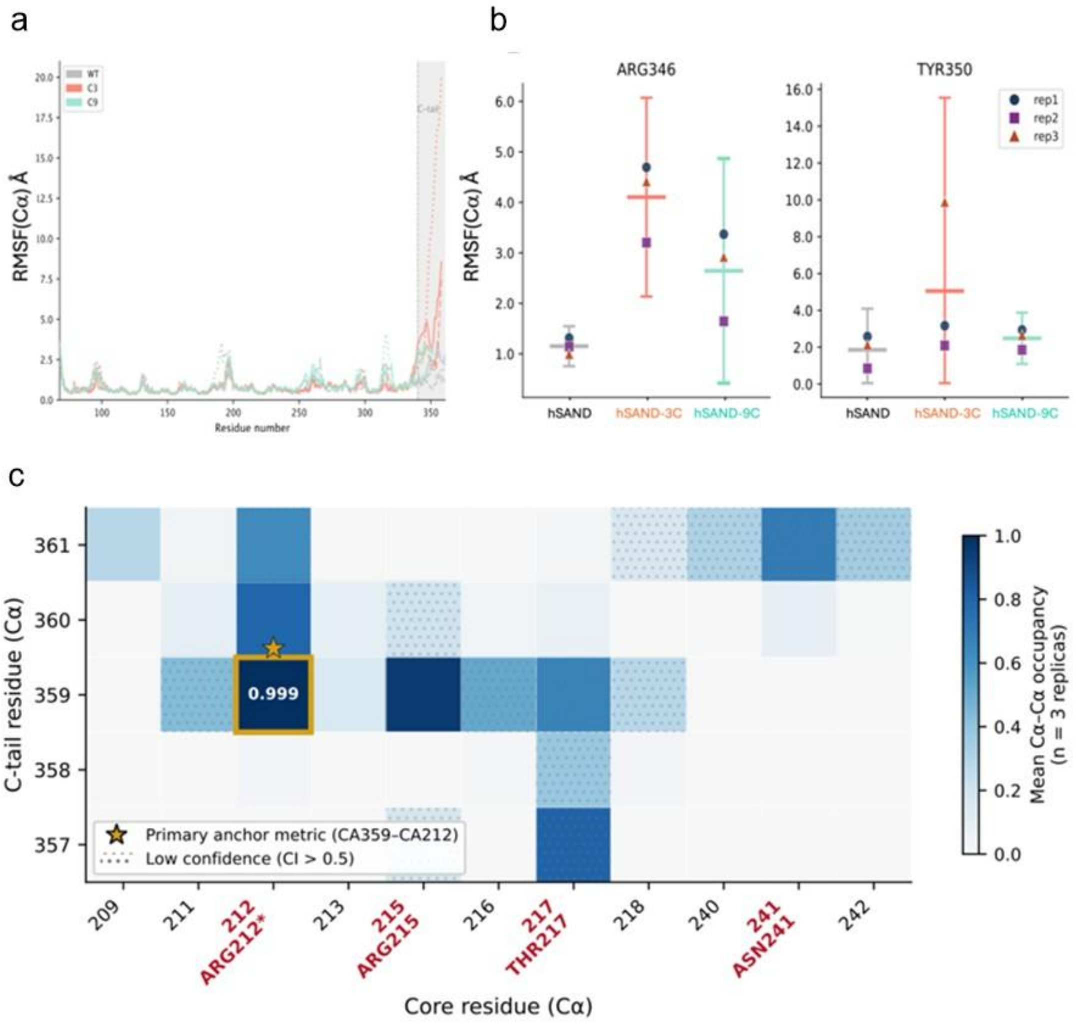


**Figure S14.** Distance measurements. (a) Per-residue Cα root-mean-square fluctuations (Cα-RMSF) calculated from three independent 200 ns MD simulations of WT hSAND (grey), hSAND-3C (orange), and hSAND-9C (green). The truncated C-terminal regions are shaded in grey. The highly similar RMSF profiles of the protein core indicate that C-terminal truncation has only a minor effect on the overall conformational dynamics of the folded domain. (b) Cα-RMSF of the phosphate-coordinating C-tail residues Arg346 and Tyr350. In WT hSAND, the C-tail is stabilised by anchoring interactions, resulting in low flexibility of Arg346 (1.15 ± 0.40 Å). Loss of the C-tail anchor in hSAND-3C markedly increases the mobility of Arg346 (4.10 ± 1.97 Å). In hSAND-9C, Arg346 also exhibits increased flexibility (2.64 ± 2.22 Å), although with greater inter-replica variability, consistent with partial sampling of ordered C-tail conformations despite geometric exclusion from the anchoring pocket. (c) Persistent Cα–Cα contacts identify Arg212 and Arg215 as the primary anchoring residues for the hSAND C-tail. Contact occupancy maps between the C-terminal residues 357–361 and core residues 209–242 of WT hSAND, calculated from three independent MD simulations. The highest contact occupancies (0.999 ± 0.004) are observed between Arg212 and the three residues of the conserved LDW motif, as well as between Arg215 and Leu359 (0.956 ± 0.037). Based on these highly persistent interactions, Arg212 and Arg215 are defined as the principal anchoring residues for C-tail binding throughout this work. Contacts were defined using a Cα–Cα distance cutoff of 8 Å, and occupancy is reported as the fraction of simulation frames in which a given contact was present.

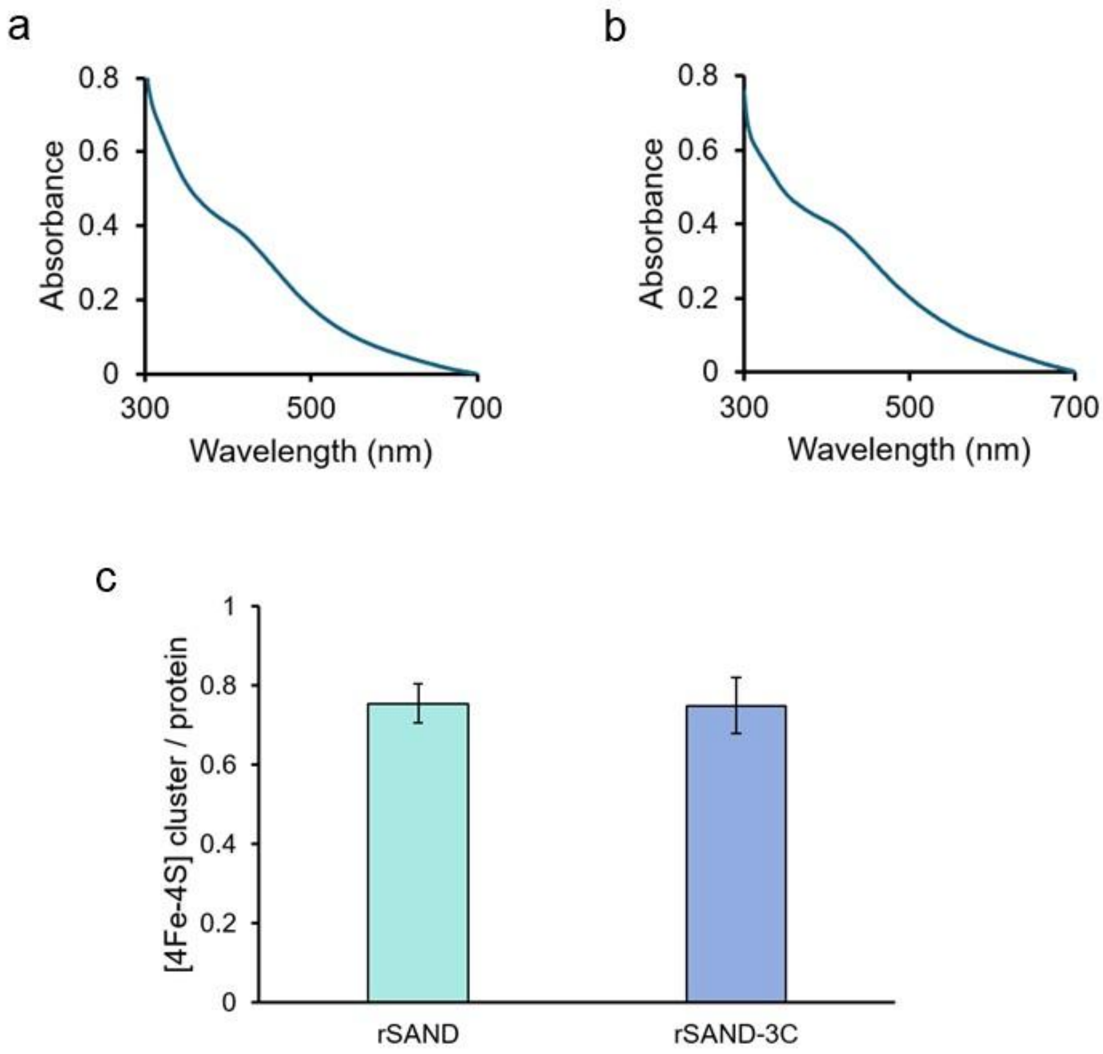


**Figure S15.** Removal of the C-terminal tripeptide in rSAND does not affect the FeS cluster. (a) UV-visible absorbance spectra of wild-type rSAND (WT-rSAND) and (b) mSNAD-3C lacking the tripeptide LDW. Protein concentrations were 18.0 ± 1.5 µM. The spectra show a peak at around 420 nm characteristic of the oxidized $[4Fe\text{-}4S]^{1+}$ cluster. (c) The number of [4Fe-4S] clusters per protein is not affected by the removal of the LDW tripeptide. Data are the average of two measurements using different batches of proteins ± errors

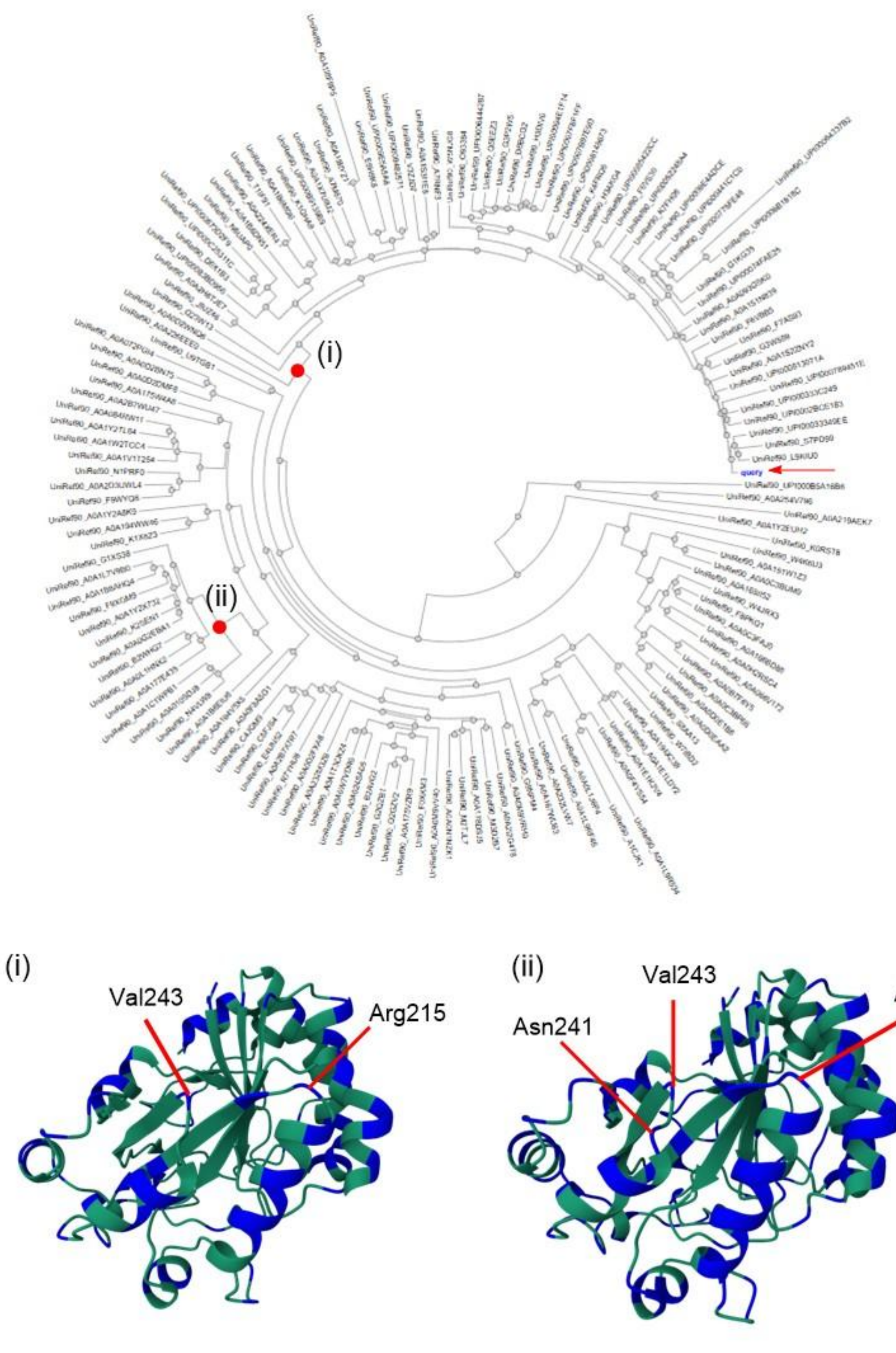


**Figure S16.** Ancestral sequence reconstruction (ASR) confirms that the folded protein context adjacent to the tripeptide latch is a hot spot for substitutions. The phylogenetic tree was generated from ancestral sequence reconstruction using FireProt. The query, hSAND, is marked with a red arrow. Two nodes were selected (i) and (ii) as examples. The representative amino acid substitutions for each node were highlighted in the wild-type hSAND structure in blue. The residues of the folded domain adjacent to the tripeptide are numbered. The C-terminal IDR is not shown for clarity.

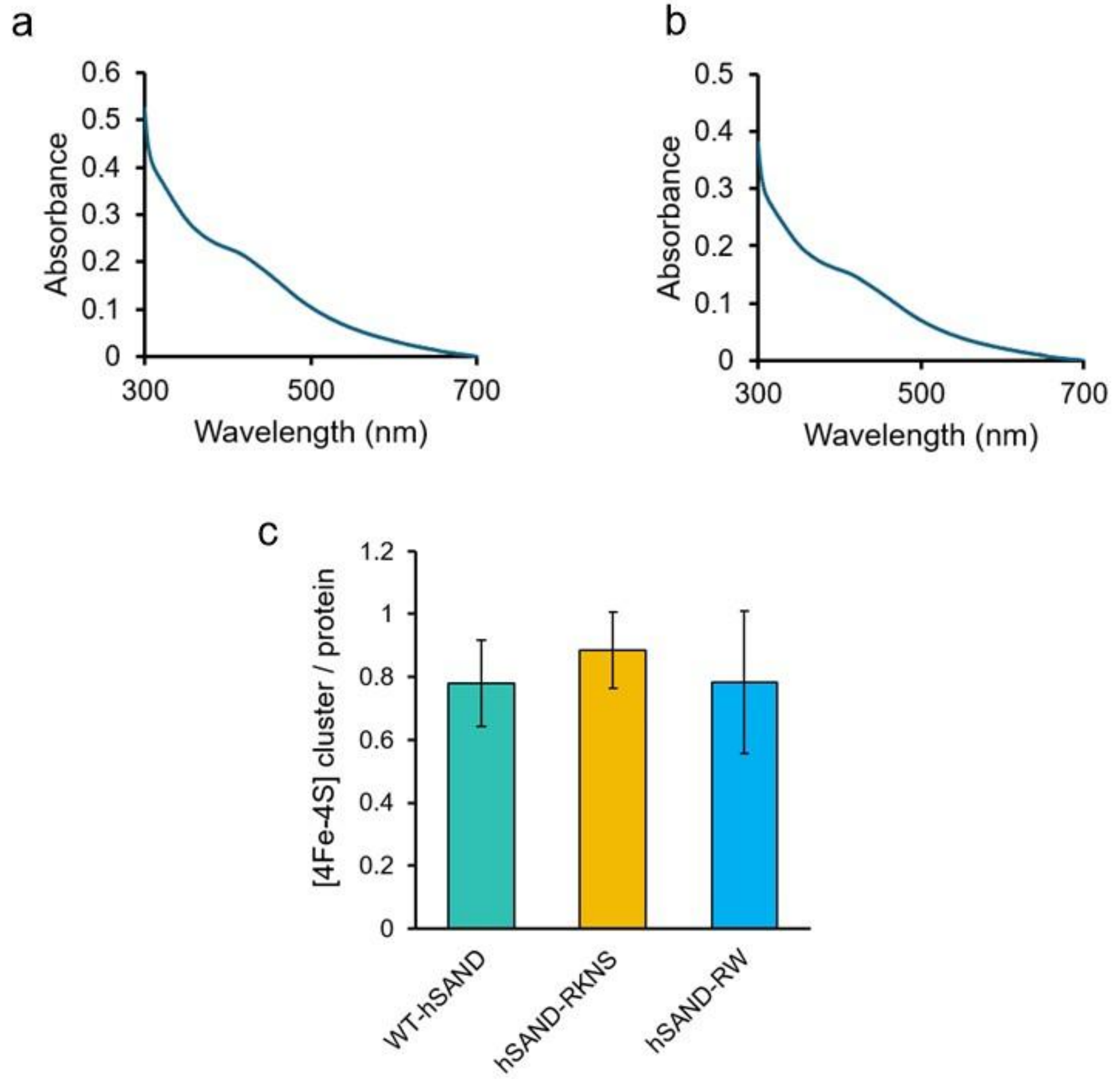


**Figure S17.** Mutation of the folded radical-SAM domain of hSAND adjacent to the LDW tripeptide does not affect the FeS cluster. (a) UV-visible absorbance spectra of hSAND-RKNS, encompassing Arg215Lys and Asn241Ser substitutions, and (b) that of hSAND-RW, encompassing Arg215Trp substitution. The spectra show an absorbance peak at around 420 nm, characteristic of an oxidized $[4Fe\text{-}4S]^{1+}$ cluster. The protein concentration was 13.5 ± 1.0 µM. (c) Quantification of the number of clusters per protein shows no significant change. Data are the average of two measurements from different protein batches, ± error.

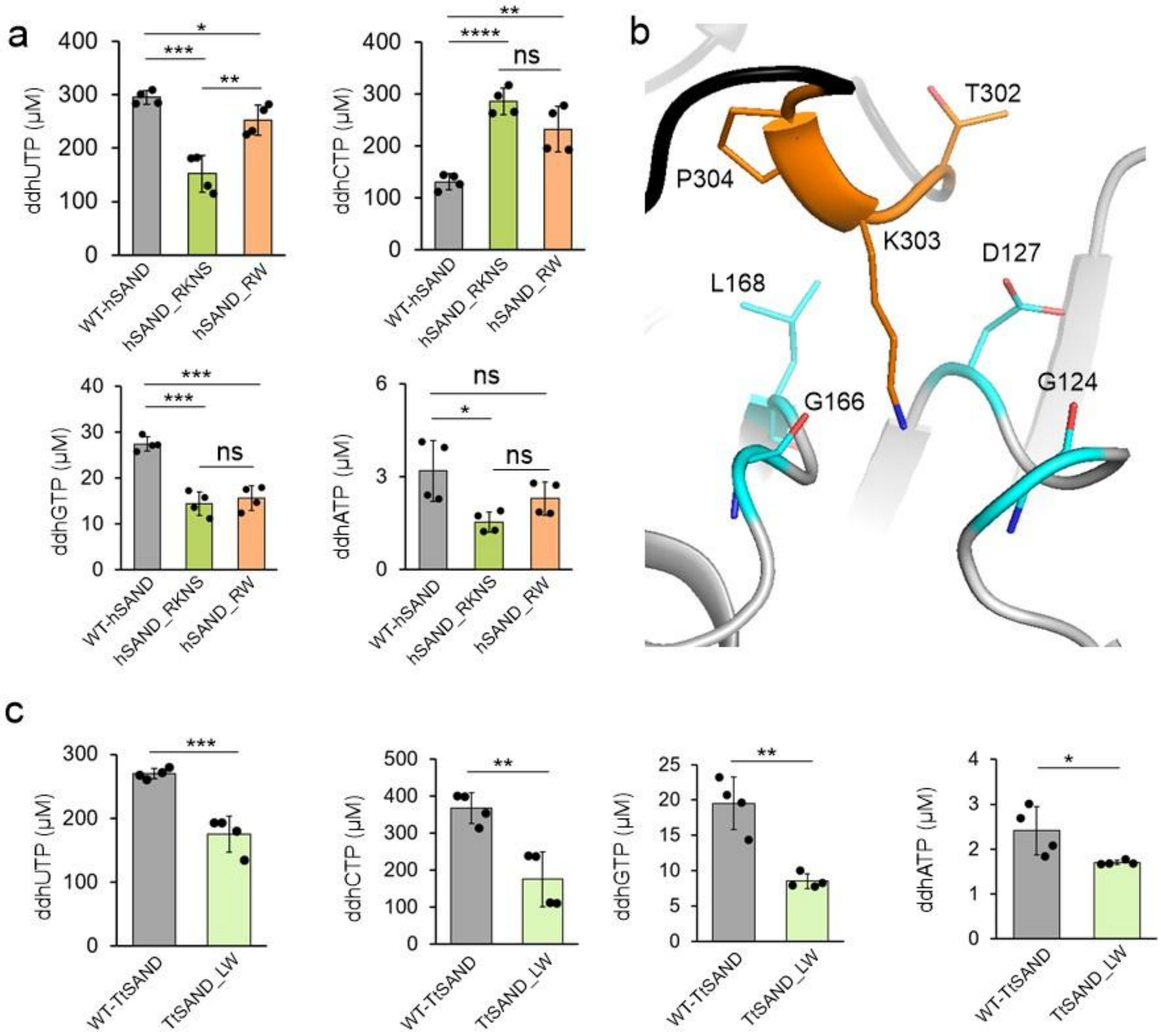


**Figure S18.** The IDR tripeptide context modulates enzyme output. (a) Modification of the tripeptide context in hSAND changes the activity profile. Quantification of the amount of ddhNTP formed by WT-hSAND, hSAND_RKNS (R215K and N241S), and hSAND_RW (R215W) (40 μM). (b-c) Modification of the tripeptide context in TtSAND changes the activity profile. (b) Predicted structure of TtSAND showing the tripeptide latch and its context. (c) Quantification of the amount of ddhNTP formed by WT-TtSAND and TtSAND_LW (L168W) (110 μM). One-way ANOVA was used to calculate p-values. (*p value $< 0.05$; **p value $< 0.01$; ***p value $< 0.001$; ****p value $< 0.0001$).

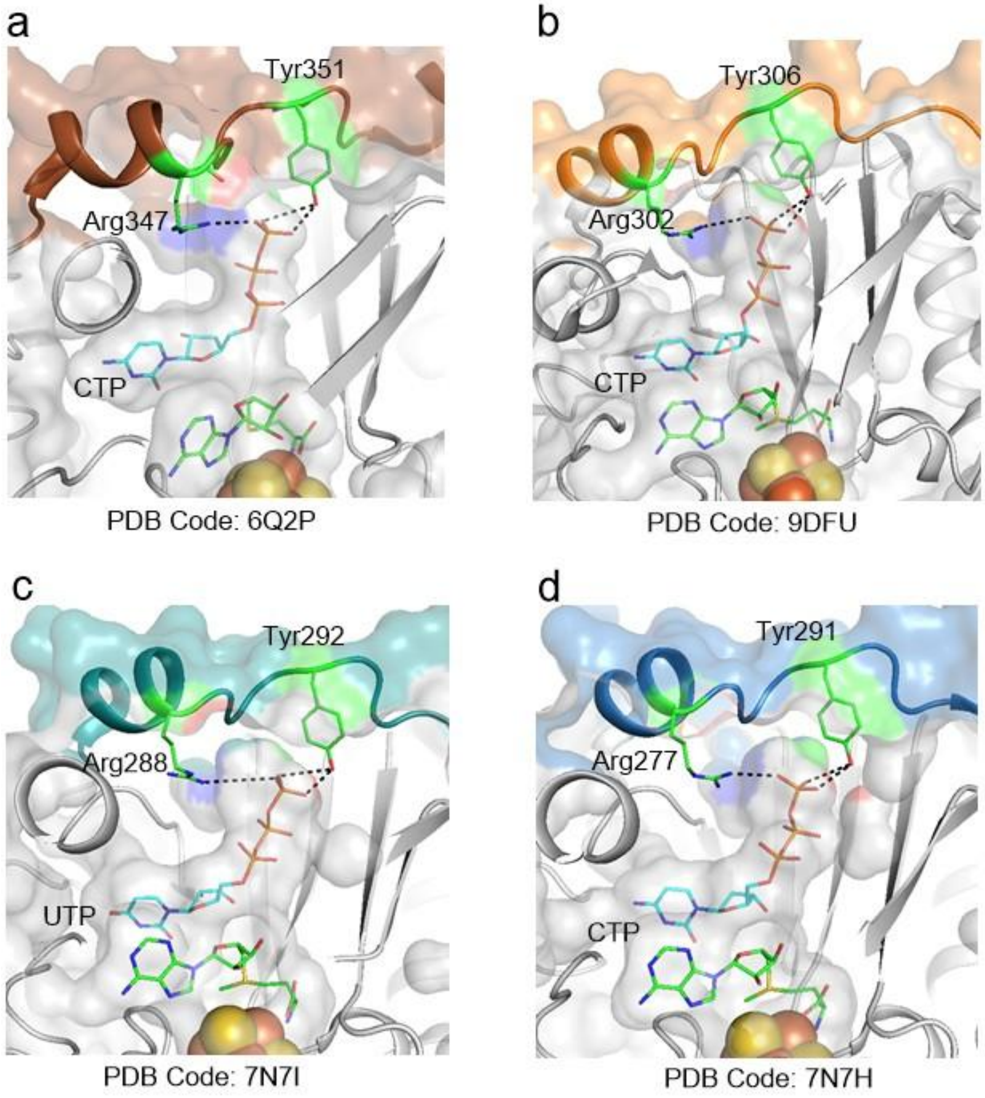


**Figure S19.** Highly conserved arginine and tyrosine residues of the IDR form hydrogen bonds with the terminal phosphate group of an NTP. (a) rSAND (PDB Code: 6Q2P), (b) (PDB Code: 9DFU), (c) (PDB Code: 7N7I), (d) (PDB Code: 7N7H).

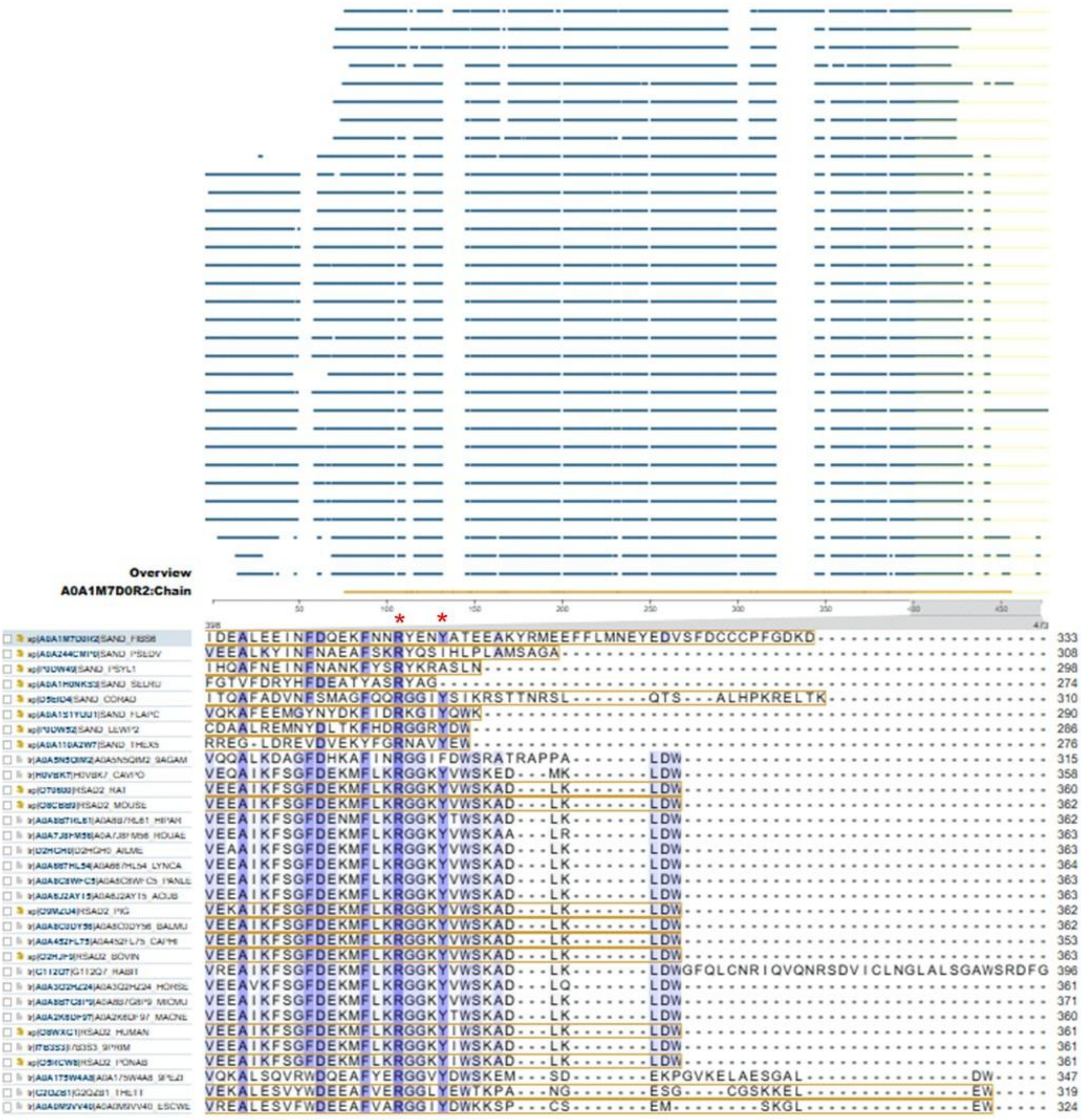

**Figure S20.** Highly conserved arginine and tyrosine residues of IDR form hydrogen bonds with the terminal phosphate group in an NTP substrate. These residues are highlighted by a red star.

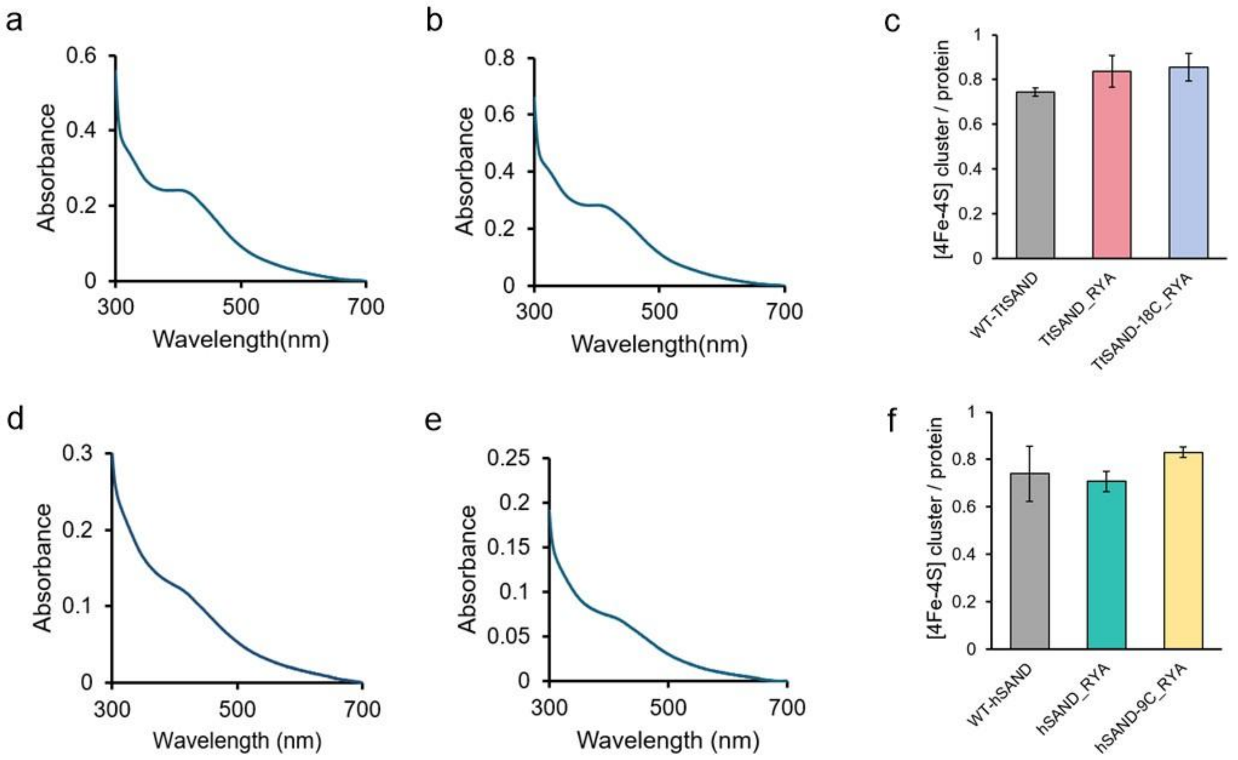


**Figure S21.** Mutation of conserved arginine and tyrosine in the IDR does not affect the [4Fe-4S] cluster. (a-b) UV-visible absorbance spectra of (a) TtSAND-RYA (25 μM) and (b) TtSAND-18C-RYA (27 μM). (c) Quantification of the [4Fe-4S] cluster per protein. Data are the average of two independent biological replicates ± errors. (d-e) UV-visible absorbance spectra of (d) hSAND-RYA (10 μM) and (e) hSAND-9C-RYA (9 μM). (f) Quantification of the [4Fe-4S] cluster per protein. Data are the average of two independent biological replicates ± errors.

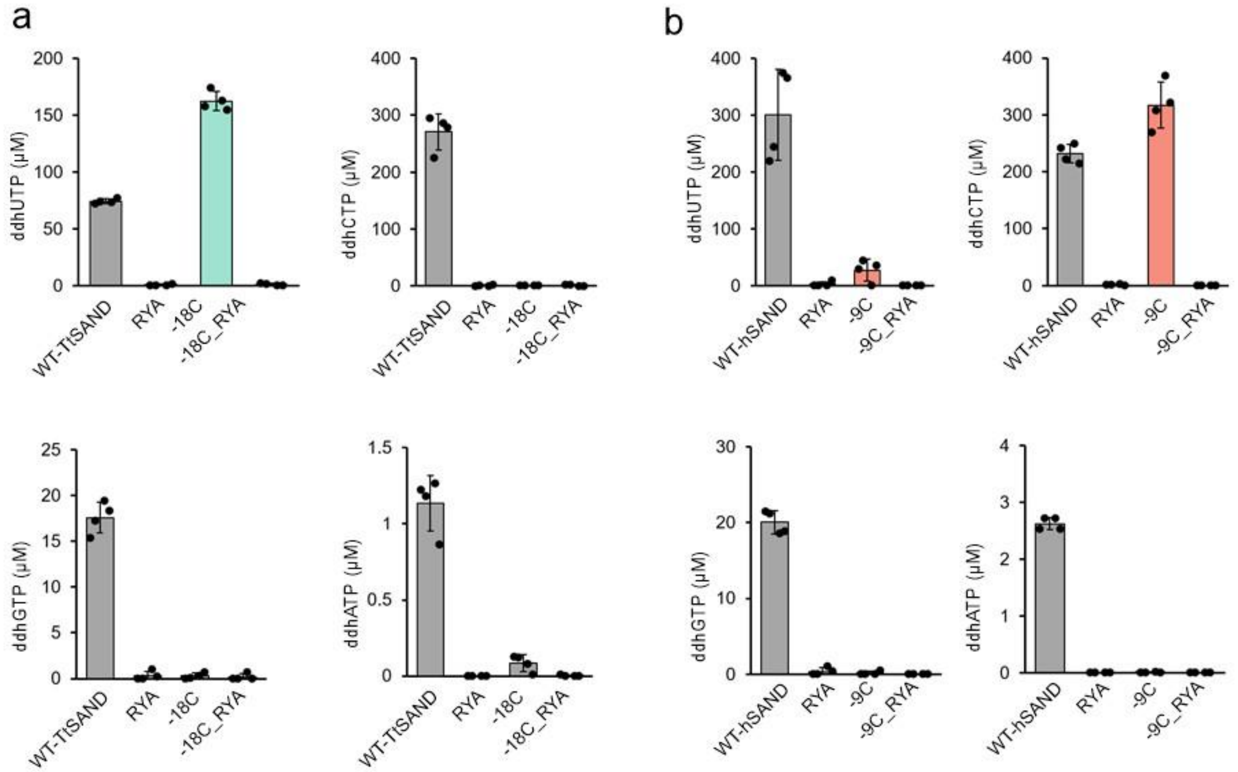


**Figure S22.** The IDR clamp sequence is vital for activity. (a-b) Substitution of arginine and tyrosine residues with alanine (designated as RYA mutant) fully abolishes the formation of all ddhNTPs. (a) Quantification of ddhNTP formation by (c) hSAND and its RYA variants (R346A and Y350A) (40 μM) and (b) WT-TtSAND RYA variants (R295A and Y299A) (110 μM). Data are the average of two measurements ± standard deviation. One-way ANOVA was used to calculate p-values. (all p values: ****p value < 0.0001).

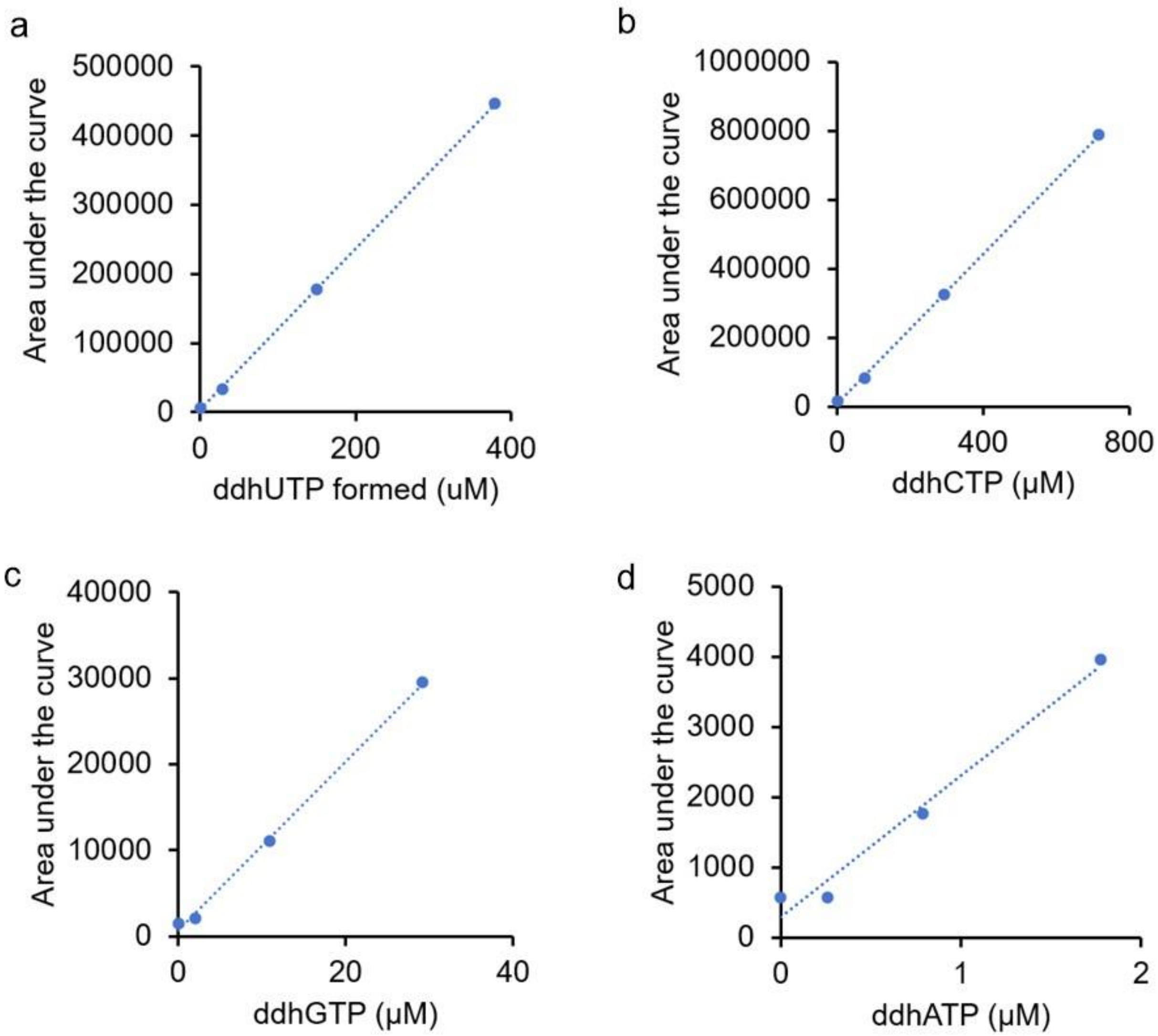


**Figure S23.** Standard curves for (a) ddhUTP, (b) ddhCTP, (c) ddhGTP, and (d) ddhATP. The area under the total ion chromatograms for ddhUTP, ddhCTP, ddhGTP, or ddhATP is plotted as a function of ddhNTP concentration.

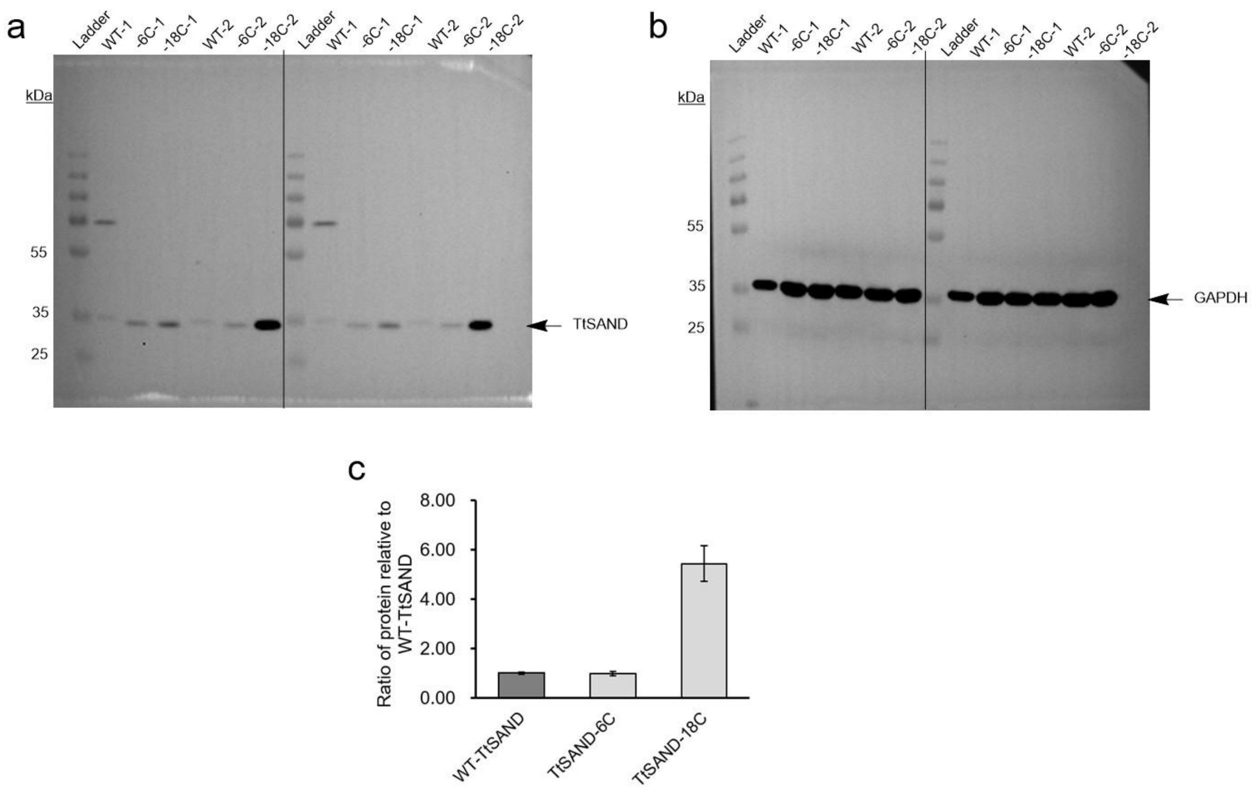


**Figure S24.** Analysis of WT-TtSAND and its C-terminal-truncated variant expression for the VITAS assay. (A-B) Western blot analysis of the expression of (a) TtSAND and (b) GAPDH loading control. An HRP-conjugated His-tag antibody was used to detect ectopic TtSAND expression in *E. coli* cells. (c) Quantification of the amount of protein expressed relative to GAPDH control. The quantification was used to normalize the data from the VITAS assay, SI Methods.

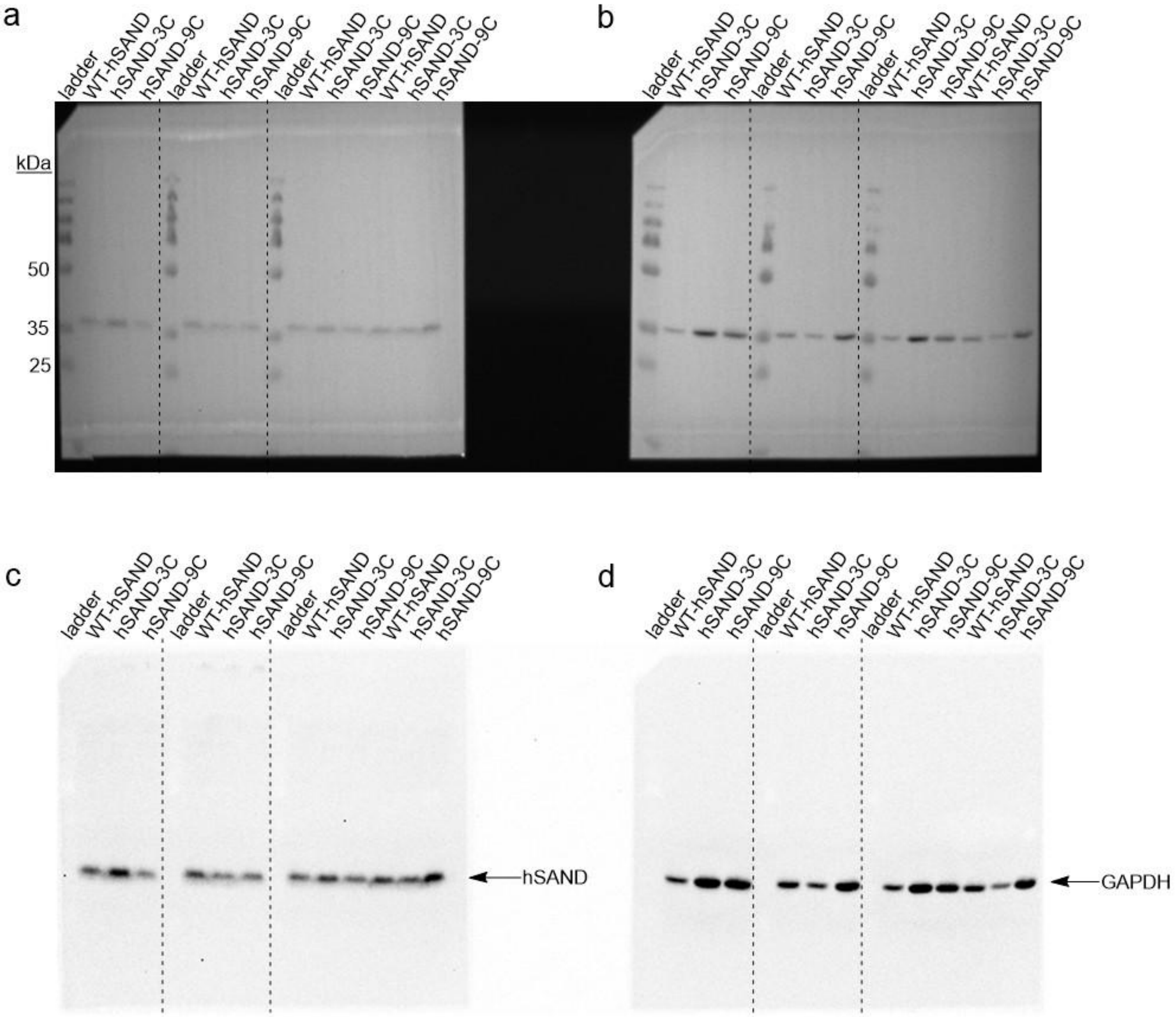


**Figure S25.** Western blot analysis of the expression of WT-hSAND, hSAND-3C, and hSAND-9C for the VITAS assay. (a-b) Overlay of the chemiluminescence and the image of the blot showing the ladder. The images of (a) hSAND detection using His-tag antibody and (b) GAPDH detection using HRP-conjugated GAPDH antibody are presented. (c-d) The chemiluminescence images of (a) and (b). Two different batches of hSAND were used. Each expression was repeated twice. Protein-level quantification was used to correct the data obtained from the VITAS assay.

**Table S1.** HR LC-MS analysis of ddhCTP ($[M\text{-}H]^{-1}$ m/z=463.9678) and ddhUTP ($[M\text{-}H]^{-1}$ m/z=464.9513) formation in HEK293 cells co-expressing hSAND and its electron-transfer partner protein CYB5R3, or in cells expressing hSAND alone, or CYB5R3 alone.

| **hSAND + CYB5R3** | | **Sample 1** | **Sample 2** | **Sample 3** |
|---|---|---|---|---|
| **Retention time (min)** | **ddhCTP** | 7.34 | 7.39 | 7.36 |
| | **ddhUTP** | 7.21 | 7.26 | 7.34 |
| **Area** | **ddhCTP** | 399994 | 316194 | 203294 |
| | **ddhUTP** | 113837 | 86934 | 126668 |
| **Score** | **ddhCTP** | 73.91 | 78.11 | 83.72 |
| | **ddhUTP** | 82.27 | 64.72 | 92.61 |
| **hSAND alone** | | | | |
| **Retention time (min)** | **ddhCTP** | 7.24 | 7.19 | 7.44 |
| | **ddhUTP** | 7.17 | 7.19 | 7.15 |
| **Area** | **ddhCTP** | 10805 | 62862 | 21987 |
| | **ddhUTP** | 32595 | 30132 | 37983 |
| **Score** | **ddhCTP** | 47.1 | 65.4 | 80.22 |
| | **ddhUTP** | 52.94 | 56.63 | 60.19 |
| **CYB5R3 alone** | | | | |
| **Retention time (min)** | **ddhCTP** | 7.3 | 7.40 | 7.38 |
| | **ddhUTP** | 7.37 | 7.31 | 7.1 |
| **Area** | **ddhCTP** | 215461 | 283014 | 156682 |
| | **ddhUTP** | 58770 | 49664 | 52136 |
| **Score** | **ddhCTP** | 80.6 | 78.43 | 76.6 |
| | **ddhUTP** | 83.1 | 78.25 | 76.8 |

**Table S2.** Kinetic parameters obtained from fitting a slow-binding inhibition model to the progress curve of ddhNTP formation (ddhCTP for hSAND or ddhUTP for TtSAND). The estimated $K_{obs}$ for TtSAND and hSAND are within experimental error, similar to those reported for rSAND[14,15] and other radical-SAM enzymes.[16]

| Enzyme | $v_i$ ($\mu M$ ddhNTP/hr) | $v_s$ ($\mu M$ ddhNTP/hr) | $K_{obs}$ ($hr^{-1}$) |
|---|---|---|---|
| WT-TtSAND | 40 ± 6 | 6 ± 2 | 2.0 ± 0.5 |
| TtSAND-6C | 35 ± 5 | 6 ± 1 | 2.0 ± 0.4 |
| TtSAND-12C | 60 ± 9 | 7 ± 1.5 | 1.9 ± 0.6 |
| TtSAND-18C | 115 ± 15 | 9 ± 2 | 2.1 ± 0.5 |
| WT-hSAND | 35 ± 5 | 3.5 ± 1.1 | 1.8 ± 0.5 |
| hSAND-3C | 75 ± 8 | 8.0 ± 2 | 2.0 ± 0.8 |
| hSAND-9C | 65 ± 5 | 7.0 ± 1 | 1.9 ± 0.5 |
